# Evidence of selection, adaptation and untapped diversity in Vietnamese rice landraces

**DOI:** 10.1101/2020.07.07.191981

**Authors:** Janet Higgins, Bruno Santos, Tran Dang Khanh, Khuat Huu Trung, Tran Duy Duong, Nguyen Thi Phuong Doai, Nguyen Truong Khoa, Dang Thi Thanh Ha, Nguyen Thuy Diep, Kieu Thi Dung, Cong Nguyen Phi, Tran Thi Thuy, Nguyen Thanh Tuan, Hoang Dung Tran, Nguyen Thanh Trung, Hoang Thi Giang, Ta Kim Nhung, Cuong Duy Tran, Son Vi Lang, La Tuan Nghia, Nguyen Van Giang, Tran Dang Xuan, Anthony Hall, Sarah Dyer, Le Huy Ham, Mario Caccamo, Jose De Vega

**Affiliations:** Earlham Institute, Norwich Research Park, Norwich, NR4 7UZ, UK; NIAB, 93 Lawrence Weaver Road, Cambridge, CB3 0LE, UK; Agriculture Genetics Institute (AGI), Hanoi, Vietnam; Vietnam National University of Agriculture, Hanoi, 131000, Vietnam; Graduate School of Advanced Science and Engineering, Hiroshima University, Hiroshima, 739-8529, Japan; Faculty of Biotechnology, Nguyen Tat Thanh University, Ho Chi Minh, 72820, Vietnam; Institute of Research and Development, Duy Tan University, Da Nang, 550000, Vietnam; Plant Resource Center, An Khanh, Hoai Duc, Hanoi, 152900, Vietnam; Faculty of Pharmacy, Duy Tan University, Da Nang, 550000, Vietnam

**Keywords:** Rice, breeding, adaptation, QTL, genetic diversity, GWAS, landraces

## Abstract

Vietnam possesses a vast diversity of rice landraces due to its geographical situation, latitudinal range, and a variety of ecosystems. This genetic diversity constitutes a highly valuable resource at a time when the highest rice production areas in the low-lying Mekong and Red River Deltas are enduring increasing threats from climate changes, particularly in rainfall and temperature patterns.

We analysed 672 Vietnamese rice genomes, 616 newly sequenced, that encompass the range of rice varieties grown in the diverse ecosystems found throughout Vietnam. We described four Japonica and five Indica subpopulations within Vietnam likely adapted to the region of origin. We compared the population structure and genetic diversity of these Vietnamese rice genomes to the 3,000 genomes of Asian cultivated rice. The named Indica-5 (I5) subpopulation was expanded in Vietnam and contained lowland Indica accessions, which had with very low shared ancestry with accessions from any other subpopulation and were previously overlooked as admixtures. We scored phenotypic measurements for nineteen traits and identified 453 unique genotype-phenotype significant associations comprising twenty-one QTLs (quantitative trait loci). The strongest associations were observed for grain size traits, while weaker associations were observed for a range of characteristics, including panicle length, heading date and leaf width. We identified genomic regions selected in both Indica and Japonica subtypes during the breeding of these subpopulations within Vietnam and discuss in detail fifty-two selected regions in I5, which constitute an untapped resource of cultivated rice diversity.

Our results highlight traits and their associated genomic regions, which were identified by fine phenotyping and data integration. These are a potential source of novel loci and alleles to breed a new generation of sustainable and resilient rice.

## Background

Rice production in Vietnam is of great value for export and providing daily food for more than 96 million people. However, agricultural production, especially rice cultivation, is inherently vulnerable to climate variability across all regions in Vietnam. Based on the records of monthly precipitation and temperature from 1975 to 2014 [1], the areas of highest crop production in the low lying Mekong and Red River Deltas are particularly vulnerable to the increasing threat from climate change. In 2017, the total planted area of rice in Vietnam was 7.7 million hectares. This includes 4.2 million hectares in the Mekong River Delta and 1.1 million hectares in the Red River Delta [2]. These are also the areas where most of the population of the county is concentrated. In the Mekong River Delta, the damaging effects of salinisation and drought to rice production have increasingly manifested themselves in recent years [3–6].

Vietnam possesses a vast diversity of native and traditional rice varieties due to its geographical situation, latitudinal range and diversity of ecosystems [7]. This diversity constitutes a largely untapped and highly valuable genetic resource for local and international breeding programs. Vietnamese landraces are disappearing as farmers switch to modern elite varieties. To limit this erosion of genetic resources, several rounds of collection of landraces, particularly from the northern upland areas, have been undertaken since 1987. Thousands of rice accessions have been deposited in the Vietnamese National Genebank at the Plant Resources Center (PRC, Hanoi, Vietnam), together with passport information detailing their traditional name and province of origin. One hundred and eighty-two traditional Vietnamese accessions were selected for a genotype by sequencing (GBS) study in 2014 [8]. This study yielded 25,971 single nucleotide polymorphisms (SNPs) and was used to describe four Japonica and six Indica subpopulations. These subpopulations were classified by region, ecosystem and grain-type using passport information (province and ecosystem) and phenotyping. This dataset had subsequently been used for genome-wide phenotype-genotype association studies (GWAS) relating to root development [9], panicle architecture [10], drought tolerance [11], leaf development [12] and Jasmonate regulation [13].

An international effort to re-sequence Asian rice accessions known as the “3000 Rice Genomes Project” (3K RGP) has provided the rice community with a better understanding of Asian rice diversity and evolutionary history, as well as providing valuable knowledge to enable more efficient use of these accessions for rice improvement [14, 15]. However, only 56 of these accessions originated from Vietnam, suggesting that the rice diversity within this country may not be fully captured within the 3K RGP. While the original 3K RGP analysis described nine subpopulations [15], subsequent reanalysis had shown that the 3K RGP could be further subdivided into fifteen subpopulations [16].

In this paper, we newly sequenced 616 Vietnamese rice accessions using whole-genome sequencing (WGS), most of them being native landraces. 164 of these rice accessions were in common with a previous study [8] based on a genotyping-by-sequencing (GBS) approach. We supplemented this dataset with all 56 Vietnamese genotypes from the 3K RGP to form a native diversity panel. We analysed this diversity panel of 672 accessions to explore the history of rice breeding in Vietnam, which is reflected in detectable changes in the allele frequency at sites under selection and their flanking regions. We also carried out a comprehensive analysis of the population structure of the combined 3,635 rice genomes obtained from joining our diversity panel and the complete 3K RGP datasets. We completed a GWAS on the diversity panel with 672 accessions (and separately for the Japonica and Indica subtypes within it) on thirteen phenotypes, which are available for around two-thirds of the samples. Finally, we looked for regions of selection between the subpopulations within Vietnam to reveal 200 regions spanning 7.8% of the genome, which might reflect their adaptation to local agricultural practices and farming conditions [17]. We used a similar approach to the following two studies; a comparison of upland and irrigated rice accessions to identify ecotype differentiated regions related to phenotypic differences [18], and a comparison of Indica semi-dwarf modern bred varieties (IndII) with taller Chinese landraces (IndI).

Our results highlight genomic differences between traditional Vietnamese landraces, which are likely the product of adaption to multiple environmental conditions and regional culinary preferences in a very diverse country.

## Results

### Sequencing rice diversity from Vietnam

Whole-genome sequencing was carried out on 616 rice accessions. 511 of the accessions were obtained from the PRC (Plant Resource Centre, Hanoi, Vietnam, http://csdl.prc.org.vn), together with their passport data, which shows that they were collected from all eight administrative regions of Vietnam (Additional file 1: Table S1). The remaining samples were obtained from AGI’s collection (Agricultural Genomics Institute, Hanoi, Vietnam). Three reference accessions (Nipponbare, a temperate Japonica; Azucena, a tropical Japonica; and two accessions of IR64, an Indica) obtained from the PRC, were included in the dataset. A total of 1,174 Giga base-pairs (Gbps) of data was generated for the 616 samples representing an average sequencing depth of 30x for 36 “high coverage” samples and 3x for 580 “low coverage” samples (Additional file 1: Table S1). These 616 newly-sequenced accessions were classified into 379 Indica and 202 Japonica subtypes, with the remaining 35 (including the Aus and Basmati varieties) being classified as admixed, based on the STRUCTURE [19] output for K=2 using a subset of 163,393 SNPs.

### Population structure of rice within Vietnam

The population structure of rice within Vietnam was analysed using the diversity panel of 672 samples, comprising 616 newly sequenced accessions and 56 Vietnamese genotypes from the 3K RGP. We assigned the 672 samples to four Japonica subpopulations and five Indica subpopulations (Additional file 1: Table S1) using (i) the population structure information obtained from the STRUCTURE analysis (Fig. 1), (ii) the previous characterisation of a panel of Vietnamese native rice varieties using GBS [8], and (iii) the assessment of the optimal number of subpopulations (Additional file 2: Figure S1) using the method described in Evanno et al. [20]. Subpopulations were named as in Phung et al. [8], except that we considered the I6 subpopulation to be part of the I3 subpopulation. Although the previous study used a limited number of GBS markers, 129 of the 164 common samples were assigned to the same subpopulations in both studies. Most differences were due to samples being classified as admixed in either one of the studies. We classified 48 (11%) of the Indica (Im), and eight (4%) of the Japonica samples (Jm) as admixed. The reference varieties Nipponbare (Temperate Japonica), Azucena (Tropical Japonica), and IR64 (Indica) were classified as J4, J1 and I1, respectively.

**Figure 1:**
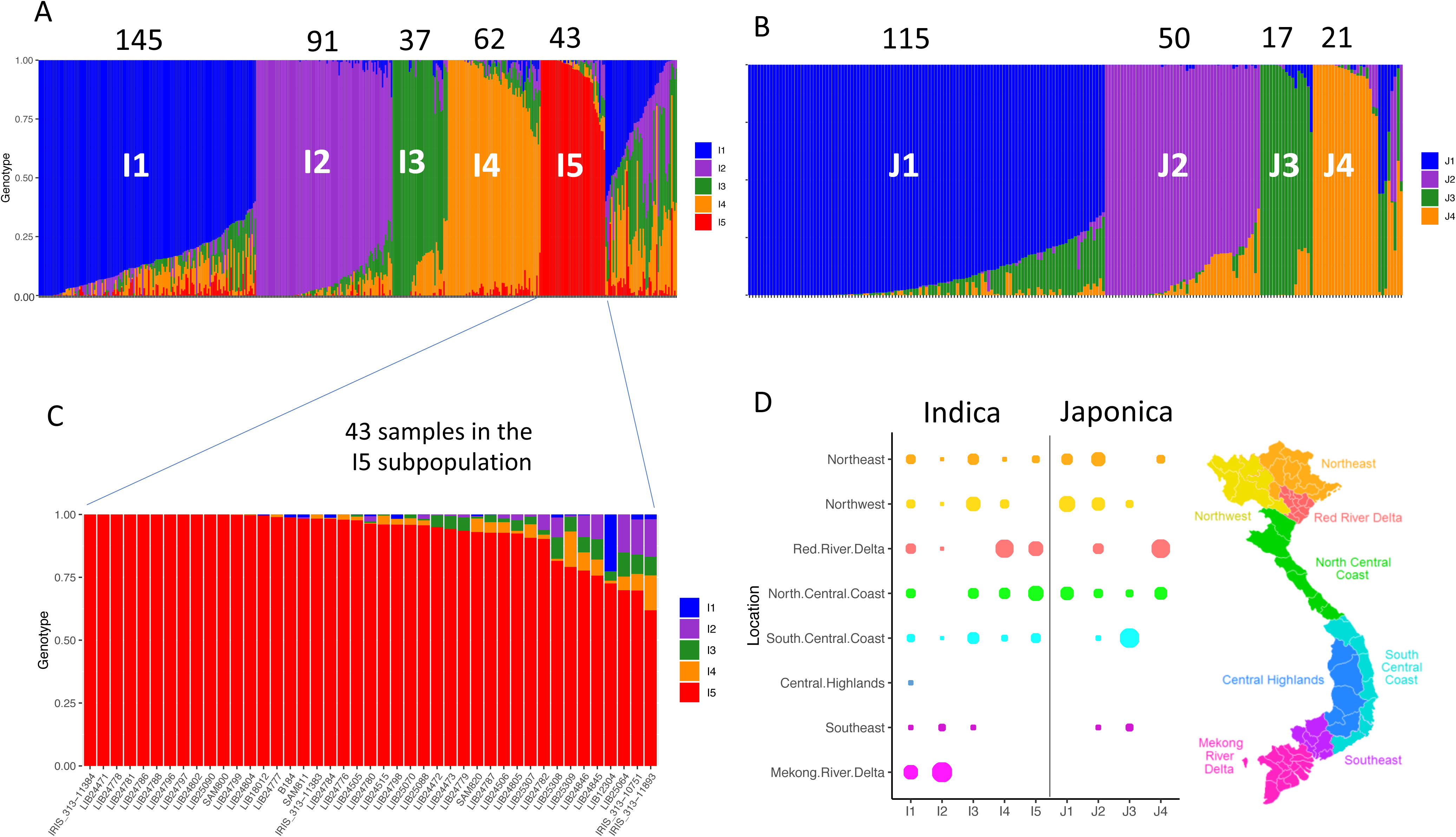
Population structure and location of the Indica and Japonica subpopulations within Vietnam. **a** STRUCTURE results (mean of 10 replicates) at K=5 for 426 Indica subtypes. Each colour represents one subpopulation. Each accession is represented by a vertical bar and the length of each coloured segment in each bar represents the proportion contributed by each subpopulation. The cut off for inclusion in each subpopulation is 0.6. The number of samples in each subpopulation is shown above, a further 48 samples were classified as admixed. **b** STRUCTURE results (mean of 10 replicates) at K=4 for 211 Japonica subtypes. The cut off for inclusion in each subpopulation is 0.6. The number of samples in each subpopulation is shown above, a further 8 samples were classified as admixed. **c** STRUCTURE results for the I5 subpopulation expanded to show individual samples. **d** The proportion of each population originating from each of the 8 regions in Vietnam (based on a subset of 377 samples, 54% of Indica samples and 85% of Japonica samples).

Each Indica subpopulation contained shared ancestry (admixed components) with other Indica subpopulation (Fig. 1a). The admixed components are shown in detail for the 43 samples in the I5 subpopulation (Fig. 1c) namely 38 samples from our dataset and the following five samples from the 3K RGP; IRIS 313-11384 (IRGC 127275), B184 (IRGC 135862), IRIS 313-11383 (IRGC 127274), IRIS 313-10751 (IRGC 127577) and IRIS 313-11893 (IRGC 127519). The Japonica subtropical J1 subpopulation shared ancestry (between 0 and 25% of the genome) with the Japonica tropical J3 subpopulation, whereas the two temperate subpopulations, J2 and J4 shared ancestry dominantly with each other. The tropical J3 subpopulation contained four samples with around 20% of the haplotypes in common with the temperate J4 subpopulation. Using the passport information available from the PRC, the proportion of each subpopulation originating from each of the “administrative regions” of Vietnam is shown in Fig. 1d. Only the I1 and I2 Indica subpopulations were collected from the Mekong River Delta regions, I2 being almost exclusively grown there whereas I1 was more widespread than I2. The I4 and J4 subpopulations were mainly collected from the Red River Delta areas. The J1 and J3 subpopulations were closely related; the J1 subpopulation was predominantly from the North of Vietnam whereas the J3 subpopulation was concentrated around the South-Central Coast region. Small variations in the percentage of reads mapping were observed for each of the subpopulations (Additional file 2: Figure S2).

A Principal Component Analysis (Fig. 2a and 2b) showed the relationship between these nine Vietnamese subpopulations [16]. Concerning the Vietnamese genotypes from the 3K RGP dataset included in the diversity panel, the Indica I1 subpopulation included two XI-1B modern varieties and eight admixed (XI-adm) accessions. I2 included fourteen XI-3B1 genotypes, which comprises Southeast Asian accessions, and similarly, I3 and I4 included one and ten XI-3B2 genotypes, respectively. Finally, I5 included five XI-adm accessions and clustered distinctly away from all the other subpopulations (Fig. 2a). On the other hand, J1 included the two subtropical (GJ-sbtrp) accessions from the Vietnamese 3K RGP genotypes, and J3 included one tropical (GJ-trp1) accession from the Vietnamese 3K RGP genotypes (Fig. 2b). These results correlate well with the latitudinal distinction between these subpopulations. J2 and J4 included two and one temperate (GJ-tmp) accessions, respectively; and split into two clear subpopulations in Vietnam compared with the East Asian temperate subpopulation described by the 3K RGP.

**Figure 2:**
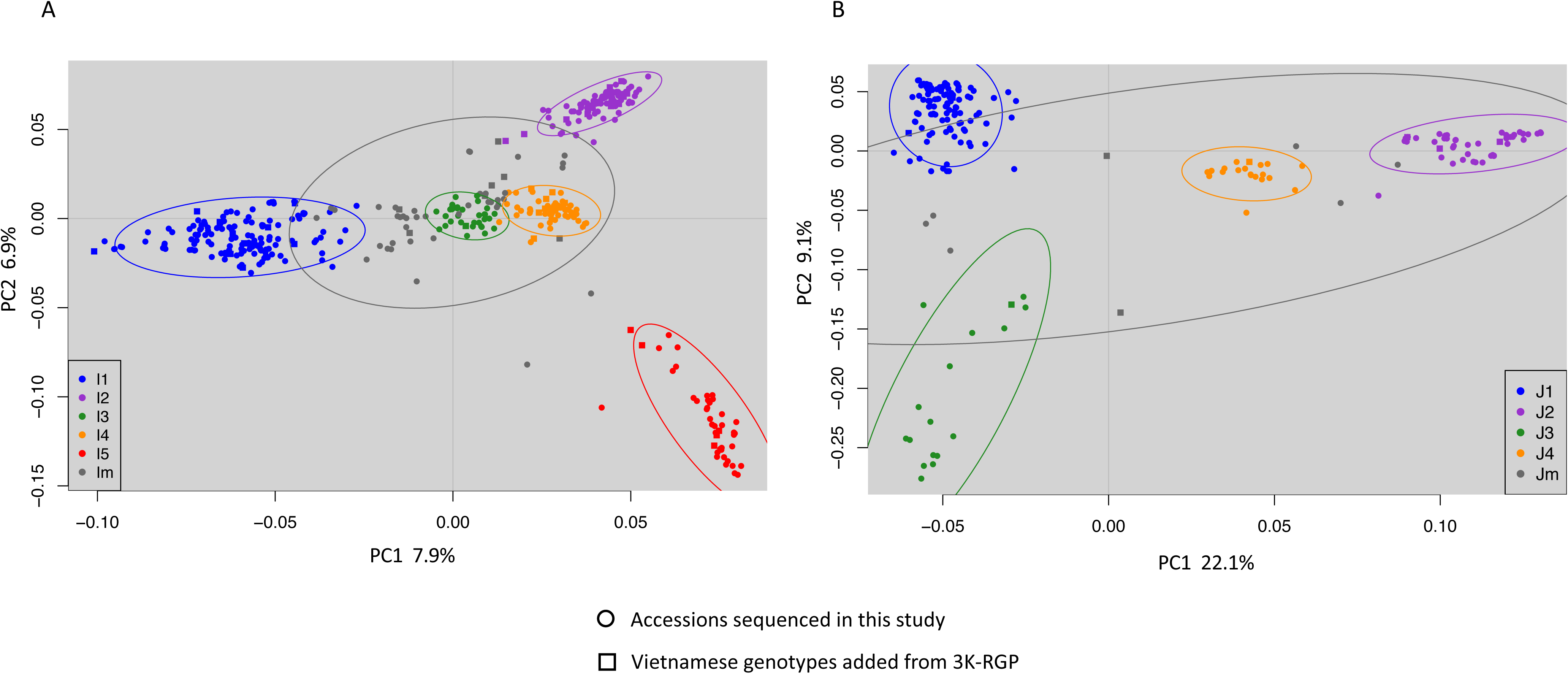
PCA analysis of Indica and Japonica Vietnamese subpopulations. **a** PCA analysis of 426 accessions from Vietnam using the top two components to separate the five Indica subpopulations. The ellipses show the 95% confidence interval. **b** PCA analysis of 211 accessions from Vietnam using the top two components to separate the four Japonica subpopulations. The ellipses show the 95% confidence interval.

### Population structure of the combined 3,635 Asian cultivated rice genomes

612 of the 616 newly sequenced accessions from this study and the 3,023 accessions from the 3K RGP were combined and classified into 9 and 15 subpopulations (Additional file 1: Table S2), and compared with the subpopulations from the 3K RGP analysis [15, 16]. For clarity, we used the prefix Jap- and Ind- to label these subpopulations from our analysis.

When the combined dataset of 3,635 samples was classified into nine subpopulations (Figure S3a), we found that 95% of the 3K RGP accessions (2,882 out of 3,023) were assigned into the same subpopulations. The remaining 5% lines were either (i) previously classified as admixture and our analysis placed into a subpopulation, or (ii) were previously classified in a subpopulation and were now classified as admixture. The 612 newly sequenced Vietnamese accessions were placed in three Indica clusters (187 accessions), three Japonica clusters (176 accessions), the Basmati and Sadri aromatic cB group (11 accessions), or the Aus cA subpopulation (one accession). In more detail, the three Indica clusters included three Im accessions in the East Asian cluster (Ind-1A), seventy-six I1 accessions in the cluster of modern varieties of diverse origins (Ind-1B), and 108 accessions (I2, I3 and Im) in the Southeast Asian cluster (Ind-3). Whereas, the three Japonica clusters included 54 accessions (J2, J4 and Jm) in the primarily East Asian temperate cluster (Jap-tmp), 119 accessions (J1, J3 and Jm) in the Southeast Asian subtropical cluster subpopulation (Jap-sbtrp) and three J3 accessions in the Southeast Asian Tropical subpopulation (Jap-trp). Any remaining accession with admixture components over 65% either Indica or Japonica were classified as Ind-adm (191 accessions) or Jap-adm (27 accessions), respectively. Finally, the remaining accessions were considered as Admix (19 accessions). Notably, all thirty-seven I5 accessions were placed in Ind-adm, and ten of the sixteen J3 accessions were placed in Jap-adm.

When the combined dataset of 3,635 samples was reclassified into 15 subpopulations (K15_new, Figure S3b), we noticed the following differences in the distribution of subpopulation compared to the 3K RGP analysis for the same number of 15 subpopulations (K15_3KRGP); we did not observe the division of the Aus samples into cA-1 and cA-2, and we subdivided the Indica subtypes and Japonica subtypes into eight and five subpopulations, respectively. A Principle Coordinate (PCO) analysis of the Indica and Japonica subpopulations is shown in Fig. 3, highlighting our new eight Indica and five Japonica subpopulations (In addition the Vietnamese and 3K RGP subpopulations are shown in Figures S5 and S6).

**Figure 3:**
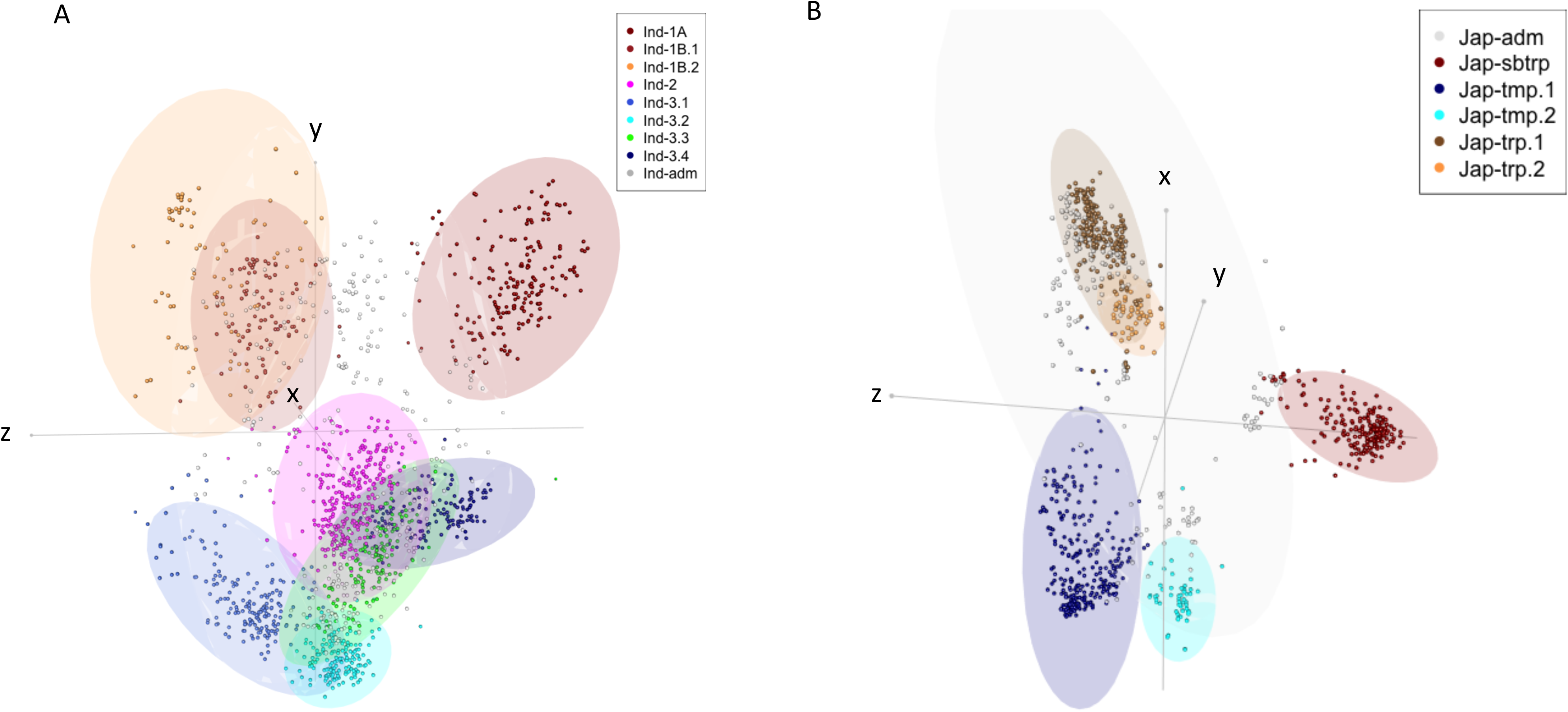
PCO analysis of Indica and Japonica Vietnamese subpopulations. **a** PCO analysis of 1605 Indica samples (omitting the samples classified as XI-adm and Ind-adm outside Vietnam for clarity). The ellipses show the 95% confidence interval for the K15_new subpopulations (the K15_3KRGP and five Vietnamese Indica subpopulations are shown in Figure S5). X = PC1, Y=PC4, Z=PC5. **b** PCO analysis of 982 Japonica samples (omitting the samples classified as GJ-adm and Jap-adm outside Vietnam for clarity) showing the K15_new subpopulations (the K15_3KRGP and four Vietnamese Japonica subpopulations are shown in Figure S6) X = PC3, Y=PC4, Z=PC5.

The relation between the subpopulations in our comprehensive analysis (3,635 accessions) and the 3K RGP (3,023 accessions) was as follows: (i) The Ind-1A, Ind-1B.1 and Ind-1B.2 were equivalent to XI-1A, XI-1B1 and XI-1B2, respectively. Forty-three of the Vietnamese I1 accessions were in the Ind-1B.1 subpopulation, and the remaining 102 I1 accessions were classified as admixed. (ii) The Ind-2 was equivalent to XI-2A and XI-2B, and as expected, this geographically distant South Asian subpopulation was not present in Vietnam. (iii) The previously observed split of the Indica-3 subpopulation into 3A and 3B was also observed in our analysis, where Ind-3.1 was equivalent to XI-3A and did not contain any Vietnamese accessions. (iv) The remaining Ind-3.2, Ind-3.3 and Ind-3.4 were a rearrangement of the XI-3B1 and XI-3B2 subpopulations. (v) The 89 Vietnamese I2 accessions belonged to Ind-3.2, which was a subset of XI-3B1. (vi) Ind-3.3 contained 16 of the 37 Vietnamese I3 accessions. (vii) 72% of the accessions in Ind-3.4 were from Vietnam, which contained 13 of the 37 I3 accessions, 61 of the 62 I4 accessions, and all I5 accessions. Within Ind-3.4, the admixture components of I3, I4 and I5 subpopulations (Figure S7) showed that I3 accessions were highly admixed, some I4 and I5 accessions were completely within Ind-3.4, while other I4 and I5 accessions showed admixture with Ind-3.3 (I5) or Ind.2, Ind-3.2, and Ind-3.3 (I4). To clarify these relations, a principle component analysis (PCA) with a reduced number of accessions was carried out using the 723 sample dataset (672 Vietnamese accessions and 51 genotypes from neighbouring Southeast Asian Countries; Figure S8), this supported the close relationships of I2 with XI-3B1, I4 with XI-3B2, I5 with XI-adm, J1 with GJ-sbtrp, and that both J2 and J4 were within GJ-tmp.

### Phenotypic and genetic diversity analysis of the Vietnamese Indica and Japonica subpopulations

Phenotypic measurements for 19 traits were scored in field conditions in the Hanoi area by breeders from the Agricultural Genomics Centre (AGI) for approximately two-thirds of the samples in our study. For five of these traits, additional scores were also included from trials by the Vietnamese Plant Resource Centre. In addition, phenotypic data were available for eleven of the traits in 38 of the 56 samples sourced from the 3K-RGP dataset (Additional file 1: Table S3, Table S4). Finally, the grain length to grain width ratio (GL/GW) was calculated to give a total of 20 traits (Additional file 1: Table S5). Scores were available for between 328 and 503 of the 672 samples (Indica subpanel, 170 – 297 samples and Japonica subpanel, 134 – 178 samples).

There were significant differences in measurements between the Indica and Japonica subtypes for ten of the traits; these are detailed in Additional file 1: Table S5 and histograms are shown in Fig. 4 for selected phenotypes. The Indica subtypes had significantly (p-value <0.0001) higher values for grain length to width ratio, leaf pubescence, culm number, culm length, and floret pubescence. In contrast, the Japonica subtypes had significantly higher values for grain width, leaf width, flag leaf angle, panicle length, and floret colour. The Indica I1 subpopulation (mostly elite varieties) was the most phenotypically distinct when compared to the rest of the Indica samples (mostly native landraces). I1 samples had longer grains (p-value = 2.2e-16), earlier heading date (p-value = 9.9e-12), higher culm strength (p-value = 2.2e-16), shorter leaf length (p-value = 2.7e-14) and shorter culm length (p-value < 2.2e-16). Similar values were obtained when comparing I1 to just the I5 subpopulation (Fig. 4). The I5 subpopulation was not phenotypically distinct (p-value < 0.001) from the other landrace subpopulations I2, I3 and I4, except for a significantly lower measurement of leaf pubescence (p-value = 0.0007). The Japonica J2 subpopulation had a significantly lower grain length to width ratio than J1 (p-value = 1.8e-13) and J3 (p-value = 5.7e-07). A correlation analysis carried out between the 20 phenotypes (Additional file 2: Figure S9) showed that the highest correlation (*r* = 0.6) was between leaf length and culm length (excluding the correlation between grain length to width ratio and grain length and grain width). Histogram and correlation plots are available for the 13 traits used for the GWAS analysis in Additional file 2: Figure S10 comparing the Indica and Japonica subtypes and in Additional file 2: Figure S11 comparing subpopulations I1 and I5. Further boxplots showing the phenotypic distribution according to subpopulation for culm length, grain length, grain width and heading date are available in Additional file 2: Figure S12.

**Figure 4:**
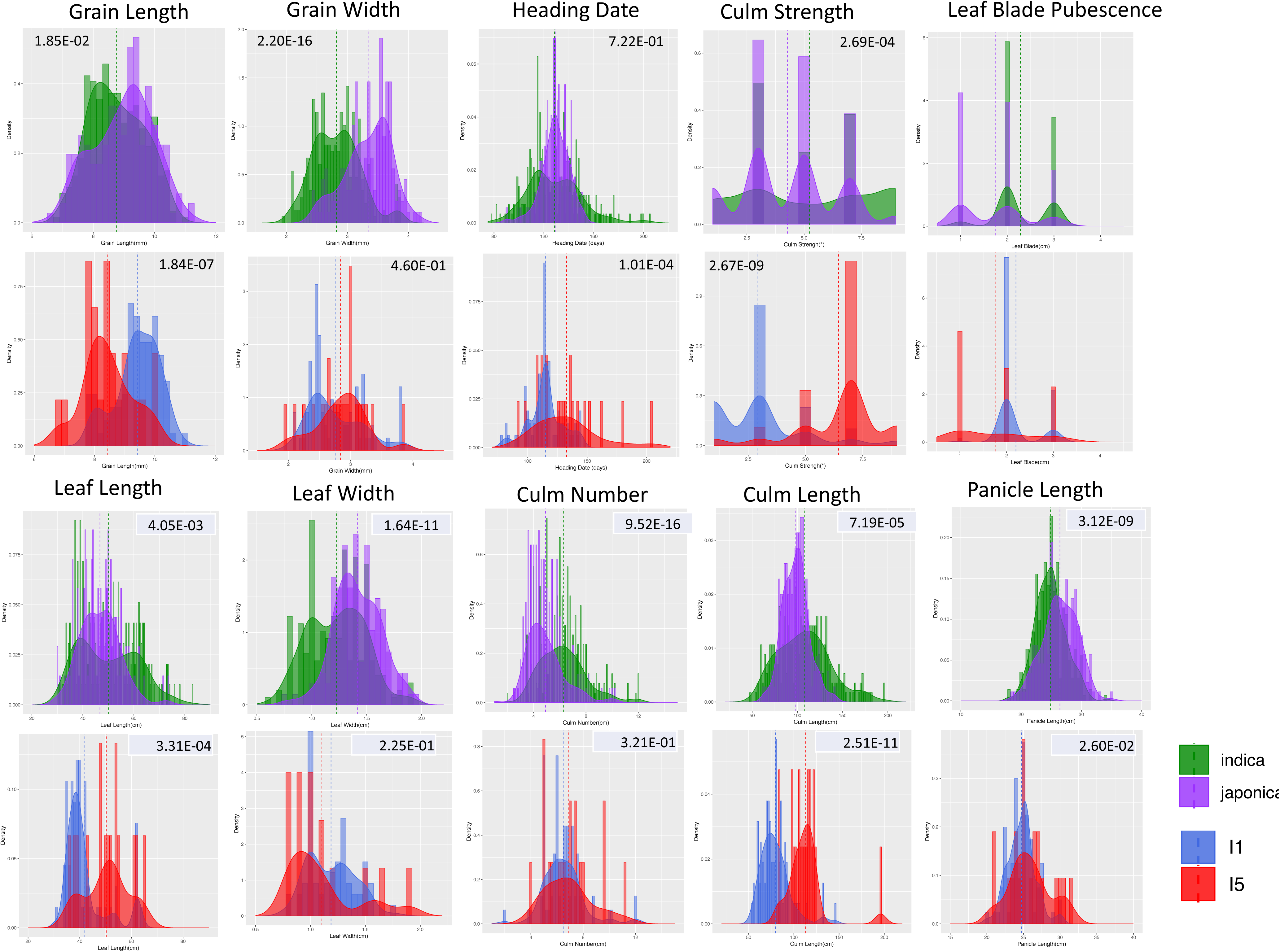
Histograms comparing the Indica and Japonica subtypes and the I1 and I5 subpopulations. Histogram are shown for 8 of the 13 traits used in the GWAS analysis. The Japonica and Indica subtypes are shown in green and purple respectively and underneath a histogram is shown for a subset of the Indica values comparing subpopulations I1 and I5. The mean value is shown by a dotted line and the p value (T-test) is shown at the top of each plot. A ggpairs histogram and correlation plot is available for all 13 traits in Additional file 2: Figure S7, Figure S8.

The Japonica subtypes had a lower nucleotide diversity (π = 0.000912) than the Indica subtypes (π = 0.00167). Looking at the individual subpopulations (Additional file 1: Table S6), the elite I1 subpopulation is the most diverse (π = 0.00144), and the I5 subpopulation is the least diverse (π = 0.00103). Regions of the genome with low diversity in all Indica subpopulations, and regions with low diversity in specific subpopulations, were observed when plotting diversity along each chromosome (Additional file 2: Figure S13). The J3 subpopulation is the most diverse of the four Japonica subpopulations. (π = 0.000697). Large genomic regions with very low diversity were observed in chromosomes 2, 3, 4 and 5 in all Japonica subpopulations (Additional file 2: Figure S14).

### Genome-wide association analysis

Three independent GWAS were conducted using the full panel (672 samples, 361,191 SNPs), the Indica subpanel (426 samples, 334,935 SNPs) and the Japonica subpanel (211 samples, 122,881 SNPs). Thirteen (13) of the 20 traits were suitable for GWAS based on the variance (CV < 56% for the full panel). The full list of phenotypic measurements is available in Additional file 1: Table S3. We found 643 significant phenotype-genotype associations. These associations were organised into 21 QTLs (Table 1, Additional file 1: Table S7). The GWAS Manhattan and Quantile-Quantile plots are available in Additional file 3: Figure S17 and Additional file 4: Figure S18. The QTLs ranged from 41 kb (16_FP) to 3,148 kb (5_GS). The 21 QTLs contained 1,730 genes and covered a total of 11 Mbp over ten chromosomes, and contained 453 SNPs with a significant association to a trait in at least one diversity panel (Fig. 5). The list of genes within each QTL is available in Additional file 1: Table S8. Functional enrichment was found within 9 of the QTL (Additional file 1: Table S9).

**Figure 5:**
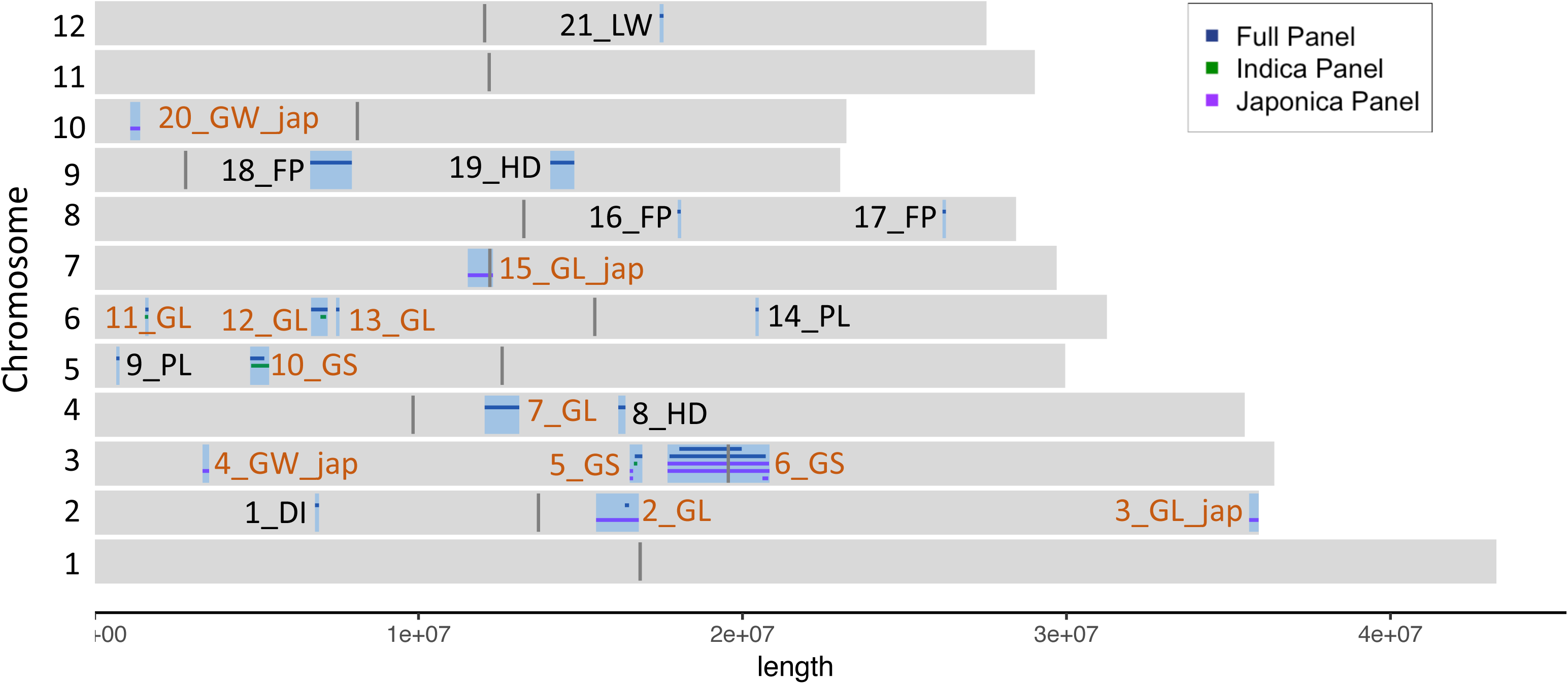
The distribution of 21 QTL. 21 significant associations for 8 of the 13 traits (-log_10_ (p value) ≥ 8.0). The 33 individual associations for the full panel and the Japonica and Indica subpanels were merged to form the 21 final QTLs. The QTLs for grain length, grain width and grain length/width ratio were merged into QTLs for grain size, these are labelled in brown. The remaining QTLs are labelled in black; Leaf width (LW), Panicle Length (PL), Heading Date (HD), Floret Pubescence (FP), Diameter Internode (DI). Regions smaller than 100 kb are extended to 50kb either side of SNP with maximum p value. Centromeric regions are shown as 100 kb regions in dark grey.

**Table 1:**
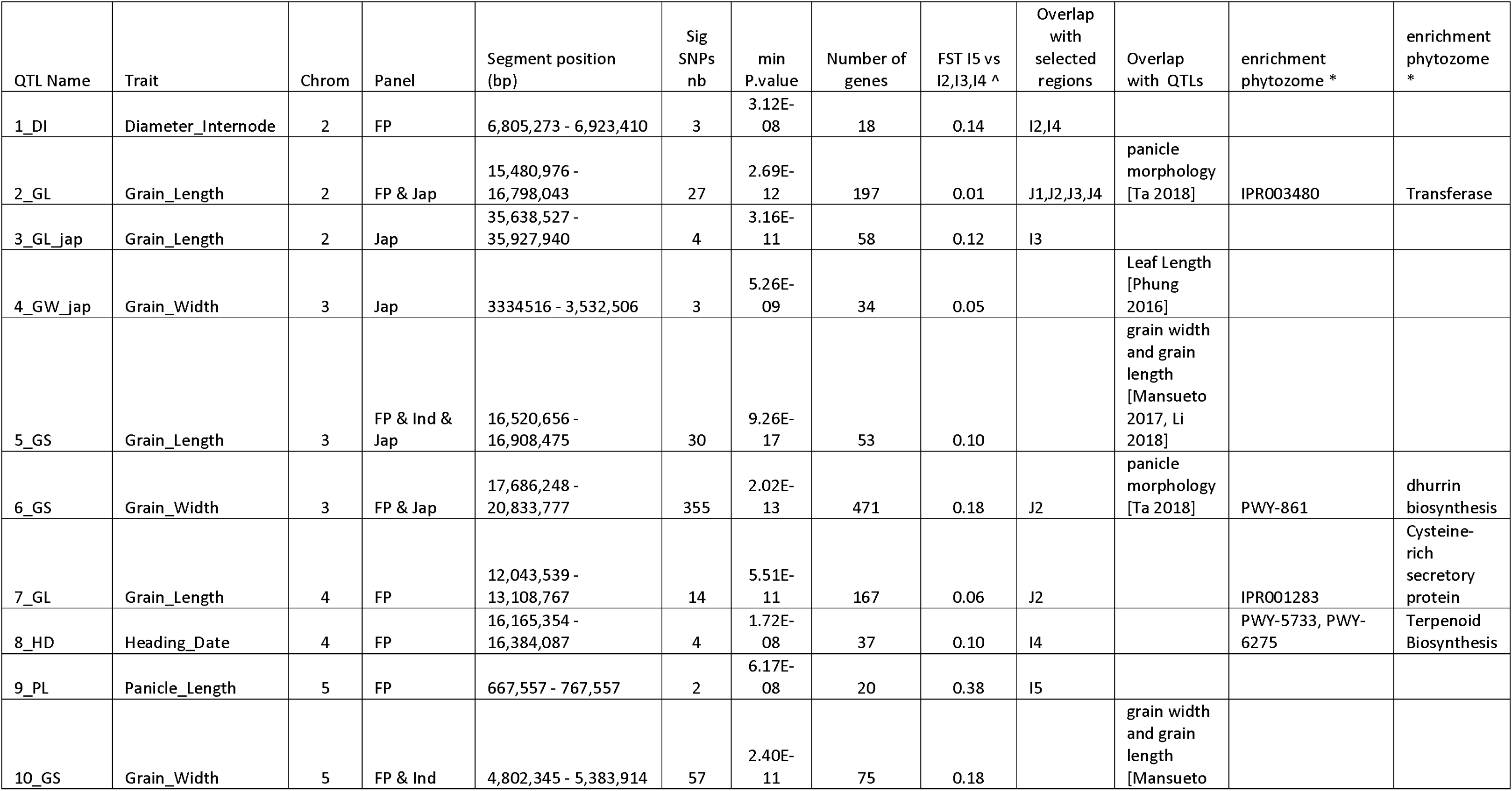
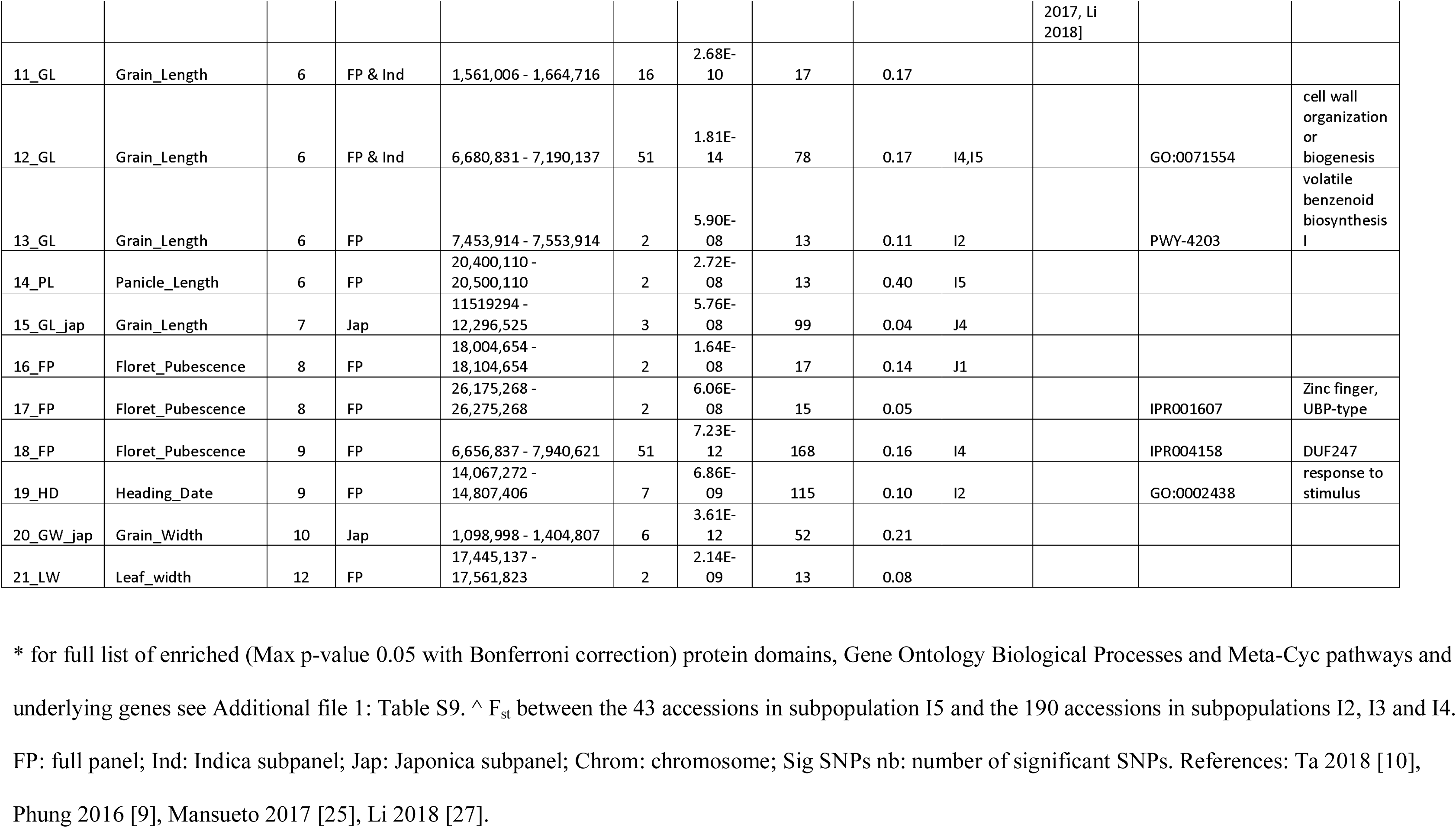
21 QTLs identified for plant description traits in the full panel, and Indica and Japonica subpanels. Detailing for the QTL analysis; significance threshold −log_10_ (p value) ≥ 8.0; panel in which significant associations were detected, highest level of significance for all panels, the occurrence of any overlap with selected regions in the four Japonica or five Indica subpopulations, any overlap with publish QTLs for Vietnamese rice populations or for the 3K RGP.

Seventeen QTLs were identified in the full diversity panel significantly associated with eight traits: grain length, grain width, grain length-to-width ratio, leaf width, panicle length, floret pubescence, heading date and internode diameter. A further 4 QTLs associated with grain length and grain width were observed only in the Japonica subpanel. Three of the QTLs, which were found in the full panel, were also observed in the Indica subpanel.

The set of 3.8M SNPs (see methods), representing one SNP every 99 bases, was annotated based on the potential effect of each SNP in protein function using SnpEff (Additional file 1: Table S3). 526,138 (4.79%) of the SNPs were in genes. There were 21,639 (0.197%) SNPs in 11,125 genes classified as having a putative “*High* impact” effect (E.g. Exon changes, frameshifts, gene fusions or rearrangements, protein structural changes, etc.). Following additional minimal allele frequency (MAF) filtering, in the Indica dataset (MAF 5%, 2,027,294 SNPs), there were 11,906 “*High* impact” SNPs in 7,396 genes and the Japonica dataset (MAF 5%, 1,125,716 SNPs), there were 6,240 “*High* impact” SNPs in 4,439 genes of which 2,818 were present in both Indica and Japonica.

None of the 453 SNPs with a significant association was annotated as resulting in protein changes (“*High* impact” SNPs). However, “*High* impact” effects were identified in other SNPs within the QTL. Among the total 1,730 genes in the 21 QTLs, we annotated 309 genes with “*High* impact” SNPs in the Indica subpanel, 248 genes with “*High* impact” SNPs in the Japonica subpanel, including 137 “*High* impact” SNPs common between the two sets. 129 of the 309 genes and 94 of the 248 genes had functional annotations in PhytoMine [21], but no functional overrepresentation was found for these sets of genes. In addition, we looked for overlaps with the QTL in five published Vietnamese studies [9–13], which used 25,971 SNPs in 182 samples (164 in common). We found that 2_GL and 6_GS overlapped with QTL for panicle morphological traits [10]; 2_GL overlapped with QTL9 for secondary branch number, and spikelet number (SBN and SpN), and 2_GS overlapped with QTL12 for secondary branch average length (SBL). 4_GW_jap overlapped with “q1” for longest leaf length (LLGHT) [9].

### Differential selection between subpopulations

To identify genomic regions which have been selected during the breeding of rice within Vietnam, we searched for genomic regions with distorted patterns of allele frequency that cannot be explained by random drift using XP-CLR [22]. Selected regions between pairs of either Indica or Japonica subpopulations were identified first. These regions were subsequently merged into a final set of selected regions for each subpopulation when regions were found to be selected against at least three subpopulations for Indica (Additional file 1: Table S10, Additional file 5: Figure S19) or at least two subpopulations for Japonica (Additional file 1: Table S10, Additional file 5: Figure S20). Here, we describe the procedure in more detail for the comparison of the I5 subpopulation to the other four Indica subpopulations: I5 vs I1 yielded 207 regions with a mean length of 267 kbp (14.8% of the genome); I5 vs I2 yielded 120 regions with a mean length of 204 kbp (6.57% of the genome); I5 vs I3 yielded 14 regions with a mean length of 162 kbp (0.61% of the genome); I5 vs I4 yielded 122 regions with a mean length of 122 kbp (6.02% of the genome). Regions selected against three or more subpopulations were merged to give 52 selected regions in I5, these had a mean length of 584 kbp covering 30 Mbp, which represented 8.13% of the rice genome and contained 4,576 genes. The selected regions for all of the subpopulations are plotted along each of the chromosomes in Fig. 6a and 6b for the Indica and Japonica subtypes, respectively. The list of genes selected in each subpopulation is available in Additional file 1: Table S11. The list of genes selected for each of the 52 regions in subpopulation I5 is available in Additional file 1: Table S12. Functional enrichment was found within 34 of the 52 regions (Additional file 1: Table S13). The mean whole-genome XP-CLR scores for each comparison are summarised in Fig. 6c and 6d. The I5 subpopulation showed the highest XP-CLR score, with an average of 41.4. The I3, J4, J2 and I4 had XP-CLR scores from 28 to 20. The J1 and I1 subpopulations had the lowest XP-CLR scores of 10.5 and 7.6, respectively. Overall, a greater number of selected regions were identified in the Indica than in the Japonica subtypes. These selected regions were distributed throughout the genome, whereas in the Japonica subtypes fewer regions were observed concentrated in specific regions of the genome. To gain insights into which traits and underlying genes have been selected in these regions, we looked for the overlap of selected regions with the 21 QTLs (Table 1). Also, we looked for overlaps with the QTLs identified in the five Vietnamese rice studies relating to root [9] and panicle morphological traits [10], tolerance to water deficit [11], leaf mass traits [12] and growth mediated by Jasmonate [13] (Fig. 7. Additional file 1: Table S14 and Table S15).

**Figure 6:**
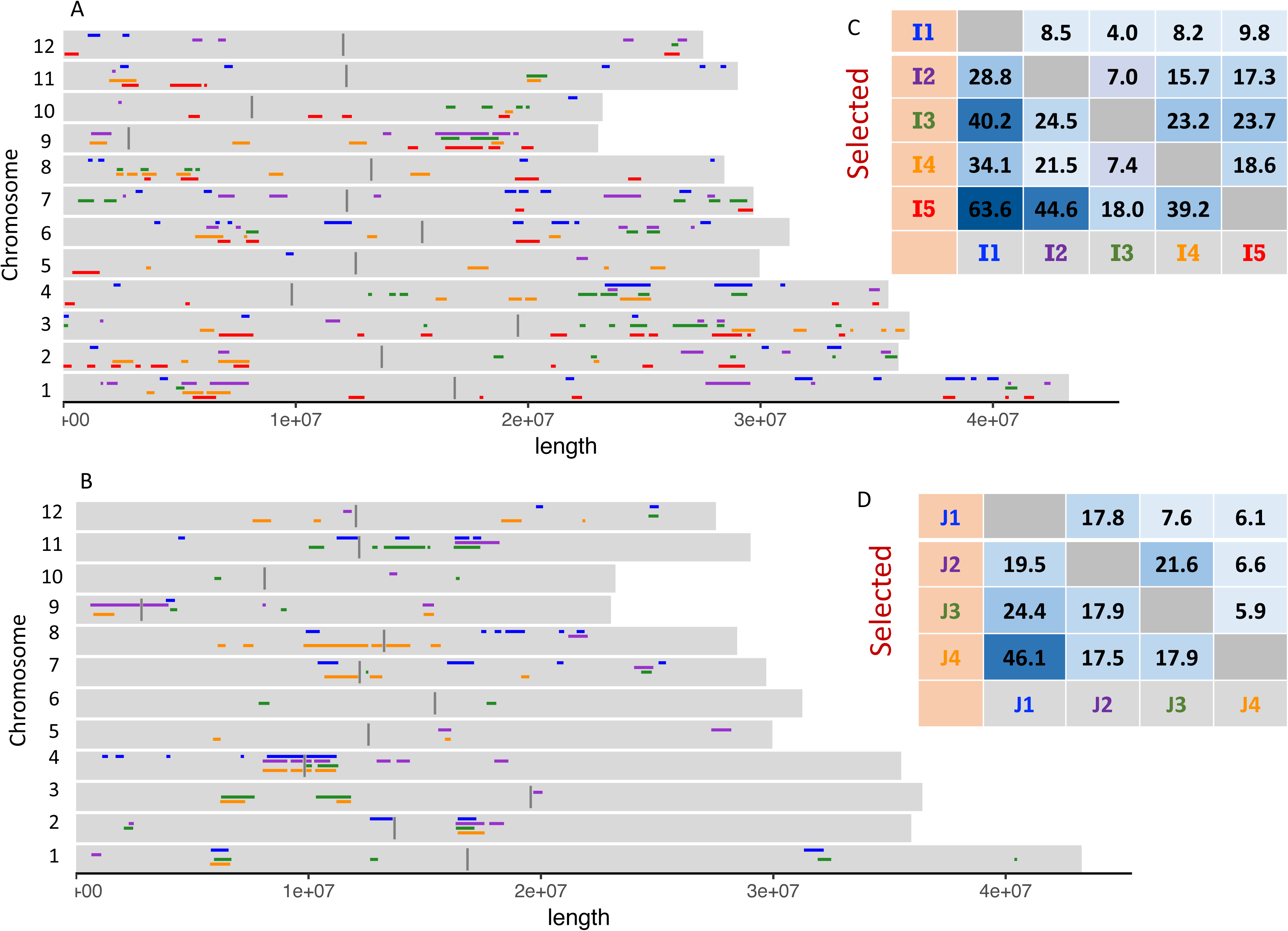
XP-CLR scores and regions of selection. **a** Selected regions for the five Indica subpopulations covering 5.4%, 6.1%, 5.3%, 6.3% and 8.1% of the genome for I1, I2, I3, I4 and I5 respectively. Centromeric regions are shown as 100 kb regions in dark grey. **b** Selected region for the four Japonica subpopulations covering 4.3%, 4.5%, 3.7% and 4.9% of the genome for J1, J2, J3 and J4 respectively. **c** and **d** Mean XP-CLR score across the whole genome for each comparison between all Indica (c) and Japonica (d) subpopulations. Darker colours indicate higher selection scores.

**Figure 7:**
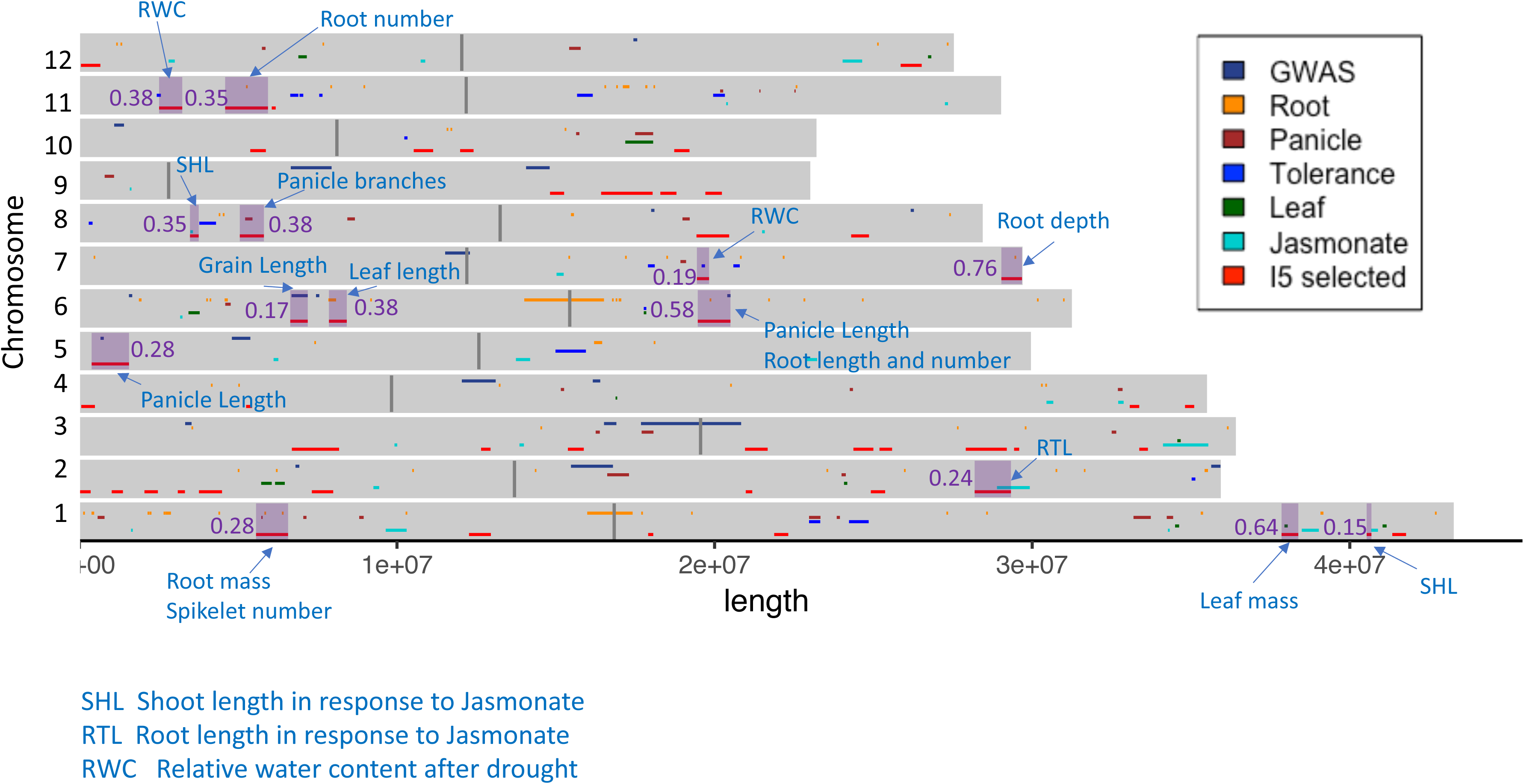
Vietnamese QTLs and their overlap with selected regions in the I5 subpopulation. QTLs from 5 published studies [9–13] and from this study are plotted along each chromosome. The QTLs which overlap with 14 of the regions selected in the I5 subpopulation are highlighted. The mean F_ST_ per region between the 43 samples in the I5 subpopulation and the 190 samples in the I2, I3 and I4 subpopulation is shown for these 14 regions.

To gain further information on the uniqueness of these regions selected in I5, we calculated the F_ST_ per SNP between the 43 samples in the I5 subpopulation and the 190 samples in the I2, I3 and I4 subpopulations. The mean F_ST_ per gene for the 4,576 genes selected in I5 is listed in Additional file 1: Table S16) and the mean F_ST_ per selected region is shown in Table 2. The 1,983,066 heterozygous SNPs in subpopulations I2, I3, I4 and I5 had a mean F_ST_ of 0.185, and this increased to 0.305 for the subset of 177,874 SNPs found within the I5 selected regions. Twenty-one genes with a putative role in salt tolerance in rice [23] fell within the regions selected in the I5 subpopulation (Additional file 1:Table S17). Fifty-six candidate genes were selected using the following criteria; F_ST_ over 0.5 for the whole selected region or for functionally enriched genes within regions, presence of “*High* impact” SNPs, and presence of candidate genes from overlapping QTL (Table 3). Allele plots for the “*High* impact” within genes are shown in Additional file 6: Figure S21.

**Table 2:** 52 regions under selection in the Indica I5 subpopulation. Detailing the overlap of selected regions with published QTLs for Vietnamese rice populations and the QTLs described in Table 1, selected regions in Indica and Japonica subpopulations, and published selected regions [Lyu 2014, Xie 2015].

| Region | Chrom | Segment position (bp) | <sup>^</sup> FST<br>I5 vs<br>I2,I3,I4<br>(a) | genes<br>per<br>region | Overlap<br>Indica<sup>*</sup> | Overlap<br>japonica<sup>**</sup> | Number of overlapping genes |  |  | Enrichment<br>phytozone |  | <sup>^</sup> FST<br>I5 vs<br>I2,I3,<br>I4 (b) | Overlap with QTLs<br>(c) |
| --- | --- | --- | --- | --- | --- | --- | --- | --- | --- | --- | --- | --- | --- |
|  |  |  |  |  |  |  | ecotype<br>differentia<br>ted<br>genes<sup>*</sup> | tall<br>(Ind1)<sup>*</sup> | semi-dwarf<br>(IndII)<sup>*</sup> | identifier | function |  |  |
| I5_1 | 1 | 5,563,164 - 6,569,946 | 0.28 | 138 | I2, I4 | J1, J3, J4 |  | 39 |  | PWY-6303 | methyl indole-3-acetate<br>interconversion | 0.22 | Root mass [Phung<br>2016], panicle<br>morphology [Ta<br>2018] |
| I5_2 | 1 | 12,270,588 - 12,957,024 | 0.33 | 94 |  | J3 |  |  |  | PWY-7445 | luteolin triglucuronide<br>degradation | 0.27 |  |
| I5_3 | 1 | 17,910,736 - 18,069,653 | 0.23 | 24 |  |  |  |  |  |  |  |  |  |
| I5_4 | 1 | 21,880,287 - 22,319,111 | 0.05 | 59 | I1 |  | 1 |  |  | IPR004883 | Lateral organ boundaries,<br>LOB | 0.03 |  |
| I5_5 | 1 | 37,850,965 - 38,378,420 | 0.64 | 84 | I1 |  |  |  |  | IPR004183 | Extradiol ring-cleavage<br>dioxygenase | 0.72 | Leaf mass [Hoang<br>2019] |
| I5_6 | 1 | 40,530,842 - 40,679,473 | 0.15 | 19 | I2, I3 |  |  |  | 19 | PWY-5980 | xylogalacturonan<br>biosynthesis | 0.18 | Jasmonate SHL [To<br>2019] |
| I5_7 | 1 | 41,340,158 - 41,769,828 | 0.31 | 65 |  |  |  |  |  | GO:0005975 | carbohydrate metabolic<br>process | 0.37 |  |
| I5_8 | 2 | 677 - 354,653 | 0.39 | 64 |  |  |  |  |  |  |  |  |  |
| I5_9 | 2 | 1,020,920 - 1,369,616 | 0.30 | 64 | I1 |  |  |  |  | IPR000864 | Proteinase inhibitor I13 | 0.51 |  |
| I5_10 | 2 | 2,050,537 - 2,469,973 | 0.24 | 60 | I4 | J2, J3 |  |  |  | IPR008930 | Terpenoid cyclases/protein<br>prenyltransferase alpha-<br>alpha toroid | 0.29 |  |
| I5_11 | 2 | 3,101,251 - 3,324,479 | 0.25 | 44 |  |  |  |  |  | IPR001611 | Leucine-rich repeat | 0.18 |  |
| I5_12 | 2 | 3,765,987 - 4,489,973 | 0.25 | 114 |  |  |  |  |  | GO:0006952 | defense response | 0.24 |  |
| I5_13 | 2 | 7,320,174 - 7,989,585 | 0.07 | 81 | I4 |  |  |  |  | GO:0006979 | response to oxidative<br>stress | 0.05 |  |
| I5_14 | 2 | 20,981,182 - 21,185,348 | 0.44 | 26 |  |  |  |  |  |  |  |  |  |
| I5_15 | 2 | 24,920,001 - 25,369,666 | 0.12 | 62 |  |  |  |  | 8 | GO:0006559 | L-phenylalanine catabolic<br>process | 0.17 |  |
| I5_16 | 2 | 28,191,142 - 29,329,745 | 0.24 | 168 | I3 |  |  |  |  |  |  |  | Jasmonate RTL [To<br>2019] |
| I5_17 | 3 | 6,691,656 - 8,177,456 | 0.39 | 224 | I1 | J3, J4 |  |  |  | IPR000971 | Globin | 0.63 |  |
| I5_18 | 3 | 12,650,311 - 12,946,658 | 0.21 | 45 |  |  |  |  |  |  |  |  |  |
| I5_19 | 3 | 15,380,808 - 15,879,517 | 0.10 | 71 | I3 |  |  |  | 23 | IPR001563 | Peptidase S10, serine<br>carboxypeptidase | 0.06 |  |
| I5_20 | 3 | 20,960,124 - 21,669,162 | 0.25 | 114 |  |  |  |  |  | GO:0006813 | potassium ion transport | 0.18 |  |
| I5_21 | 3 | 24,370,632 - 24,999,498 | 0.23 | 74 | I1, I3 |  |  |  | 13 |  |  |  |  |
| I5_22 | 3 | 25,193,549 - 25,587,517 | 0.28 | 55 |  |  |  | 7 | 9 |  |  |  |  |
| I5_23 | 3 | 27,910,148 - 29,199,870 | 0.34 | 188 | I2, I3, I4 |  | 17 | 6 | 85 | GO:0023014 | protein phosphorylation | 0.49 |  |
| I5_24 | 3 | 29,431,523 - 29,589,724 | 0.45 | 21 | I4 |  | 1 | 2 |  |  |  |  |  |
| I5_25 | 3 | 33,372,038 - 33,639,859 | 0.42 | 45 | I3 |  |  |  |  |  |  |  |  |
| I5_26 | 4 | 62,390 - 489,186 | 0.23 | 62 |  |  |  |  |  | IPR006115 | 6-phosphogluconate dehydrogenase, NADP-binding | 0.21 |  |
| I5_27 | 4 | 5,251,107 - 5,436,839 | 0.12 | 25 |  |  |  |  |  | PWY-2981 | diterpene phytoalexins precursors biosynthesis | 0.10 |  |
| I5_28 | 4 | 33,073,892 - 33,369,648 | 0.43 | 46 |  |  |  |  |  |  |  |  |  |
| I5_29 | 4 | 34,813,879 - 35,098,724 | 0.62 | 44 | I2 |  |  | 8 |  |  |  |  |  |
| I5_30 | 5 | 386,347 - 1,563,159 | 0.28 | 190 |  |  |  |  |  |  |  |  | 9_PL |
| I5_31 | 6 | 6,640,258 - 7,189,250 | 0.17 | 80 | I1, I2, I4 |  |  | 7 |  |  |  |  | 12_GL |
| I5_32 | 6 | 7,860,166 - 8,418,475 | 0.38 | 70 | I3, I4 | J3 |  | 3 |  | PWY-6917 | vernolate biosynthesis III | 0.37 | Leaf length [Phung 2016] |
| I5_33 | 6 | 19,470,641 - 20,499,968 | 0.58 | 165 | I1 |  |  |  |  | IPR001841 | Zinc finger, RING-type | 0.74 | Panicle length [Ta 2018], root length and number [Phung 2016] |
| I5_34 | 7 | 19,443,608 - 19,825,988 | 0.19 | 54 | I1 | J4 |  |  |  | IPR021470 | Protein of unknown function DUF3123 | 0.17 | Water content after drought [Hoang 2019] |
| I5_35 | 7 | 29,030,233 - 29,677,525 | 0.76 | 97 | I3 |  |  |  |  |  |  |  | Root depth [Phung 2016] |
| I5_36 | 8 | 3,484,045 - 3,758,632 | 0.35 | 39 | I3, I4 |  |  |  |  |  |  |  | Jasmonate SHL [To 2019] |
| I5_37 | 8 | 5,052,017 - 5,809,093 | 0.38 | 127 | I3, I4 |  |  |  |  | IPR001929 | Germin | 0.55 | Panicle branches [Ta 2018] |
| I5_38 | 8 | 19,431,460 - 20,459,346 | 0.22 | 149 | I1 |  |  |  |  | IPR010683 | Protein of unknown function DUF1262 | 0.23 |  |
| I5_39 | 8 | 24,300,313 - 24,859,863 | 0.23 | 92 |  |  |  |  |  | IPR002935 | O-methyltransferase, family 3 | 0.24 |  |
| I5_40 | 9 | 14,820,651 - 15,259,615 | 0.24 | 61 |  | J2, J4 | 2 |  |  | IPR002867 | IBR domain | 0.32 |  |
| I5_41 | 9 | 16,430,191 - 18,049,085 | 0.54 | 252 | I2, I3 |  |  |  |  | PWY-6303 | methyl indole-3-acetate interconversion |  |  |
| I5_42 | 9 | 18,292,494 - 18,798,654 | 0.49 | 78 | I2, I3, I4 |  |  |  |  | IPR029071 | Ubiquitin-related domain | 0.89 |  |
| I5_43 | 9 | 19,710,325 - 20,229,472 | 0.11 | 83 |  |  |  |  |  | PWY-5176 | coumarin biosynthesis (via 2-coumarate) | 0.09 |  |
| I5_44 | 10 | 5,381,471 - 5,869,967 | 0.27 | 62 |  |  |  |  |  | PWY-6303 | methyl indole-3-acetate interconversion | 0.64 |  |
| I5_45 | 10 | 10,528,884 - 11,139,609 | 0.38 | 118 |  |  |  |  |  | IPR027923 | Hydrophobic seed protein | 0.76 |  |
| I5_46 | 10 | 11,991,467 - 12,409,929 | 0.26 | 69 |  |  |  |  |  |  |  |  |  |
| I5_47 | 10 | 18,732,199 - 19,209,687 | 0.51 | 80 | I4 |  |  |  | 64 | IPR002885 | Pentatricopeptide repeat | 0.48 |  |
| I5_48 | 11 | 2,510,079 - 3,239,747 | 0.38 | 109 | I1, I4 |  |  | 56 |  | GO:0050794 | regulation of cellular process | 0.45 | Water content after drought [Hoang 2019] |
| I5_49 | 11 | 4,590,276 - 5,937,318 | 0.35 | 200 |  | J1 |  | 3 | 14 | IPR001810 | F-box domain | 0.38 | Root number [Phung 2016] |
| I5_50 | 11 | 6,060,058 - 6,179,872 | 0.23 | 20 |  |  |  |  |  |  |  |  |  |
| I5_51 | 12 | 50,720 - 659,181 | 0.30 | 108 |  |  |  |  |  |  |  |  |  |
| I5_52 | 12 | 25,861,119 - 26,518,838 | 0.11 | 93 | I2, I3 |  |  |  |  |  |  |  |  |

\* for full list of enriched (Max p-value 0.05 with Bonferroni correction) protein domains, Gene Ontology Biological Processes and Meta-Cyc pathways and underlying genes see Additional file 1: Table S13. ^ $F_{ST}$ between the 43 samples in subpopulation I5 and the 190 samples in subpopulations I2, I3 and I4 (a) mean $F_{ST}$ for all the genes in the selected region. (b) mean $F_{ST}$ for the genes showing functional enrichment. (c) details in Additional file 1: Table S14 and S15. References Phung 2016 [9], Ta 2018 [10], Hoang 2019\_1 [12], To 2019 [13], Hoang 2019\_2 [11]. & Overlap with regions selected in other Indica subpopulations. && Overlap with regions selected in other Japonica subpopulations.

**Table 3:**
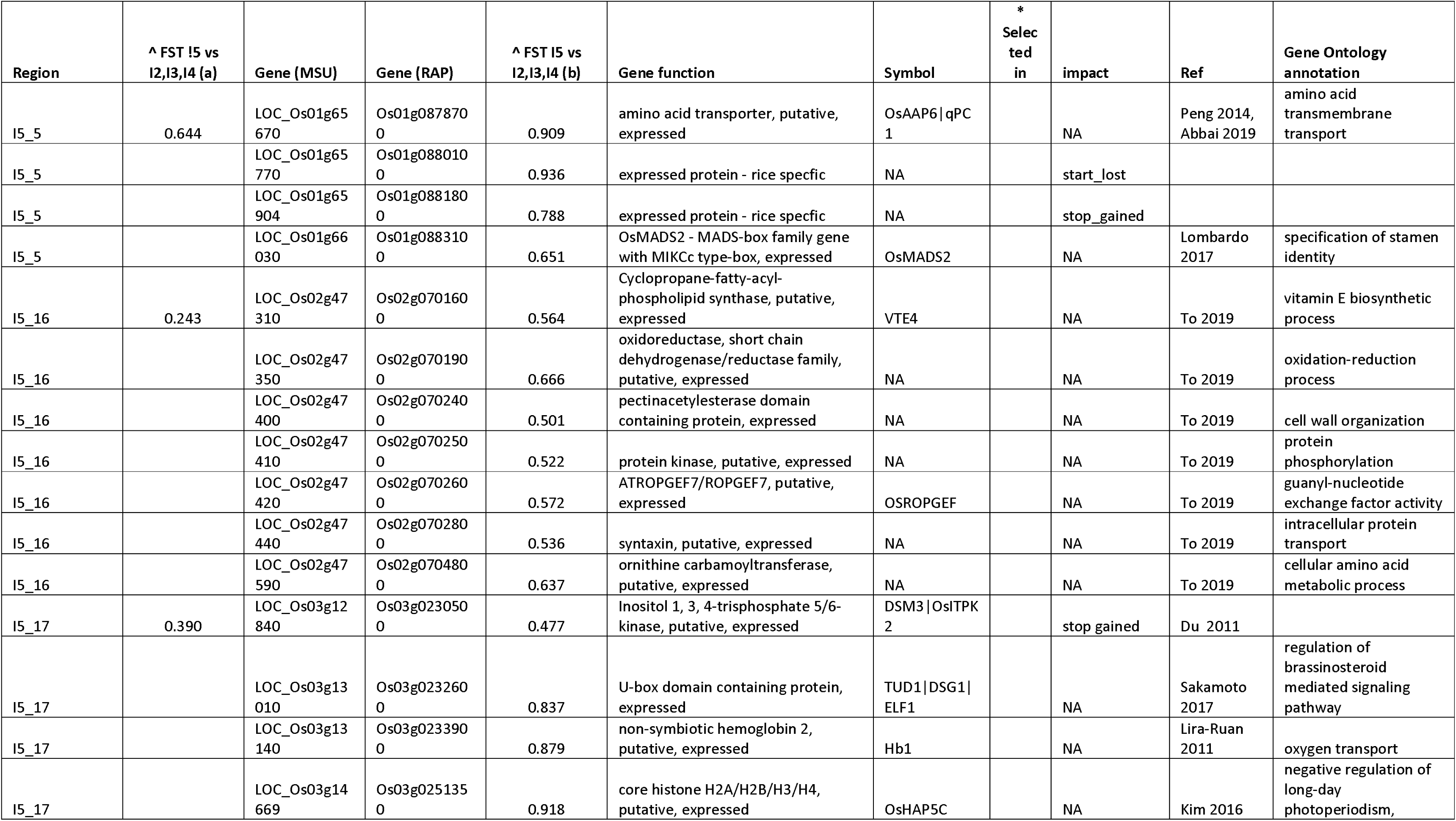
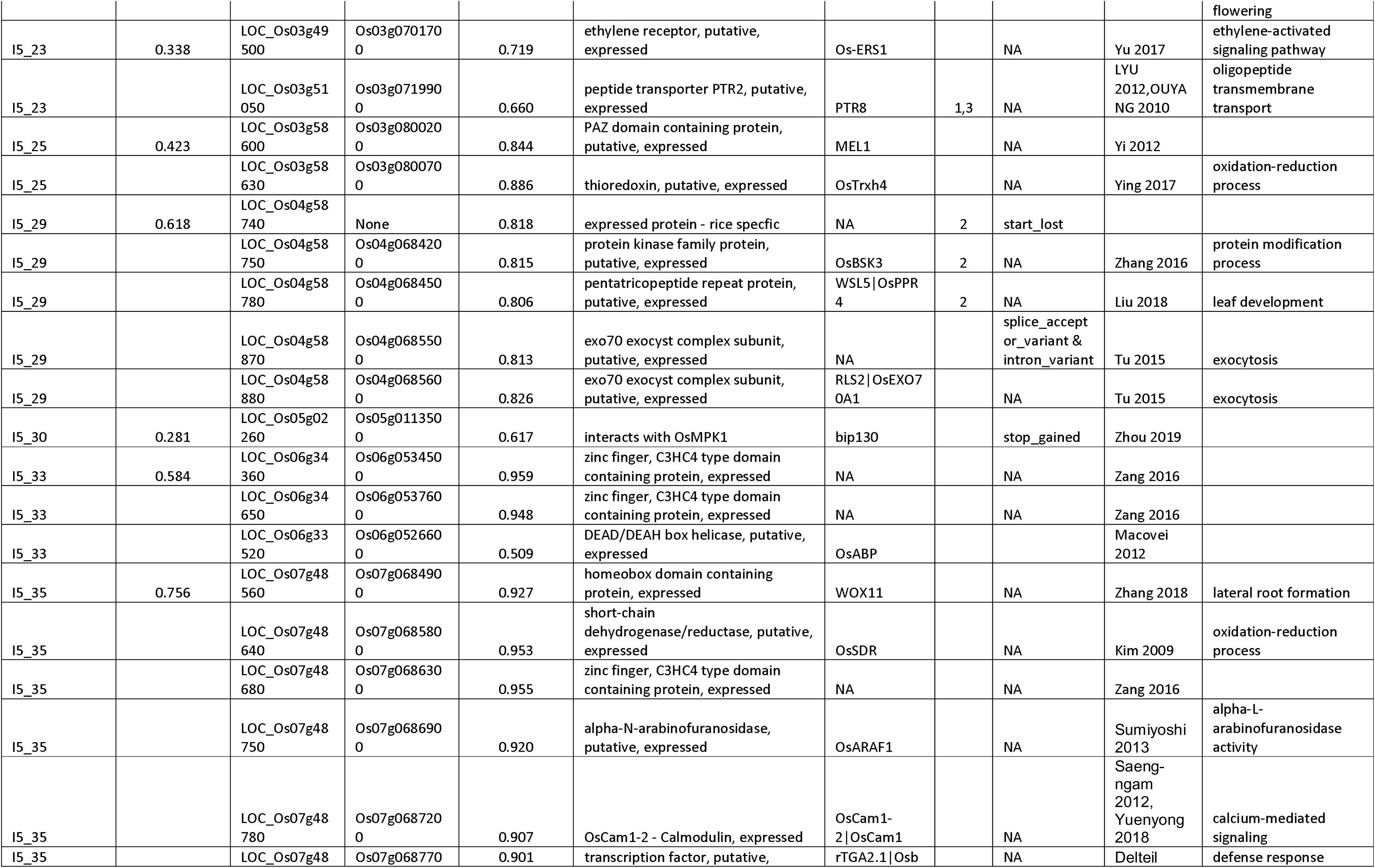
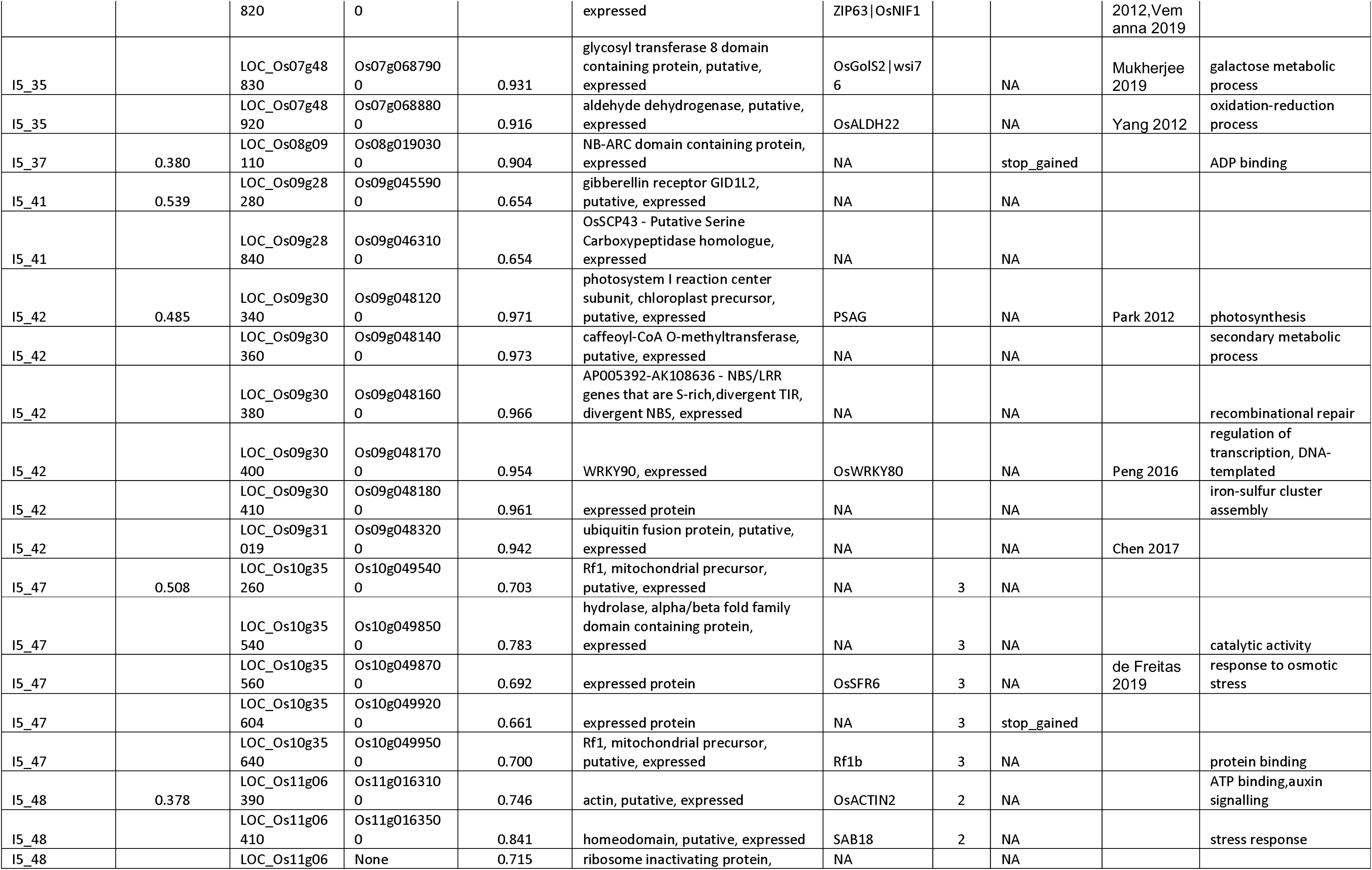
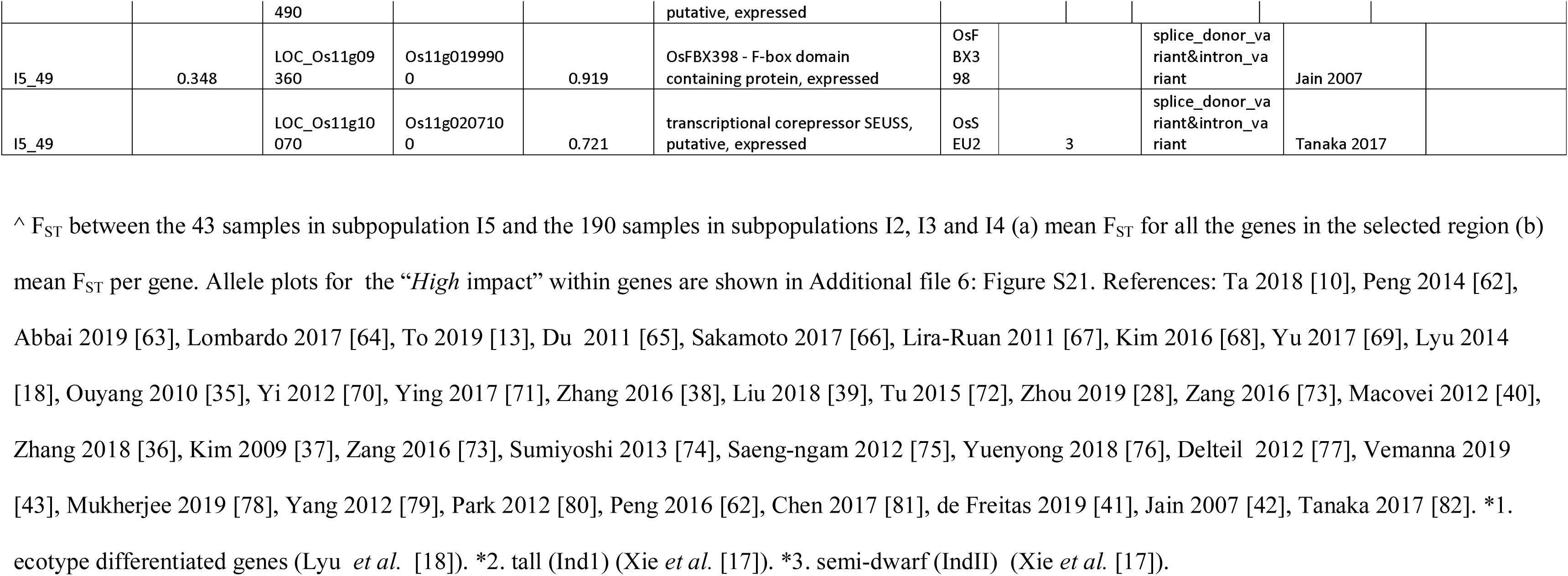
Candidate genes under selection in the Indica I5 subpopulation. Functional annotation of the 56 candidate genes and overlap with genes selected in previous studies [17, 18].

## Discussion

### Indica and Japonica rice subpopulations within Vietnam

Whole-genome sequencing of 616 Vietnamese rice accessions, predominantly landraces, plus 56 Vietnamese genotypes previously sequenced by the 3K RGP, provides us with a diversity panel to clarify the structure of rice subpopulations in Vietnam. Here, we describe five Indica subpopulations and four Japonica subpopulations using phenotypic measurements from this study, passport information available from the Vietnamese National Genebank (PRC), and the agronomic and geographical annotations from Phung et al. [8]. In general terms, our population structure within Vietnam agreed with the previous study, which used a smaller number of markers and 182 samples and is approximately a third of our diversity panel [8]. Subpopulation I1 is the most phenotypically distinct of the Indica subpopulations and shows typical phenotypes of ‘elite’ varieties, such as short height, strong culm strength, long slender grains and a short growth-duration (less than 120 days from sowing to harvest). I1 accessions are grown throughout Vietnam in irrigated ecosystems but predominantly in the Mekong River Delta in the south of the country. Subpopulation I2 is mainly composed of long growth-duration (over 140 days), tall varieties grown in the rainfed lowland and irrigated ecosystems of the Mekong River Delta with a broad diversity of grain shapes. The remaining three Indica subpopulations are intermediate between I1 and I2 for growth-duration, height and culm strength, have a broad diversity of grain shapes, and are not grown in the Mekong River Delta. Subpopulation I3 has the highest proportion of upland varieties but also includes some lowland varieties from the “South Central Coast” region many of which were classified as an independent subpopulation (I6) by Phung et al. [8]. Subpopulation I4 is mainly grown in the rainfed lowland and irrigated ecosystems of the Red River Delta. Subpopulation I5 is grown in a range of ecosystems but concentrated around the North Central Coast and Red River Delta regions, but excluding the Northwest region suggesting that it is the main lowland subpopulation. The J1 and J3 subpopulations are closely related upland varieties and the J2 and J4 subpopulations are closely related lowland varieties. Subpopulation J1 is mostly composed of medium growth-duration upland varieties from the mountainous regions in the North of Vietnam, with long large grains typical of upland varieties. Subpopulation J2 is grown throughout Vietnam in a range of ecosystems but has consistently short grains. Subpopulation J3 is mainly grown in the “South Central Coast” region and has long large grains. Subpopulation J4 is primarily grown in the Red River Delta region in lowland and mangrove ecosystems and has short grains.

The drought tolerance of these subpopulations can be inferred from the root traits measured by Phung et al. [9]The J1 and J3 upland subpopulations have deeper and thicker roots than the thinner shallower roots in the J2 and J4 subpopulations, which are grown in irrigated and mangrove ecosystems [9]. This suggests that the J1 and J3 subpopulations, which are grown mainly in rainfed upland regions, would be more drought tolerant than the others. Similarly, the I3 subpopulation has the deepest and thickest roots. It would, therefore, be more drought tolerant than the I1 and to a lesser extent the I5 subpopulation, which has the thinnest, shallowest root systems.

### A comprehensive analysis of the available 3,635 Asian cultivated rice genomes

The comprehensive analysis of the combined 3,635 Asian cultivated rice genomes obtained by joining our diversity panel with the full 3K RGP dataset resulted in a similar assignation to the previous 3K RGP analysis in 84 % of the cases. The largest differences were that the 3K RGP split the cA and XI-2 subpopulations, while our analysis split the GJ-tmp and rearranged the two XI-3B subpopulations into Ind-3.2, Ind-3.3 and Ind-3.4. The single temperate subpopulation (GJ-tmp) from the 3K RGP is further split in our study between the Jap-tmp.1 and Jap-tmp.2 subpopulations, with 88% of the samples in Jap-tmp.2 coming from Vietnam and forming the J2 subpopulation. These differences are likely due to changes in the distribution of genetic variants in subpopulations expanded within Vietnam.

### Vietnamese rice subpopulations in the context of the 3K RGP Asian cultivated rice subpopulations

The Indica I1 subpopulation, which contains a high proportion of elite varieties, clustered with the X1-1B1 subpopulation of modern varieties. The Southeast Asian native subpopulations (XI-3B1 and XI-3B2) clustered with the I2 and I4 subpopulations, respectively. I3 appeared to include both XI-3B1 and XI-3B2 accessions. The subpopulations from East and South Asia (XI-1A, XI-2A, XI-2B, XI-3A) had no representatives from Vietnam and fell outside of the Vietnamese subpopulation clusters, as expected. Our four Vietnamese Japonica subpopulations relate to the tropical (J1), subtropical (J3) and temperate (J2 and J4) Japonica subpopulations from the 3K RGP according to their latitudinal origin from South to North Vietnam, respectively.

The most exciting subpopulation is I5. When all 3,635 samples were considered, the subpopulation XI-3.4 included half of the I3, all but one of I4 and all I5 Vietnamese accessions, as well as half of the Southeast Asian native XI-3B2 genotypes from the 3K RGP. The remaining XI-3B2 were classified as Indica admix (Ind-adm). However, when only the Vietnamese samples were considered in the analysis, I5 clustered distinctly away from I3 and I4 subpopulations (Fig. 2A) and included five accessions from the 3K RGP, which had very low shared ancestry (admixture components) with other 3K RGP samples. Notably, Vietnamese landrace IRIS 313-11384 (IRGC 127275) had no shared ancestry with any other Vietnamese 3K RGP genotypes. Remarkably, a recent study on genomic signals of admixture and alien introgression in a core collection of 948 accessions representative of the earlier Asian Rice Landraces [24] included IRIS 313-10751 (IRGC 127577) and IRIS_313-11383 (IRGC 127274) from the I5 subpopulation.

### Genome-wide association analysis in Vietnamese rice landraces highlight 21 QTL

We have also extended upon five published GWAS [9–13], which focussed on specific traits but used a smaller number of markers and a third of the samples from the Vietnamese dataset. We took a similar approach of carrying out the analysis on both the full panel and the Indica and Japonica subpanels. Showing the QTL for the various traits altogether in Fig. 7 has highlighted some interesting overlaps. Notably, the overlap of QTL for panicle morphology with our QTL for grain size (2_GL and 6_GS). These previous studies found QTL in the full panel and in the Indica subpanel, but not in the Japonica subpanel. However, we found QTL for grain size that were only present in the Japonica subpanel, and all the QTL found in the Indica subpanel were also found in the full panel. These differences probably reflect our larger dataset. Comparing our results with the GWAS results from the 3K RGP (https://snp-seek.irri.org/) [25, 26], the QTL 5_GS on chromosome 3 is in the same region as a marker associated with grain length, and the QTL 10_GS on chromosome 5 is in the same region as a marker associated with both grain width and grain length. Underlying these two QTL, there are genes that have a putative role in the control of grain size in rice [27], namely GS3 (Os03g0407400) in 5_GS and GSE5 (LOC_Os05g09520, Os05g0187500) in 10_GS. We also looked for genes with “*High* impact” SNPs in QTL, relevant candidates include bip130 [28] (LOC_Os05g02260, Os05g0113500) with a stop gain mutation underlying the QTL 9_PL for panicle length and OsSPX-MFS3 (LOC_Os06g03860, Os06g0129400) [29] with a splice acceptor variant at the end of an intron underlying the QTL 11_GL for grain length.

### Breeding signatures between subpopulations focussing on the Indica I5 subpopulation

Unravelling the genomic differences between these described subpopulations, which are adapted to multiple environmental conditions and regional food preferences in Vietnam, provides an insight into the genomic regions associated with these adaptations. Selection causes detectable changes in the allele frequencies of the selected sites and their flanking regions. By jointly modelling loci allele frequency differentiation and frequency under neutrality and selection, the cross-population composite likelihood ratio test (XP-CLR) can detect selective sweeps [22]. These distorted patterns in allele frequency in contiguous SNP sites would have occurred too quickly (speed of change is assessed over expanding windows based on the length of the affected region) to be explained by random drift. XP-CLR has been used to identify regions of selection associated with domestication and improvement in a wide range of crops such as apple [30], soybean [31], cucumber [32] and wheat [33]. In rice, XP-CLR was used more specifically to compare upland and irrigated rice accessions [18] and to compare Indica semi-dwarf modern bred varieties (IndII) with taller Chinese landraces (IndI) [17] and revealed 200 regions spanning 7.8% of the genome, which might reflect their adaptation to local agricultural practices and farming conditions. We have used a similar approach to identify selected regions in all of the subpopulations, showing the strongest selection in the I5 subpopulation with fewer regions being selected overall in the Japonica subpopulations. We have examined the 52 selected regions in the I5 subpopulation in more detail. Specifically, we looked for overlaps with the selected genes identified in the above two studies (Lyu et al. [18] and Xie et al. [17]) using XP-CLR in rice. Moreover, to give us indications of the possible traits selected in these regions, we carried out a functional annotation of the regions and looked for overlaps with QTL.

Diversity is reduced when regions are under selection, but the observed diversity depends on many factors, including how long ago the selection occurred and the type of alleles selected alongside. This is referred to as the hitchhiking effect [34]. The fixation index (F_ST_) is a measure of population differentiation due to genetic structure. Both measurements vary highly along the genome but can provide additional information about the selected regions identified using XP-CLR. In this study, we calculated F_ST_ by comparing the I5 accessions to accessions in subpopulations I2, I3 and I4. We did not include the accessions in the elite I1 subpopulation, as we are specifically interested in genes that have been selected during the breeding of landraces within Vietnam.

Lyu et al. [18] identified 56 Indica-specific genes in selected regions which may account for the phenotypic and physiological differences between upland and irrigated rice. Thirty-one of these genes on chromosome 3 lie within regions also selected in the I4 and I5 subpopulations (I5_23, I4_24), the gene with the highest F_ST_ of 0.67 is *ptr8* (LOC_Os03g51050, Os03g0719900), which encodes a peptide transporter [35]. Xie et al. [17] identified 2,125 and 2,098 coding genes in regions selected in the Chinese landraces (IndI) and modern-bred (IndII) subpopulations, respectively. Comparing with the genes in selected regions in the I5 subpopulation evidenced an overlap of 131 genes with the 2,125 genes selected in the IndI subpopulation and an overlap of 235 genes with the 2,098 genes selected in the IndII subpopulation. This includes nine genes on chromosome 3, which were selected in all three subpopulations (7 genes in I5_22 and two genes in I5_23).

Of the 52 regions selected in the I5 subpopulation, the six with a mean F_ST_ over 0.5 were studied in more detail to highlight potential candidate genes. Notably, we identified the following genes in region I5_35; the transcription factor *WOX11* involved in crown root development [36] and *OsCam1*, *OsbZIP63*, and *OsSDR*, which have putative roles in defence [37]. Further genes of interest are *OsAAP6*, a regulator of grain protein content [39] in region I5_5, *OsBSK3* [38] and *WSL5* [39] which play a role in growth in region I5_29, *OsABP* which is upregulated in response to multiple abiotic stress treatments [40] falls within region I5_33 and *OsSFR6*, a cold-responsive gene [41] in region I5_47. Two of the genes contained “*high* impact” mutations, *OsFBX398*, an F-box gene with a potential role in both abiotic and biotic stresses [42, 43] in region I5_49 and *bip130* [28] in region I5_30 which regulates abscisic acid-induced antioxidant defence and fall within our QTL for panicle length (9_PL).

We have shown that subpopulation I5 constitutes an untapped resource of cultivated rice diversity. The analysis restricted to Vietnamese accessions allowed us to observe differences among the accessions within the country. Although 38 accessions (including two genotypes from the same accession in our study) are deposited in the PRC in Hanoi, and the remaining five accessions are available from the 3K RGP, there is limited information from the passport and phenotypic data to be able to understand the distinctiveness of this subpopulation fully. Further analysis of this subpopulation should encompass ‘Indica specific genes’ which may have been overlooked in our study as we used a Japonica reference. Phung et al. [8] described subpopulation I5 as “medium growth-duration accessions from various ecosystems of the North and South Central Coast regions, with rather small and non-glutinous grains”. Our I5 accessions are predominantly from the Red River Delta and contiguous coastal departments, the “North Central Coast” and “Northwest” administrative regions, but remarkably excluding the higher altitude Northwest region in the North, the more upper “Central Highlands”, as well as the whole Mekong River Delta in the south. This suggests that I5 accessions are common traditional low yielding lowland varieties with specific environmental or culinary values.

Vietnam is currently experiencing increasing variability in the local climate due to global changes and the growing severity of the El Nino-Southern Oscillation phenomenon, creating notable inter-annual variations in precipitation ranging from severe drought to large-scale floods [5]. The Mekong River Delta region is an essential region for rice production globally, but the adverse effects of salinisation have damaged rice production in recent decades [6]. In addition, long-term trends in rainfall and temperature patterns have been identified in areas with a high proportion of agricultural land. Genomic studies on the locally adapted varieties and subpopulations will provide a potential source of novel alleles which can be exploited in rice breeding programs, such as the new generation of sustainable ‘Green Super Rice’ which are designed to have lower inputs, enhanced nutritional content and suitability for growing on marginal lands [14].

## Conclusions

In this study, we generated a large genome-variation dataset for rice by sequencing 616 accessions from Vietnam and supplementing these with the data obtained for the 3K RGP. Using this resource, we incorporated the Vietnamese rice diversity within the population structure of the Asian cultivated rice. We also identified breeding signatures of selection for the four Japonica and five Indica subpopulations described in this study. The I5 Vietnamese Indica subpopulation showed the highest level of selection, and the elite I1 Indica subpopulation showed the lowest. Overall selection was higher in the Indica subtypes than the Japonica subtypes reflecting the higher diversity of the Indica subtypes. In addition, a GWAS analysis yielded the strongest associations for grain characteristics and weaker associations for a range of characteristics such as panicle length, heading date and leaf width. We used these associations together with published QTLs obtained using a subset of our accessions to give us an insight into traits underlying the regions identified as being under breeding selection. Comparing the Vietnamese subpopulations to the fifteen Asian rice subpopulations identified from the 3K RGP highlighted the I5 subpopulation as a potential source of novel variation as it forms a well-separated cluster. Subpopulation I5 originates from lowland areas such as the Red River Delta and adjacent regions. For the range of phenotypes measured in this study, the I5 subpopulation did not differ phenotypically from the other landraces, which have undergone breeding selection within Vietnam. However, compared to the ‘elite’ I1 subpopulation, I5 accessions have shorter grains, take longer to flower, having lower culm strength, longer culms and leaves. We carried out a comprehensive annotation of the 52 regions selected in I5, which represented 8.1% of the genome and contained 4,576 genes. Candidate genes were identified within these regions as potential breeding targets.

## Materials and Methods

### Sequencing of 616 accessions from Vietnam

We sequenced a total of 616 rice accessions, 612 accessions from Vietnam and three reference accessions, Nipponbare, a temperate Japonica; Azucena, a tropical Japonica; and IR64, an Indica (2 samples). 511 accessions are available from the Vietnamese National Genebank (PRC) at http://csdl.prc.org.vn (Additional file 1: Table S1). All Vietnamese native rice landraces were grown at Dai Dong Experimental Farm (Dai Dong commune, Thach That district, Hanoi, Vietnam) in 2015. The healthy seeds generated from one mature spikelet of the individual plant in each landrace were harvested and dried separately. After that, the selected seeds (35-40 seeds/landrace) were incubated and sown for two weeks to collect leaf samples (30g/sample) for genomic DNA extraction. Total genomic DNA extraction of each rice landrace was made from young leaf tissue using the Qiagen DNeasy kit (Qiagen, Germany). DNA concentration and purity of the samples were measured by the UV-VIS NanoDrop ND-2000 spectrophotometer (Thermo Fisher Scientific) at OD 260/280 nm and OD 260/230 nm wavelengths.

Sequencing was performed by Genomic Services at the Earlham Institute (Norwich, UK). Around 1μg of genomic DNA from each sample was used to construct a sequencing library. For the 36 high coverage samples (prefix: SAM) the Illumina TruSeq DNA protocol was followed, and the samples were sequenced on the HiSeq 2000 for 100 cycles. For the low coverage samples (prefix: LIB), genomic DNA was sheared to 500bp using the Covaris S2 Sonicator (Covaris and Life technologies), and samples were processed using the KAPA high throughout Library Prep Kit (Kapa Biosystems, MA, USA). The ends of the DNA were repaired for the ligation of barcoded adapters. The resulting libraries were quality checked, pooled, and quantified by qPCR. The libraries were sequenced on a HiSeq 2500 instrument following the manufacturer’s instructions.

### Phenotyping

Phenotyping experiments were conducted at the Thach That Experimental Farm of AGI in 2014 and 2015 (Dai Dong commune, Thach That district, Hanoi, Vietnam). The seeds of each rice landrace were incubated in an oven at 45°C for five days to break the seed dormancy. All rice seeds were soaked in tap water for two days and incubated at 35-40°C for four days for germinating. The fully germinated seeds of each rice landrace were directly sown in the paddy field plot (1.5m^2^ in the area). After 15 days of sowing, 24 seedlings of each landrace were carefully transplanted by hand in field plots (2×4m^2^). The fertiliser and pesticide applications were performed following the conventional methods of rice cultivation in Vietnam. The phenotypic and agronomic characteristics were carried out following the method of IRRI [44].

In addition, phenotypic data were available for eleven of the traits in 38 of the 56 genotypes sourced from the 3K-RGP dataset. These eleven traits were included in our analysis because we did not observe a significant difference (p-value > 0.07) between our dataset and the 3K-RGP dataset for the I2 subpopulation (Additional file 1: Table S5).

### Merging the SNP called in the sequenced materials and the complete 3K RGP dataset

Raw sequencing reads were mapped to the Nipponbare reference genome Os-Nipponbare-Reference-IRGSP-1.0 (IRGSP-1.0), using BWA-MEM with default parameters except for “-M-t 8”. Alignments were compressed, sorted and merged using samtools. Picard tools were then used to mark optical and PCR duplicates and add read group information. We used freebayes v1.1.0 for variant calling using default parameters. A total of 21.2 M variants were identified of which 16.4 M were SNPs, and 4.8 M were indels. The resulting VCF file was then filtered for biallelic SNPs with a minimum SNP quality of 30, resulting in 16.0 M variants. PLINK v1.9 was used to convert the VCF into a PLINK BED format. These variants were then combined with the 3K-RGP 29 M biallelic SNPs dataset v1.0 by downloading the PLINK BED files from the “SNP-seek” database (https://snp-seek.irri.org) excluding variants on scaffolds and 26,553 SNPs that were flagged as triallelic upon merging, resulting in 36.9 M SNPs. The SNPs present in both datasets were then extracted and filtered using an identical approach to Wang et al. [15], resulting in 5.9 M SNPs. For that, PLINK v1.9 “--hardy” [45] was used to obtain observed and expected heterozygosity for 100,000 SNPs. We removed SNPs in which heterozygosity exceeds Hardy–Weinberg expectation for a partially inbred species, with inbreeding coefficient (F) estimated as the median value of “1−Hobs/Hexp”, in which Hobs and Hexp are the observed and expected heterozygosity for SNPs where “Hobs/Hexp <1” and the minor allele frequency is >5% and using the cut-off value of 0.479508 for the entire 3,622 samples dataset. A further filtered set of 3.4 M SNPs was obtained by removing SNPs with >20% missing calls and MAF < 1%. Finally, a core set of 361,279 SNPs was obtained with PLINK by LD pruning SNPs with a window size of 10 SNPs, window step of one SNP and r2 threshold of 0.8, followed by another round of LD pruning with a window size of 50 SNPs, window step of one SNP and r2 threshold of 0.8. Samples with more than 50% missing data in this core set were then removed, resulting in dropping seven newly sequenced samples and one genotype from the 3K-RGP dataset.

### Population structure of the combined 3,635 samples

The population structure was analysed using the ADMIXTURE software [46] on the SNP set obtained in the previous section. First, ADMIXTURE was run from K=5 to K=15 in order to compare it with the analysis from IRRI [15, 16]. For each K, ADMIXTURE was then run 50 times with varying random seeds. Each matrix was then annotated using the subpopulation assignment from the 3K-RGP nine subpopulations. Then, up to 10 Q-matrices belonging to the largest cluster were aligned using CLUMPP software [47], these were averaged to produce the final matrix of admixture proportions. Finally, the group membership for each sample was defined by applying a threshold of ≥0.65 to this matrix. Samples with admixture components <0.65 were classified as follows. If the sum of components for subpopulations within the major groups (Ind and Jap) was ≥ 0.65, the samples were classified as Ind-adm or Jap-adm, respectively, and the remaining samples were deemed admixed (admix).

Multi-dimensional scaling analysis was performed using the ‘cmdscale’ function in R, using a distance matrix obtained in R using the Dist function from the amap package [48]. The resulting file was then passed to Curlywhirly [49] and rgl v0.100.19 (https://r-forge.r-project.org/projects/rgl/) for visualisation.

### Recalling the diversity panel with 723 samples

The 616 rice samples were mapped to the Japonica Nipponbare (IRGSP-1.0) reference with BWA-MEM using default parameters, duplicate reads were removed with Picard tools (v1.128) and the bam files were merged using SAMtools v1.5 [50]. Variant calling was completed again on the merged bam file with FreeBayes v1.0.2 [51] separately for each of the 12 chromosomes, but using the option “--min-coverage 10”. Over 6.3 M bi-allelic SNPs with a minimum allele count of ≥3 and quality value above 30 and missing in <50% of samples were obtained with VCFtools v0.1.13 [52]. BAM alignment files to the Nipponbare IRGSP 1.0 reference genome were downloaded from http://snp-seek.irri.org/ [25, 26] for 107 selected samples. Alignment statistics are included in Additional file 1: Table S18. These BAM files were merged and variant calling was similarly completed using FreeBayes v1.0.2 [51] separately for each of the 12 chromosomes using the option --min-coverage 10, and filtered with VCFtools v0.1.13 as before to obtain 6.8 M bi-allelic SNPs with a minimum allele count of ≥3 and quality value above 30 and missing in <50% of samples. The two sets of 6.3 M and 6.8 M SNPs were merged using BCFtools v1.3.1 isec to obtain 4.4 M SNPs which were present in both sets and in at least 70% of samples. These 4.4 M SNPs were then filtered to remove positions which fell outside the expected level of heterozygosity for this dataset, as previously indicated. The resulting estimate of F for the 723 samples was 0.882, so a SNP whose heterozygosity is >5x higher than the most likely value for a given frequency and the dataset’s inbreeding rate will be deemed as having an excessive number of heterozygotes. The cut-off value was 0.591, which resulted in 3.8 M SNPs passing this filter, a scatter plot indicating the SNPs which were kept and removed is shown in Additional file 2: Figure S15. Missing data was imputed in this latest dataset using Beagle v4.1 with default parameters [53]. A comparison using PCA, between the imputed and non-imputed SNP sets showed that imputation did not change the clustering of these 723 samples (Additional file 2: Figure S16). The 3.8M SNPs were subsequently filtered for minimum allele frequency (MAF), linkage disequilibrium (LD pruning or filtering), and distance between polymorphisms (thinning) in different subsets of samples to obtain fourteen sets of SNPs that ranged from 59K to 3.8M SNPs, which were appropriate for the various downstream analysis described below (Additional file 1: Table S19).

### Population structure and diversity analysis for the panel of 672 Vietnamese samples

SNP sets were filtered for MAF 5%, followed by LD filtering using PLINK --indep-pairwise 50 10 0.2, with further thinning if required. We ran STRUCTURE [19] v2.3.5 using the default admixture model parameters; each run consisted of 10,000 burn-in iterations followed by 50,000 data collection iterations. STRUCTURE was run using K=2 for the 616 samples using SNP set 1 (163,393 SNPs). Samples with admixture components <0.75 were classified as admixed, and the remaining samples were classified as Indica or Japonica. STRUCTURE was run varying the assumed number of genetic groups (K) from 3 to 10 with three runs per K value for the 672 Vietnamese samples (SNP set 9 – 80,000 SNPs); from 1 to 8 with ten runs per K value for the 426 Indica subtypes from Vietnam (SNP set 10 - 108,420 SNPs) and the 211 Japonica subtypes from Vietnam (SNP set 11 – 59,815 SNPs). The output files were visualised using the R package POPHELPER v.2.2.7 [54] including the calculation of the number of clusters (K) using the Evanno method [20, 55]. Using the combined-merged clumpp output from POPHELPER, Indica (K=5) and Japonica (K=4) samples were classified into Indica I1 to I5 and Japonica J1 to J4 subpopulations using a threshold of >= 0.6, with the remaining samples being classified as mixed (Im and Jm). The principal component analysis (PCA) was performed using the R package SNPRelate v1.16.0 [55] using method = ‘biallelic’. Nucleotide Diversity (π) was measured for each of the subpopulations with VCFtools v0.1.13 using 100-kbp windows and a step size of 10 kbp.

### Determining the effect of SNPs

The effects of all bi-allelic SNPs (low, medium and high effects) on the genome were determined based on the pre-built release 7.0 annotation from the Rice Genome Annotation Project (http://rice.plantbiology. msu.edu/) using SnpEff [56] release 4.3, with default parameters. The complete set of 3,750,621 SNPs (SNP set 2) which contained on average one variant every 99 bases was annotated. Using sequence ontology terms, the effect of each SNP was classified as described by SnpEff. A summary of the SNP effect analysis is available in Additional file 1: Table S20.

### Genome-wide association analysis

Three independent analyses were conducted using the full panel (672 samples, 361,191 SNPs), the Indica subpanel (426 samples, 334,935 SNPs) and the Japonica subpanel (211 samples, 122,881 SNPs), SNP sets 12, 13 and 14 respectively (Additional file 1: Table S19). The GWAS analysis was performed by employing the R package Genome Association and Prediction Integrated Tool (GAPIT) version 3.0 [57, 58]. The covariate matrix was generated in STRUCTURE. We used the combined-merged output from POPHELPER for the full panel (K=8), the Indica subpanel (K=5) and the Japonica subpanel (K=4). The covariate matrix and the kinship calculated in GAPIT were included in the GWAS model to control for false positives. The SUPER (Settlement of MLM Under Progressively Exclusive Relationship [59] method integrated into GAPIT, designed to increase the statistical power, was used to perform the association mapping analysis. The SUPER method was implemented in GAPIT by setting the parameter of “*sangwich.top*” and “*sangwich.bottom*” to C*MLM* and *SUPER*, respectively. A quantile-quantile (Q–Q) plot was used to check if the model was correctly accounting for both confounding variables. Associations held by peaks with −log_10_ (p-value) ≥ 8.0 were used to declare the significant associations. The Genes lying within the QTL regions were extracted and subjected to enrichment analysis using PhytoMine implemented within Phytozome [21] https://phytozome.jgi.doe.gov/ for Gene Ontology, Protein Domain and Pathway enrichment using a max p-value of 0.05 with Bonferroni correction.

### Identification of selective sweeps using XP-CLR

Selective sweeps across the genome were identified using XP-CLR, a method based on modelling the likelihood of multilocus allele frequency differentiation between two populations. An updated version (https://github.com/hardingnj/xpclr) of the code described by Chen et al. [22] was used to scan for regions of selection. We used 100-kbp sliding windows with a step size 10 kbp and the default of no more than 200 SNPs per window. XP-CLR was run between the five Indica subpopulations and the four Japonica subpopulations. Selected regions were extracted using the XP-CLR score for each 100-kb window as follows-200 kbp centromeric regions were removed, and the mean XP-CLR score and 99th percentile were calculated for comparisons between subpopulation (e.g. I5 vs I1, I2, I3, I4) and the mean of these values was used to define the cut-off level for selection in that population as shown in Additional file 1: Table S10. 100-kbp regions with an XP-CLR score higher than the cut-off were extracted and merged using BEDTOOLS v2.26.0 [60] specifying a maximum distance between regions of 100 kbp. Regions shorter than 80 kbp were then removed to give a final set of putatively selected regions for each comparison. Overlapping regions selected in comparison with at least two subpopulations for Japonica or three for Indica were then merged to obtain a final set of selected regions for each subpopulation. BEDTOOLS map was used for finding any overlap of selected regions with QTLs. The Genes lying within the selected regions were extracted and subjected to enrichment analysis as before.

### Calculating F_ST_

We calculated F_ST_ per SNP between the 43 samples in the I5 subpopulation and the 190 samples in the I2, I3 and I4 subpopulations with VCFtools using the “weir-fst-pop” option which calculates F_ST_ according to the method of Weir and Cockerham [61]. Sites which are homozygous between these populations were removed, and negative values were changed to zero. The mean F_ST_ was calculated per gene and per specified region.

## Supporting information

Table S20. Summary count of SNPs with effects on the genome (using MSU7 annotation)

Table S19. List of 14 SNP-sets used for analysis

Table S18. List of 107 IRRI rice samples from Vietnam in the 3K RGP

Table S17. List of 21 genes related to salt tolerance selected in the I5 subpopulation

Table S16. List of genes selected in I5 subpopulation

Table S13. Annotation of I5 selected regions using PhytoMine

Table S12. List of genes selected in each region for Indica I5 subpopulation

Table S11. List of genes selected in each subpopulation

Table S10. Summary of XP-CLR comparisons for the Indica and Japonica subpopulations

Table S9. Annotation of QTLs using PhytoMine

Table S8. Gene lists for the 21 QTLs

Table S6. Diversity of each subpopulation

Table S7. GWAS results; List of the 21 QTLs and the positions of the individual QTLs for each panel.

Table S5. Phenotype statistics (mean and coefficient of variation) and population comparisons

Table S4. Phenotyping abbreviations and details

Table S3. Phenotypic measurements for 20 traits for 672 samples

Table S2. Name and details of 3,635 rice varieties

Table S1. Name and details of 672 Vietnamese rice varieties

Table S15. Overlap of the selected regions in Japonica subpopulations with QTL found in Vietnamese rice datasets

Table S14. Overlap of the selected regions in Indica subpopulations with QTL found in Vietnamese rice datasets

## Acknowledgements

We thank Professor Giles Oldroyd for his contributions to the conception of this project. We are grateful for the support from Dr. Nelzo Ereful, and Matt Heaton during outreach activities in Vietnam, and Dr. Luca Venturi, Dr. Ricardo Ramirez Gonzalez, Dr. Graham Etherington for their support during summer training activities in the UK, and Dr. Chris Watkins, Dr. Helen Chapman and the Genomics Pipelines team at the Earlham Institute for the sequencing support.

## Availability of data

All sequence data used in this manuscript have been deposited as study PRJEB36631 in the European Nucleotide Archive.

## Author contributions

TDK, KHT, AH, SD, LHH, MC and JDV designed and conceived the research. TDK, KHT, TDD, NTPD, NTK, DTTH, NTD, KTD, CNP, TTT, NTT, HDT, NTT, HTG, TKN, CDT, SVL, LTN, NVG and LHH performed the phenotyping and laboratory experiments. JH and BS performed the data analysis with assistance from TDD, NTPD, DTTH, NTD, KTD, NTT, LTN, TDX, MC and JDV. JH, BS and JDV wrote the paper. All authors read and approved the final manuscript.

## Ethics approval and consent to participate

Not applicable.

## Consent for publication

Not applicable.

## Supplementary tables

**Table S1. Name and details of 672 rice varieties.** Detailing read number, mapping statistics, Vietnamese National Genebank number, local name, location, characteristic, subtype and subpopulation.

**Table S2. Name and details of 3,635 rice varieties.** Detailing the new subpopulation and PCO analysis.

**Table S3. Phenotypic measurements for 20 traits for 672 samples.** Detailing individual measurements for each sample, description of phenotypes, statistics for all samples and individually for the Indica and Japonica subtypes. Phenotypes are available for around 75% of the samples.

**Table S4. Phenotype abbreviations and details.**

**Table S5. Phenotype statistics (mean and coefficient of variation) and population comparisons.**

**Table S6.** Diversity (π) of each subpopulation.

**Table S7.** GWAS results. List of the 21 QTL and the positions of the individual QTLs for each panel.

**Table S8. Gene lists for the 21 QTL.**

**Table S9. Annotation of QTL using PhytoMine.** Enrichment analysis for protein domain, Meta-Cyc pathway and Geno Ontology using PhytoMine.

**Table S10. Summary of XP-CLR comparisons for the Indica and Japonica subpopulations.** Detailing the XP-CLR mean, cut off and number of regions for each comparison.

**Table S11. List of genes selected in each subpopulation.**

**Table S12. List of genes selected in each region for Indica I5 subpopulation.**

**Table S13. Annotation of I5 selected regions using PhytoMine.** List of all genes within each of the 52 selected regions and the results of PhytoMine Enrichment analysis for protein domain, Meta-Cyc pathway and Geno Ontology using PhytoMine.

**Table S14. Overlap of the selected regions in Indica subpopulations with QTL found in Vietnamese rice datasets.**

**Table S15. Overlap of the selected regions in Japonica subpopulations with QTL found in Vietnamese rice datasets.**

**Table S16. List of genes selected in I5 subpopulation.** Detailing MSU and RAP gene ID, annotation and enrichment in Phytomine, high impact SNPs and mean F^st^

**Table S17. List of 21 genes related to salt tolerance selected in the I5 subpopulation.**

**Table S18. List of 107 IRRI rice samples.** Detailing IRRI accession, country on origin, K9 and K15 group and Vietnamese subpopulation.

**Table S19. List of 14 SNP sets used for analysis.** Detailing filtering parameters, sample and SNP numbers for each SNP set.

**Table S20. Summary count of SNPs with effects on the genome.** Detailing SnpEff annotation of the full set of 3,750,621 SNPs using the *Oryza sativa* MSU release 7 rice annotation. Six tables detailing number of effects by impact, functional class, type, region, base changes and Ts/Tv ratio.

## Supplementary Figures

**Figure S1. Analysis of STRUCTURE output using the Evanno method.**

Evanno Plots output from Pophelper for 672 Vietnamese samples, 426 Indica samples and 211 Japonica samples.

**Figure S2. Mapping rate (%properly paired) for Japonica and Indica subpopulations.**

**Figure S3. Principal coordinate analysis (PCO) of the 3,635 Asian cultivated rice genomes.** Plots are coloured by the subpopulations **a** K9_new, **b** K15_new. The first component represents the separation between the Indica and Japonica lines. The second components show the separation of cAus and to a lesser extent cBas while the third and fourth components represent the separation within Japonica and Indica respectively. Note for (a) we display the first 3 components and for (b) we display components 1, 2 and 4.

**Figure S4. Comparison between K15_3KRGP, K15_new and Vietnamese subpopulations. a** Comparison between K15_3KRGP and K15_new using 3023 samples. **b** Comparison between K15_new and Vietnamese subpopulations using 668 samples (overlap of 56 samples from Vietnam with a). **c** Percentage of K15_new subpopulations from Vietnam. Arrow are shown for subpopulations which consist of > 50% of samples from Vietnam. Diagram generated using http://sankeymatic.com/

**Figure S5. PCO analysis of 1605 Indica samples**. Omitting the samples classified as XI-adm and Ind-adm outside Vietnam for clarity. Plot coloured by **a** K15_3KRGP, **b** K15_new including Vietnamese samples, **c** Five Vietnamese Indica subpopulations. The ellipses show the 95% confidence interval. X = PC1, Y=PC4, Z=PC5. Figure generated using rgl https://r-forge.r-project.org/projects/rgl/

**Figure S6. PCO analysis of 982 Japonica samples.** Omitting the samples classified as GJ-adm and Jap-adm outside Vietnam for clarity. Plot coloured by **a** K15_3KRGP, **b** K15_new including Vietnamese samples, **c** Four Vietnamese Japonica subpopulations. The ellipses show the 95% confidence interval. X = PC3, Y=PC4, Z=PC5. Figure generated using rgl https://r-forge.r-project.org/projects/rgl/

**Figure S7. Admixture components of the Indica I3, I4 and I5 subpopulations.**

**Figure S8. PCA analysis of Indica and Japonica Vietnamese subpopulations including 51 genotypes from outside Vietnam. a** PCA analysis of 445 accessions using the top two components to separate the five Indica subpopulations. The ellipses show the 95% confidence interval. **b** PCA analysis of 233 accessions using the top two components to separate the four Japonica subpopulations. The ellipses show the 95% confidence interval.

**Figure S9. Correlation between the 20 phenotypes.**

**Figure S10. Correlation between Indica and Japonica for the 13 phenotypes used for GWAS.** The figure was created using “ggpairs” package in R.

**Figure S11. Correlation between Indica I1 and I5 subpopulations for the 13 phenotypes used for GWAS.** The figure was created using “ggpairs” package in R.

**Figure S12. Boxplots showing the Phenotypic distribution per subpopulation for Culm Length, Grain Length, Grain Width and Heading Date.**

**Figure S13. Indica subpopulation diversity.**

Diversity (π) plotted along the 12 rice chromosomes in sliding 100kb windows.

**Figure S14. Japonica subpopulation diversity.**

Diversity (π) plotted along the 12 rice chromosomes in sliding 100kb windows.

**Figure S15. SNP filtering for heterozygosity**

Proportion of heterozygous calls versus allele frequency. Each dot represents a SNP from a random sample of 100,000 SNPs. The points have an opacity of 5% to highlight regions of higher point density. The bulk of the SNPs lie on the Hardy-Weinberg equilibrium curve scaled by a factor of around 0.118, which implies a Wright’s inbreeding coefficient of F=0.882. The SNPS have been filtered using cut off of 0.592 (5*(1-F)), the corresponding SNPs which are kept and removed are shown on the plot.

**Figure S16. PCA analysis of 723 samples before and after imputation.**

Comparing the 2,690,005 not imputed SNP set 3 to the 2,665,825 imputed SNP set 4 Both SNP set were filtered for 5% MAF. Using PC1 and PC2 to separate the Japonica subpopulations. Using PC3 and PC4 to separate the Indica subpopulations.

**Figure S17.** GWAS Manhattan and qq plots for the full panel and Indica and Japonica subpanels for Grain Length, Grain Width, Grain length-to-width ratio, Heading Date, Culm Strength, Leaf Length and Leaf Width.

**Figure S18.** GWAS Manhattan and qq plots for the full panel and Indica and Japonica subpanels for Leaf Pubescence, Culm Number, Diameter Internode, Culm Length, Panicle Length and Floret Pubescence

**Figure S19. Chromosome plots of regions selected in each Indica subpopulation showing the regions selected against each individual subpopulation and the shaded final selected regions which were selected against three subpopulations.** a) 44 regions selected in I1, b) 41 regions selected in I2, c) 42 regions selected in I3, d) 38 regions selected in I4, e) 52 regions selected in I5

**Figure S20. Chromosome plots of regions selected in each Japonica subpopulation showing the regions selected against each individual subpopulation and the shaded final selected regions which were selected against two subpopulations.** a) 28 regions selected in J1, b) 23 regions selected in J2, c) 24 regions selected in J3, d) 25 regions selected in J4

**Figure S21. Allele Plots showing the “*High* impact” SNP position within candidate genes.**

