## Supplementary material for "Evidence of selection, adaptation and untapped diversity in Vietnamese rice landraces": SUPPL FIGURES S1 TO S16

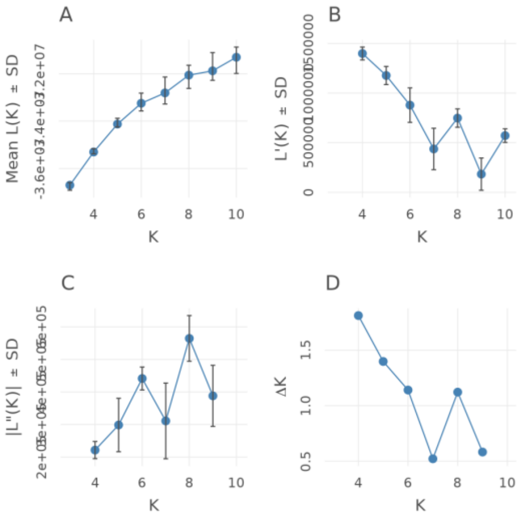


672 Vietnamese varieties

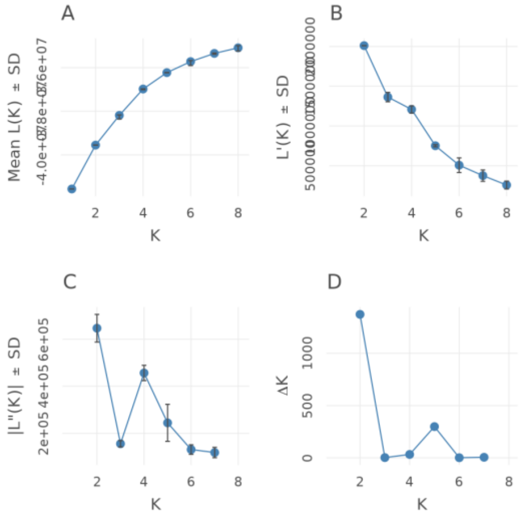


426 Indica subtypes

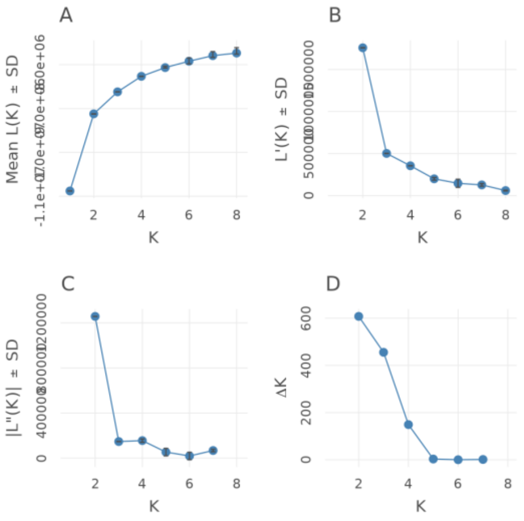


211 Japonica subtypes

**Figure S1 Analysis of STRUCTURE output using the Evanno method.**
Evanno Plots output from Pophelper for 672 Vietnamese samples, 426 Indica samples and 211 Japonica samples.

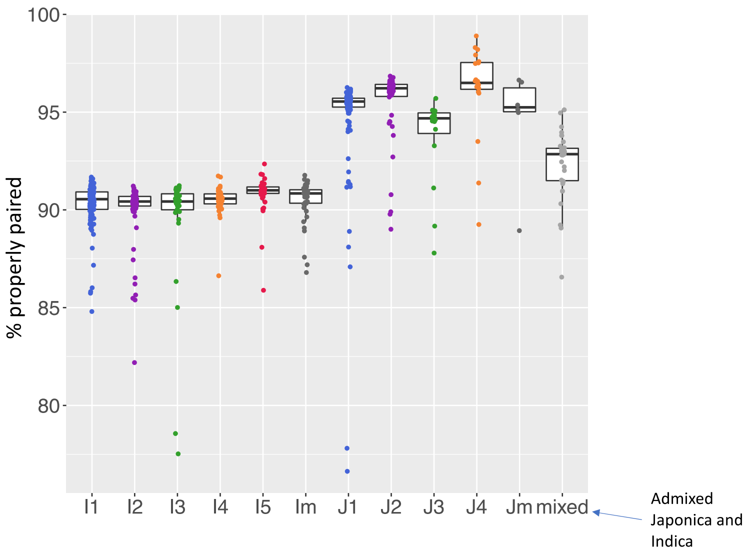


**Figure S2. Mapping rate (% properly paired) for Japonica and Indica subpopulations.**

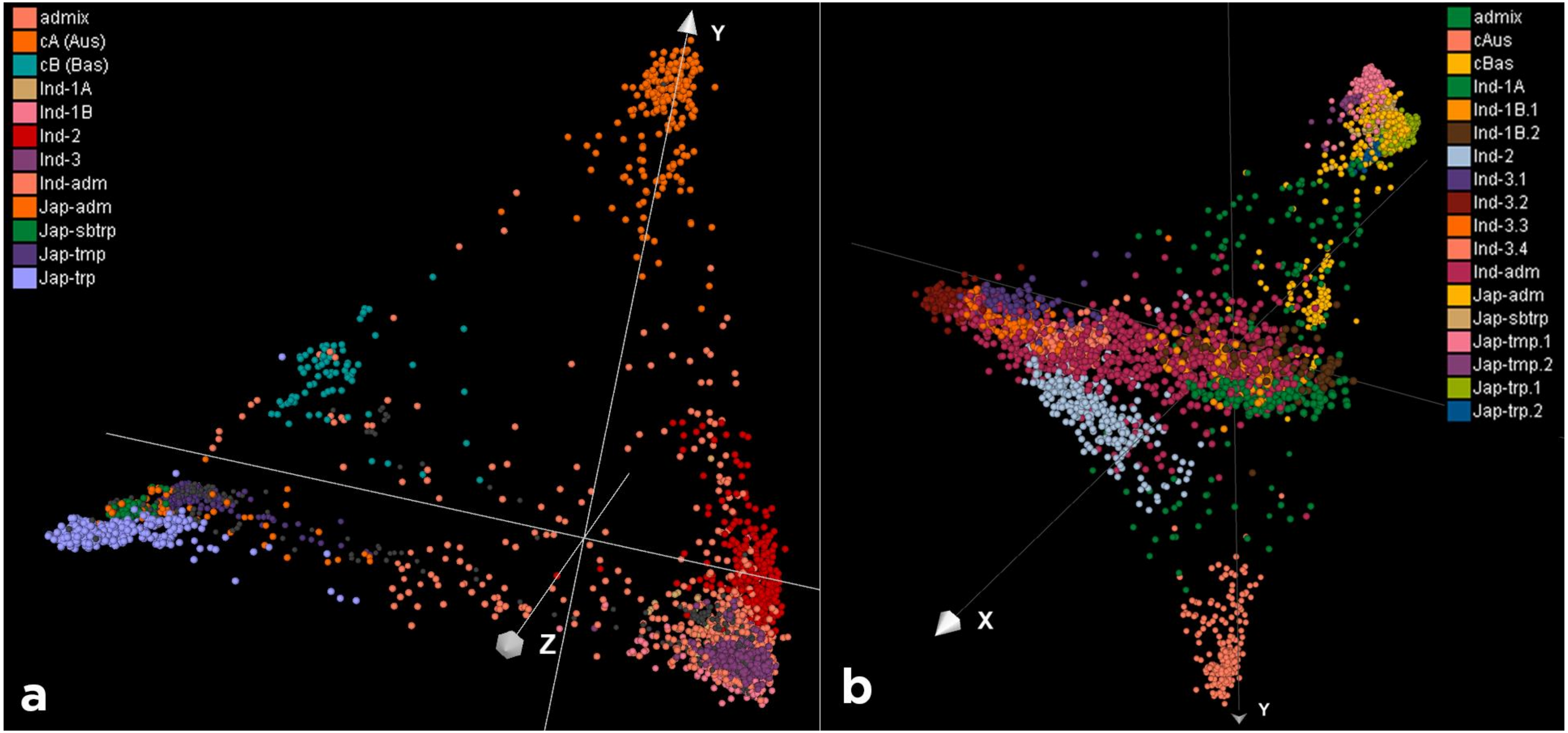


**Figure S3**. **Principal coordinate analysis (PCO) of the 3,635 Asian cultivated rice genomes**. Plots are coloured by the subpopulations **a** K9_new, **b** K15_new. The first component represents the separation between the Indica and Japonica lines. The second components show the separation of cAus and to a lesser extent cBas while the third and fourth components represent the separation within Japonica and Indica respectively. Note for (a) we display the first 3 components and for (b) we display components 1, 2 and 4.

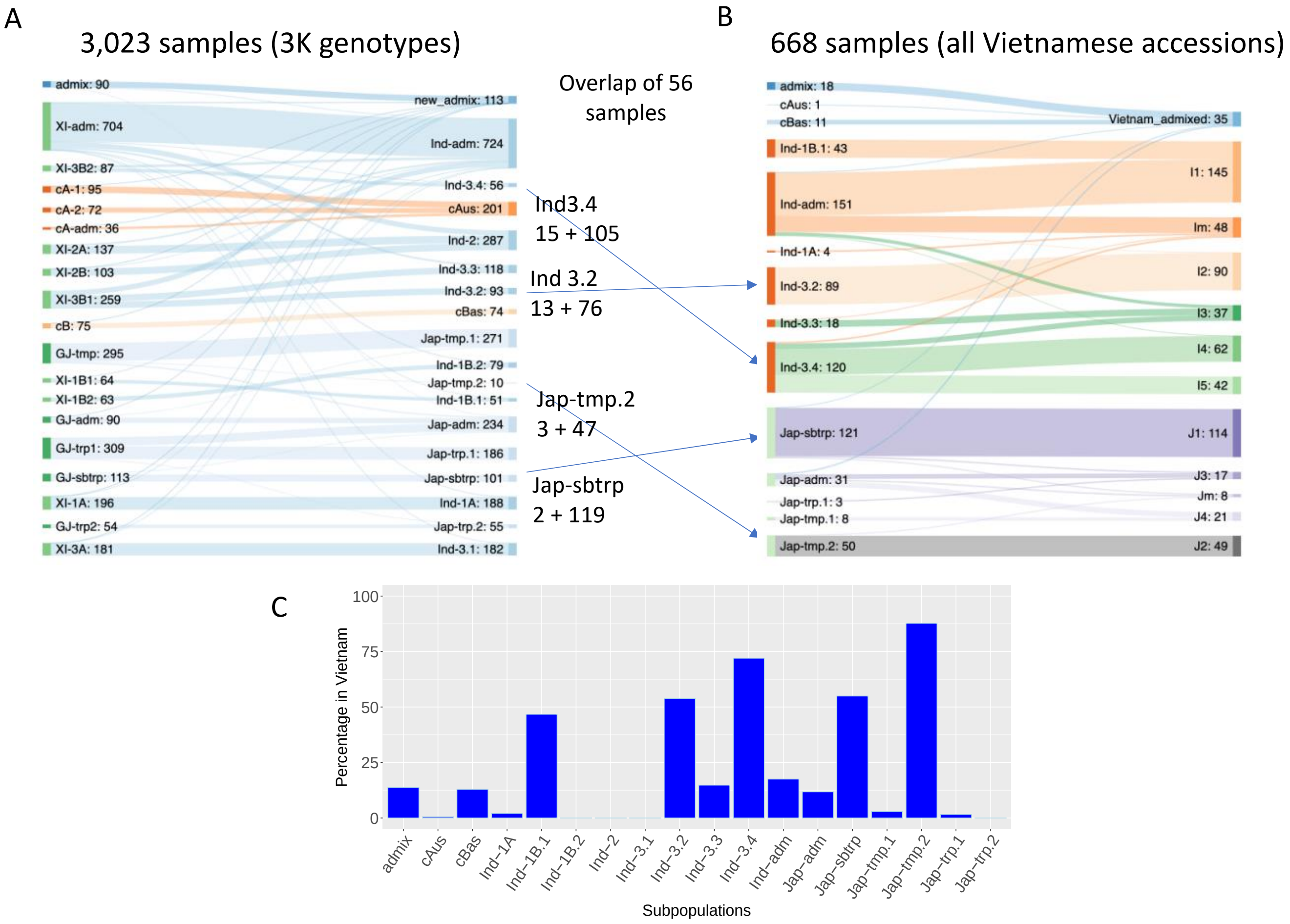


**Figure S4. Comparison between K15_IRRI, K15_new and Vietnamese subpopulations.**

**a** Comparison between K15_IRRI and K15_new using 3023 samples. **b** Comparison between K15_new and Vietnamese subpopulations using 668 samples (overlap of 56 samples from Vietnam with a). **c** Percentage of K15_new subpopulations from Vietnam

Arrow are shown for subpopulations which consist of > 50% of samples from Vietnam, showing the numbers of 3K RGP genotypes and newly sequenced accessions respectively

Diagram generated using http://sankeymatic.com/

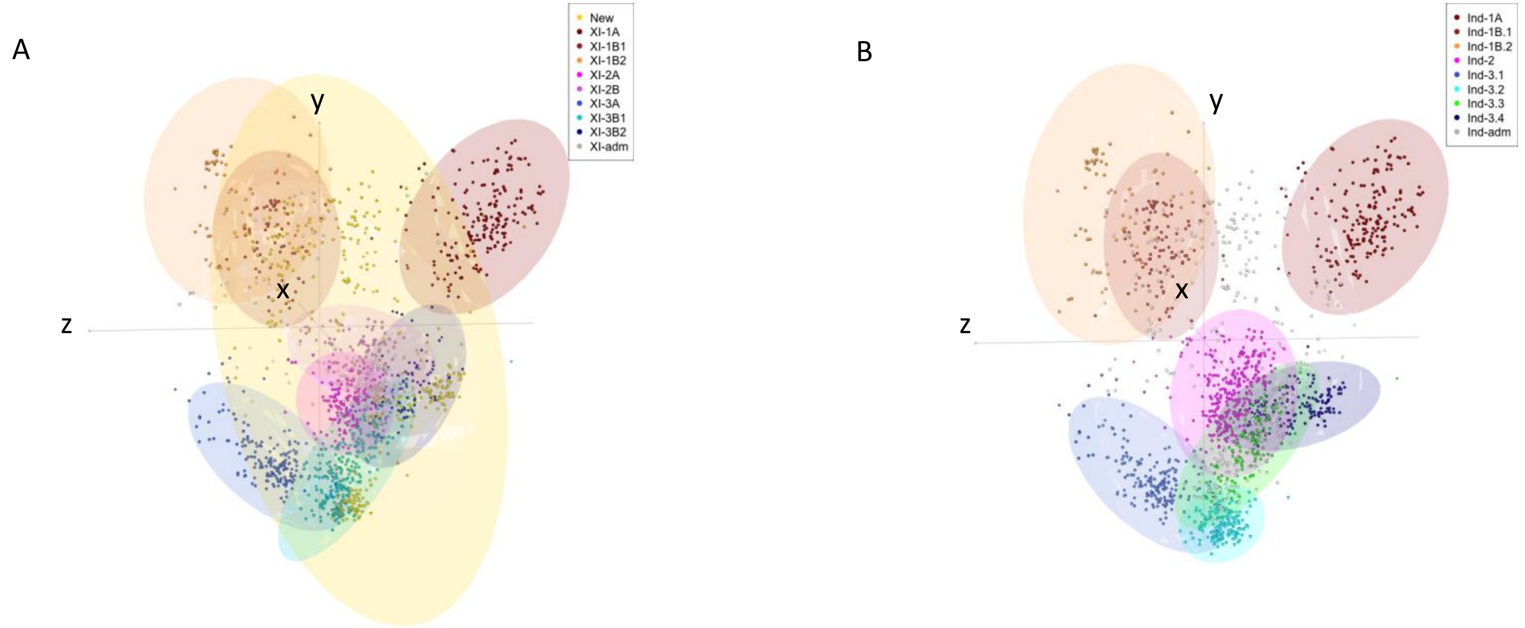


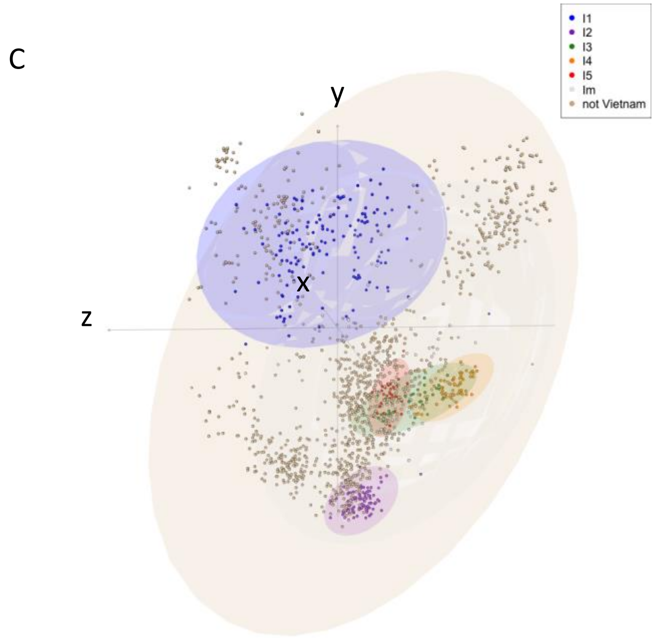


**Figure S5. PCO analysis of 1605 Indica samples.**
Omitting the samples classified as XI-adm and Ind-adm outside Vietnam for clarity. Plot coloured by **a** K15_3KRGP, **b** K15_new including Vietnamese samples, **c** Five Vietnamese Indica subpopulations.
The ellipses show the 95% confidence interval.
X = PC1, Y=PC4, Z=PC5.
Figure generated using rgl https://r-forge.r-project.org/projects/rgl/

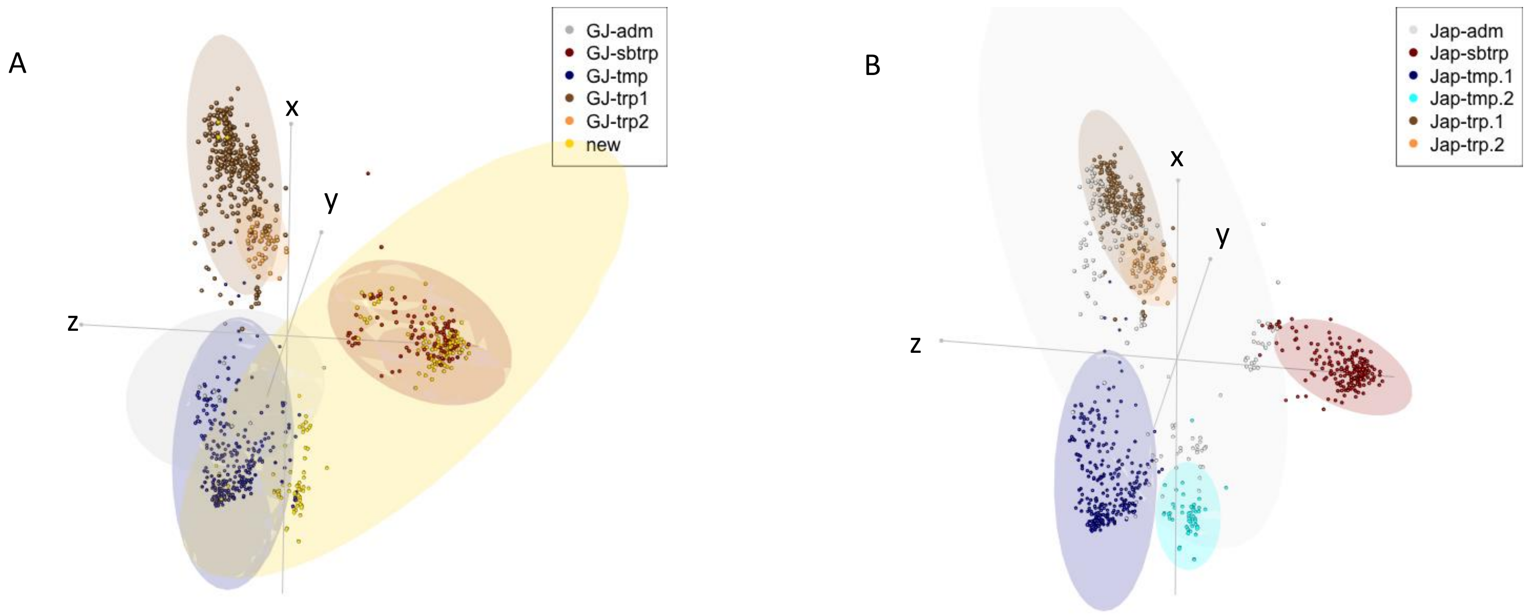


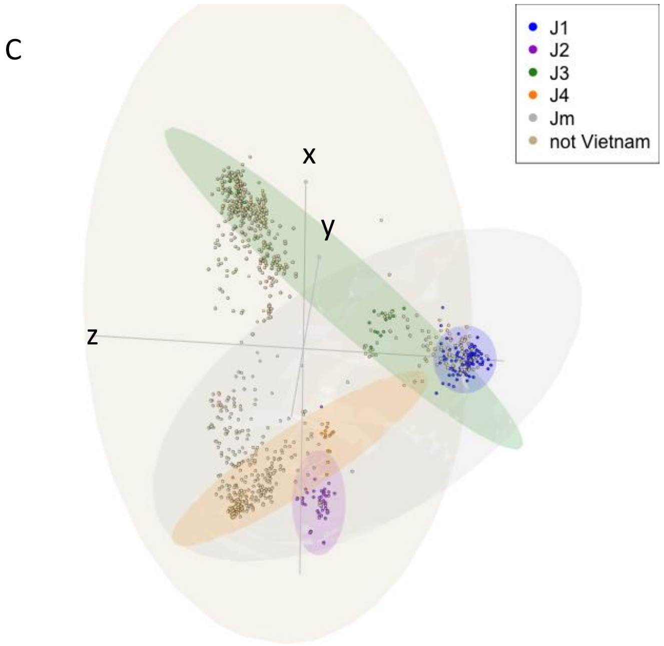


**Figure S6. PCO analysis of 982 Japonica samples.**
Omitting the samples classified as GJ-adm and Jap-adm outside Vietnam for clarity. Plot coloured by
**a** K15_3KRGP, **b** K15_new including Vietnamese samples,
**c** Four Vietnamese Japonica subpopulations.
The ellipses show the 95% confidence interval.
X = PC3, Y=PC4, Z=PC5.
Figure generated using rgl https://r-forge.r-project.org/projects/rgl/

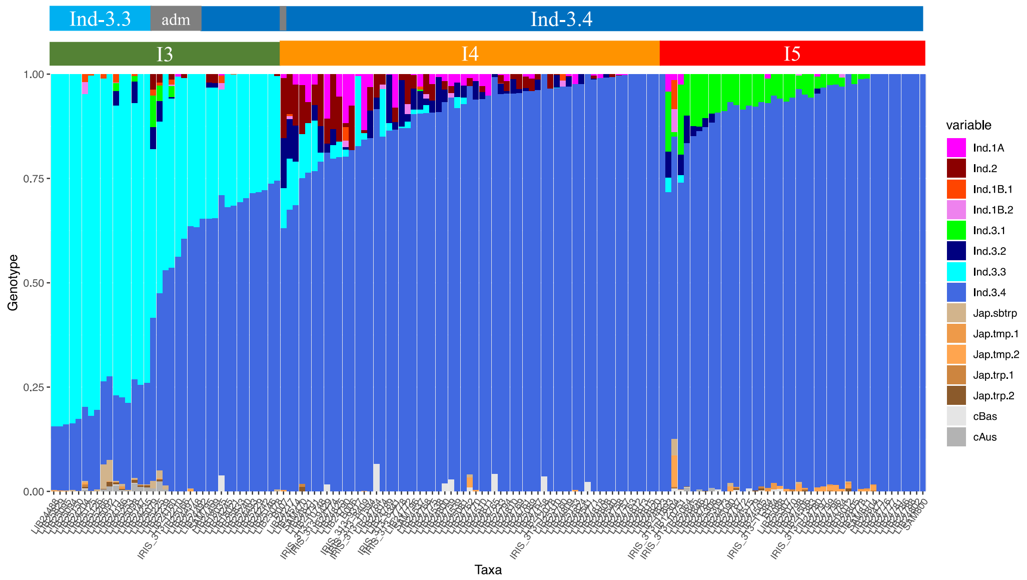


**Figure S7. Admixture components of the Indica I3, I4 and I5 subpopulations.**

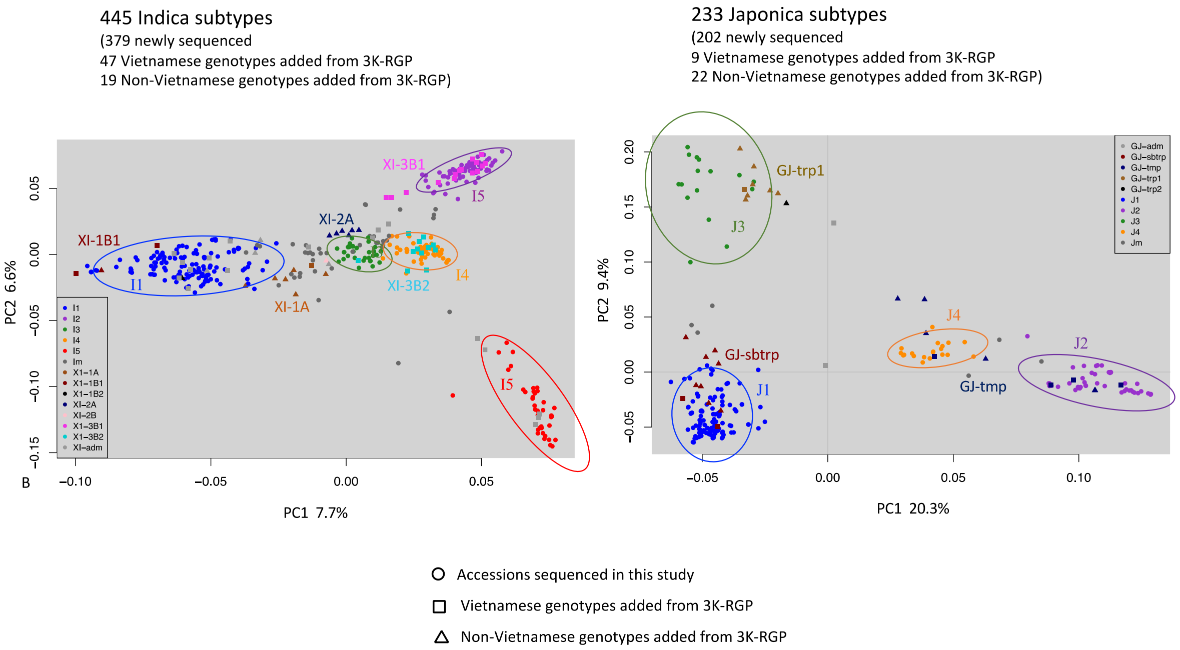


**Figure S8. PCA analysis of Indica and Japonica Vietnamese subpopulations including 51 genotypes from outside Vietnam.**

**a** PCA analysis of 445 accessions using the top two components to separate the five Indica subpopulations. The ellipses show the 95% confidence interval. **b** PCA analysis of 233 accessions using the top two components to separate the four Japonica subpopulations. The ellipses show the 95% confidence interval.

| | |
|---|---|
| LmPb | Floret_Pubescence |
| LmPc | Floret_Colour |
| An | Awning |
| PnT | Panicle_Type |
| PnL | Panicle_Length |
| CmL | Culm_Length |
| DBI | Diameter_Internode |
| CmA | Culm_Angle |
| CmN | Culm_Number |
| FLA | Flag_Leaf_Angle |
| LA | Leaf_Angle |
| LBP | Leaf_Pubescence |
| LW | Leaf_Width |
| LL | Leaf_Length |
| PnEx | Panicle_Exsertion |
| CmS | Culm_Strength |
| HD | Heading_Date |
| GL_GW | GL_GW_ratio |
| GrW | Grain_Width |
| GrL | Grain_Length |

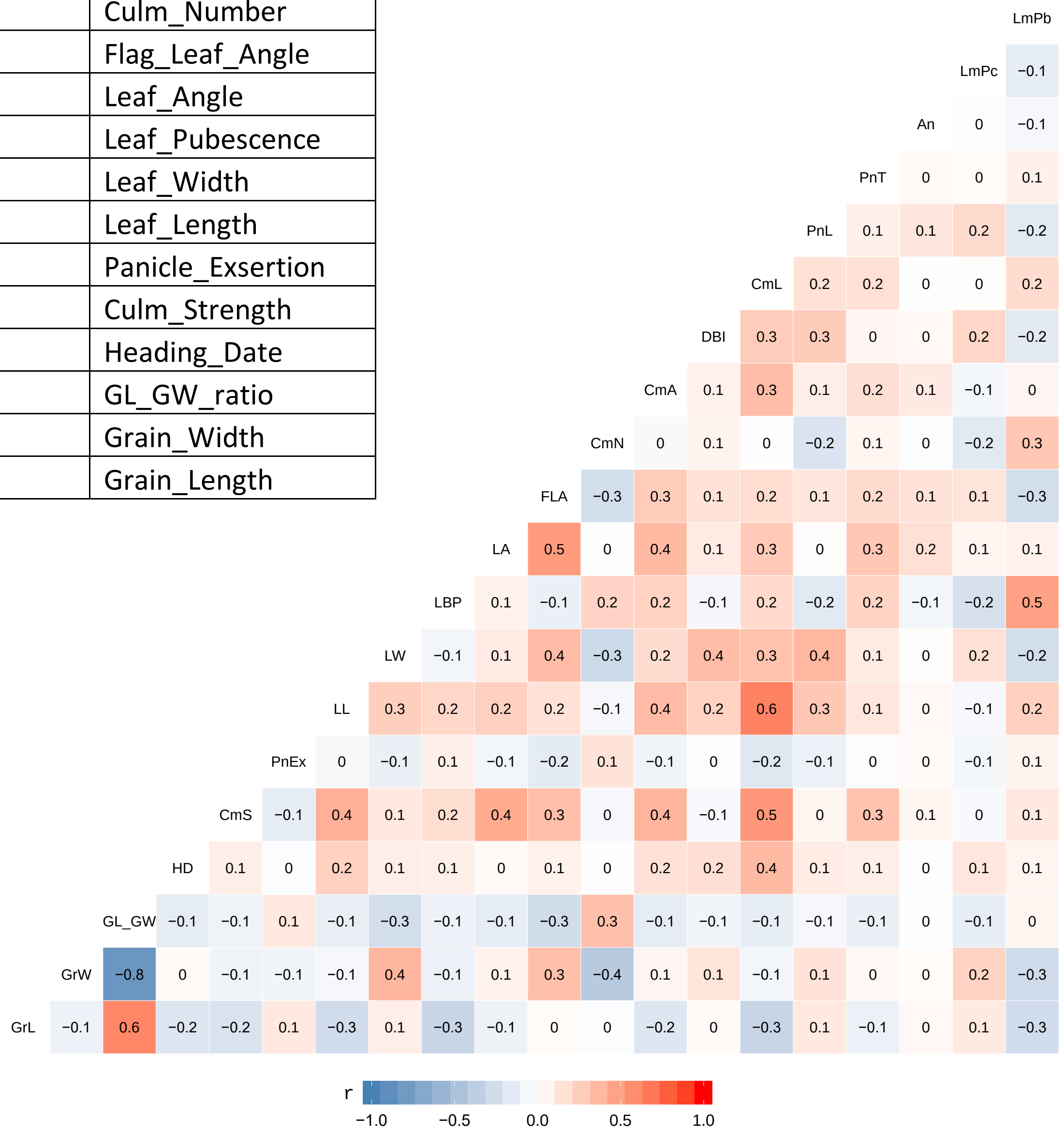


**Figure S9. Correlation between the 20 phenotypes.**

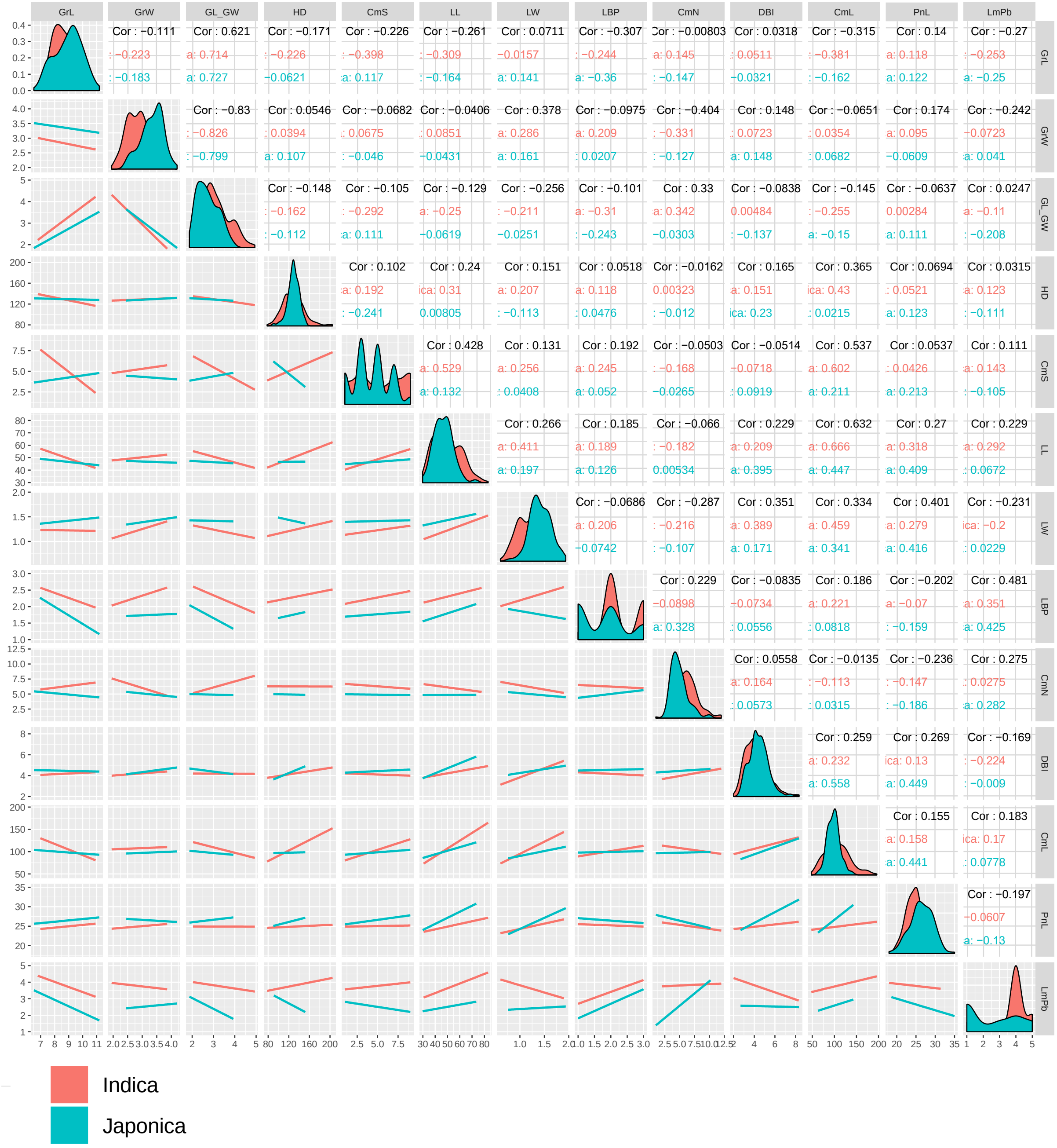

Figure S10. Correlation between Indica and Japonica for the 13 phenotypes used for GWAS

**Figure S10. Correlation between Indica and Japonica for the 13 phenotypes used for GWAS**
The figure was created using “ggpairs” package in R.

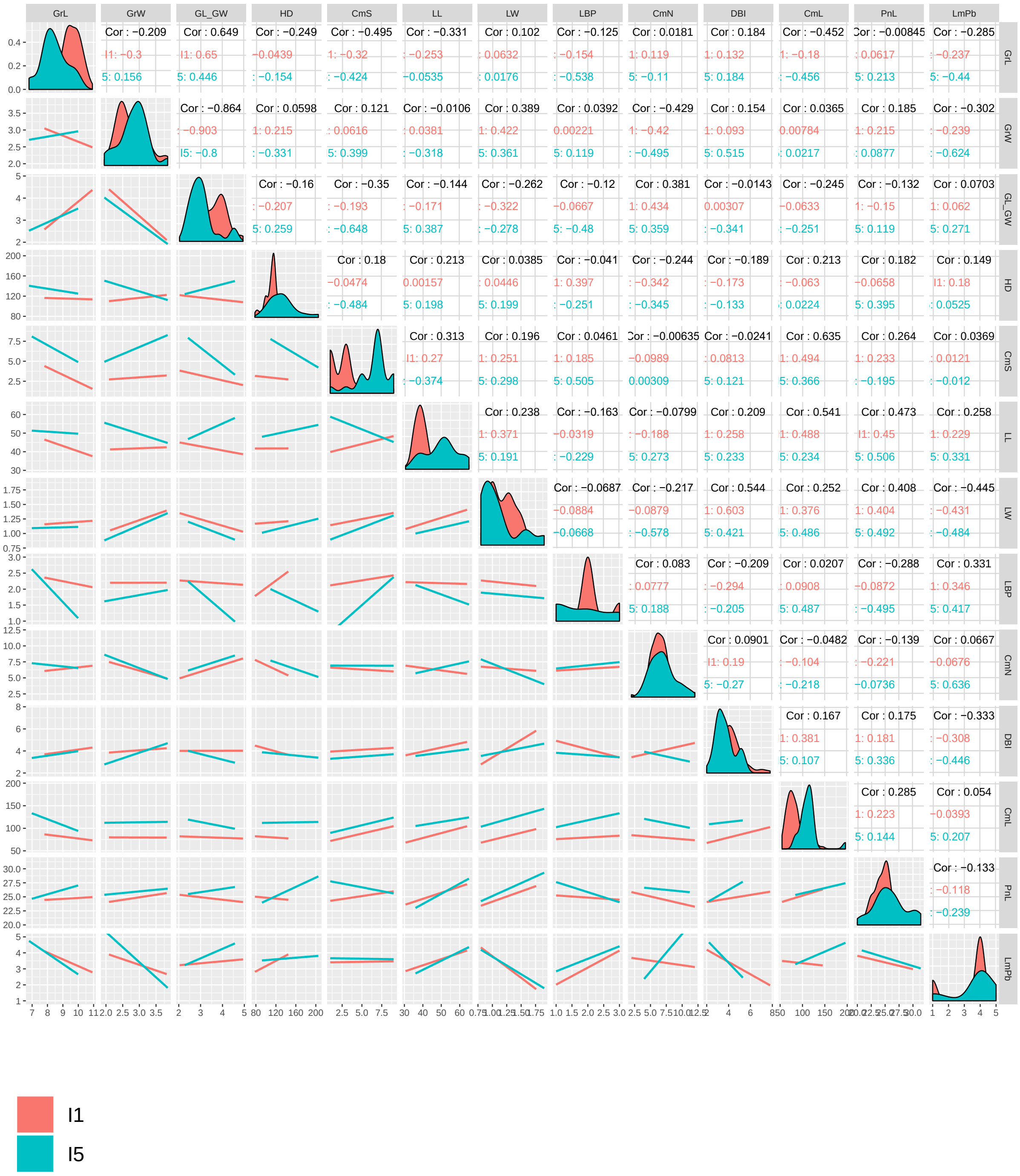


**Figure S11. Correlation between Indica I1 and I5 subpopulations for the 13 phenotypes used for GWAS.**

The figure was created using “ggpairs” package in R.

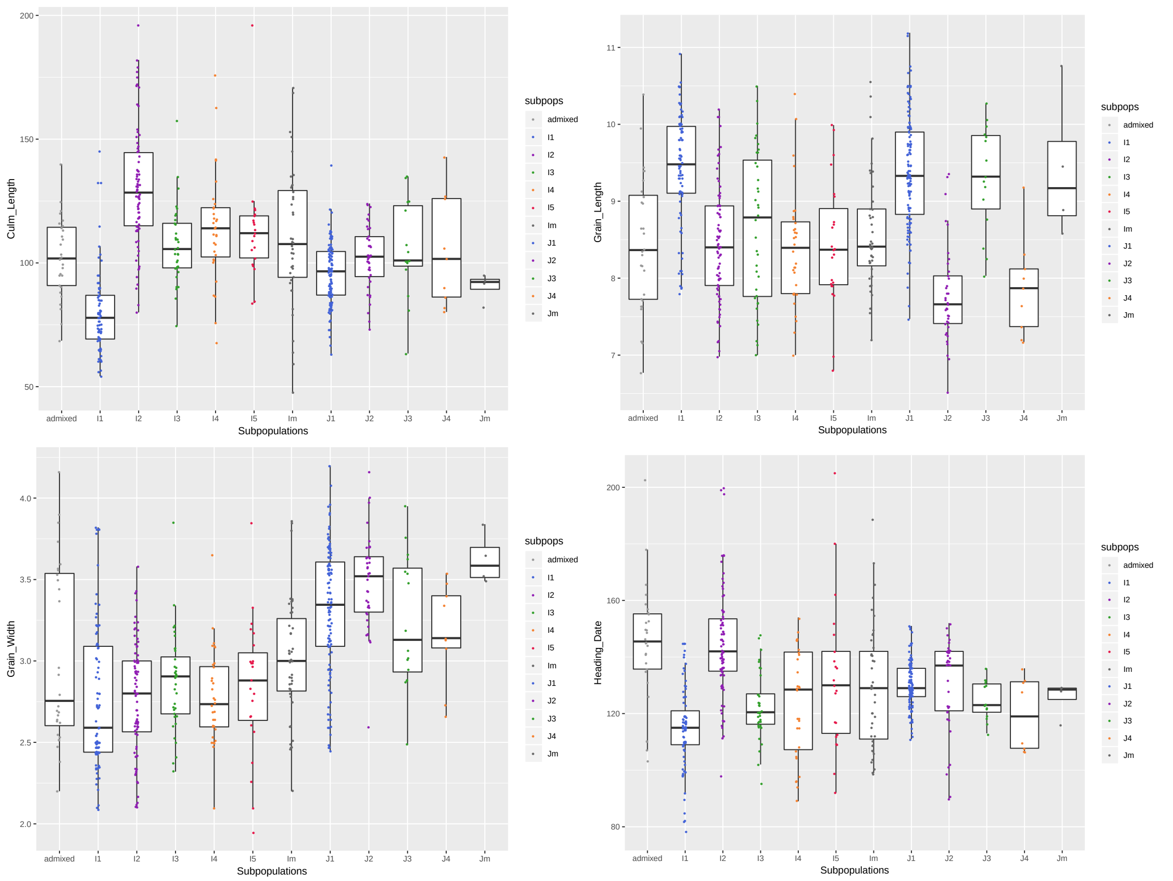


**Figure S12. Boxplots showing the Phenotypic distribution per subpopulation for Culm Length, Grain Length, Grain Width and Heading Date.**

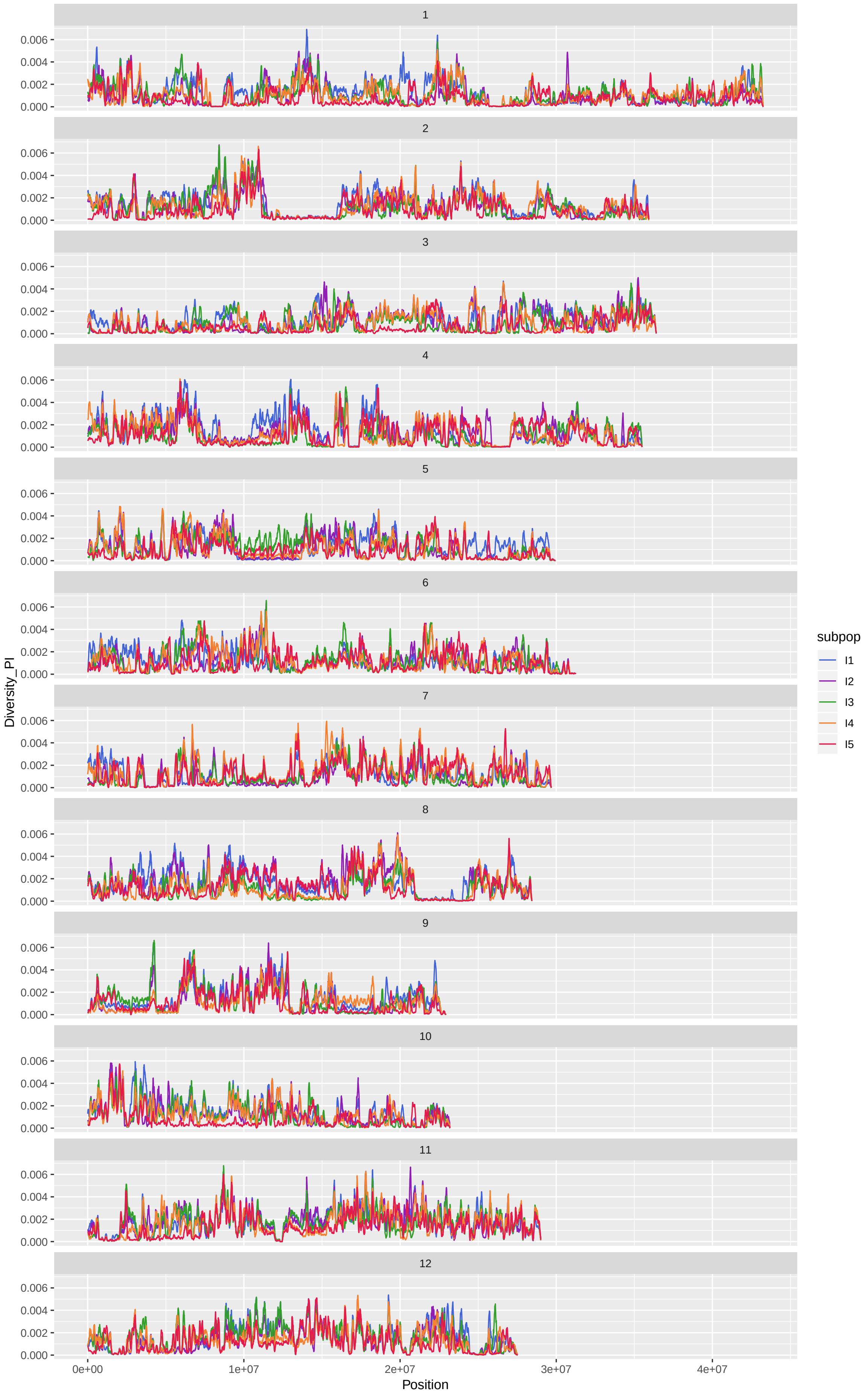


**Figure S13. Indica subpopulation diversity.**

Diversity (π) plotted along the 12 rice chromosomes in sliding 100kb windows.

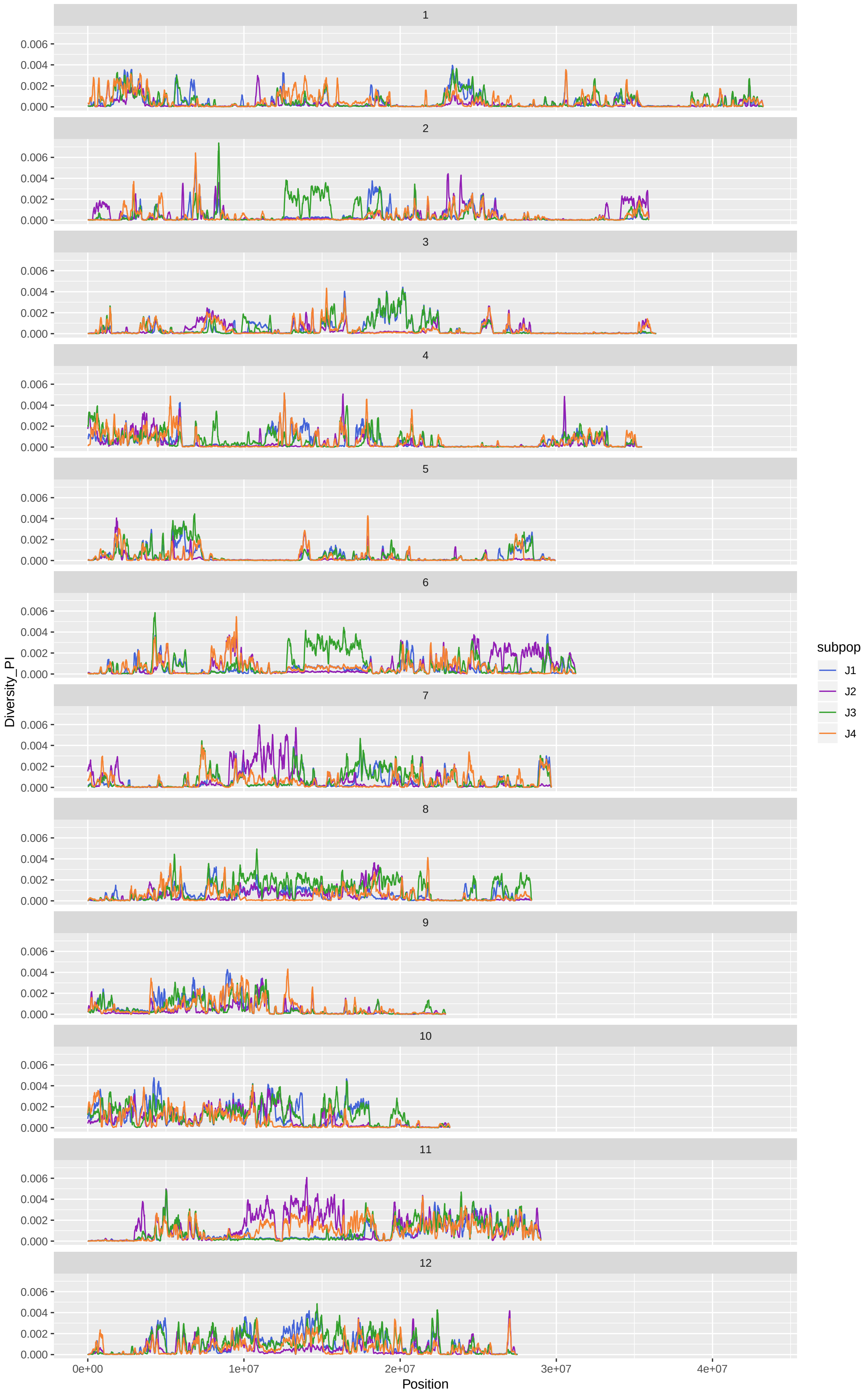


**Figure S14. Japonica subpopulation diversity.**

Diversity ($\pi$) plotted along the 12 rice chromosomes in sliding 100kb windows.

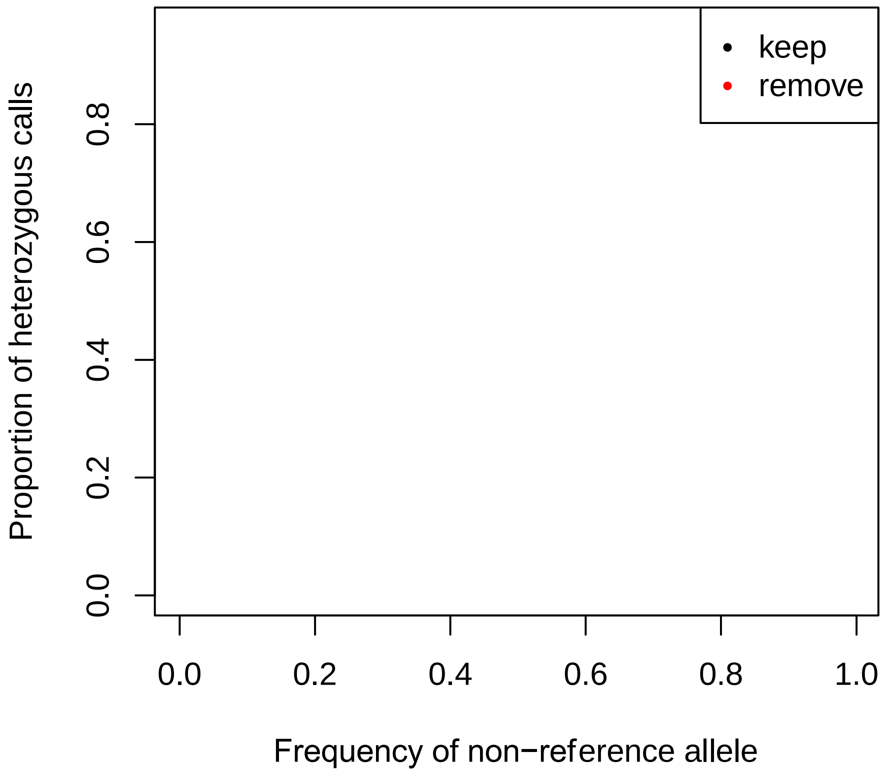


**Figure S15. SNP filtering for heterozygosity**
Proportion of heterozygous calls versus allele frequency. Each dot represents a SNP from a random sample of 100,000 SNPs. The points have an opacity of 5% to highlight regions of higher point density. The bulk of the SNPs lie on the Hardy-Weinberg equilibrium curve scaled by a factor of around 0.118, which implies a Wright's inbreeding coefficient of F=0.882. The SNPS have been filtered using cut off of 0.592 (5*(1-F)), the corresponding SNPs which are kept and removed are shown on the plot.

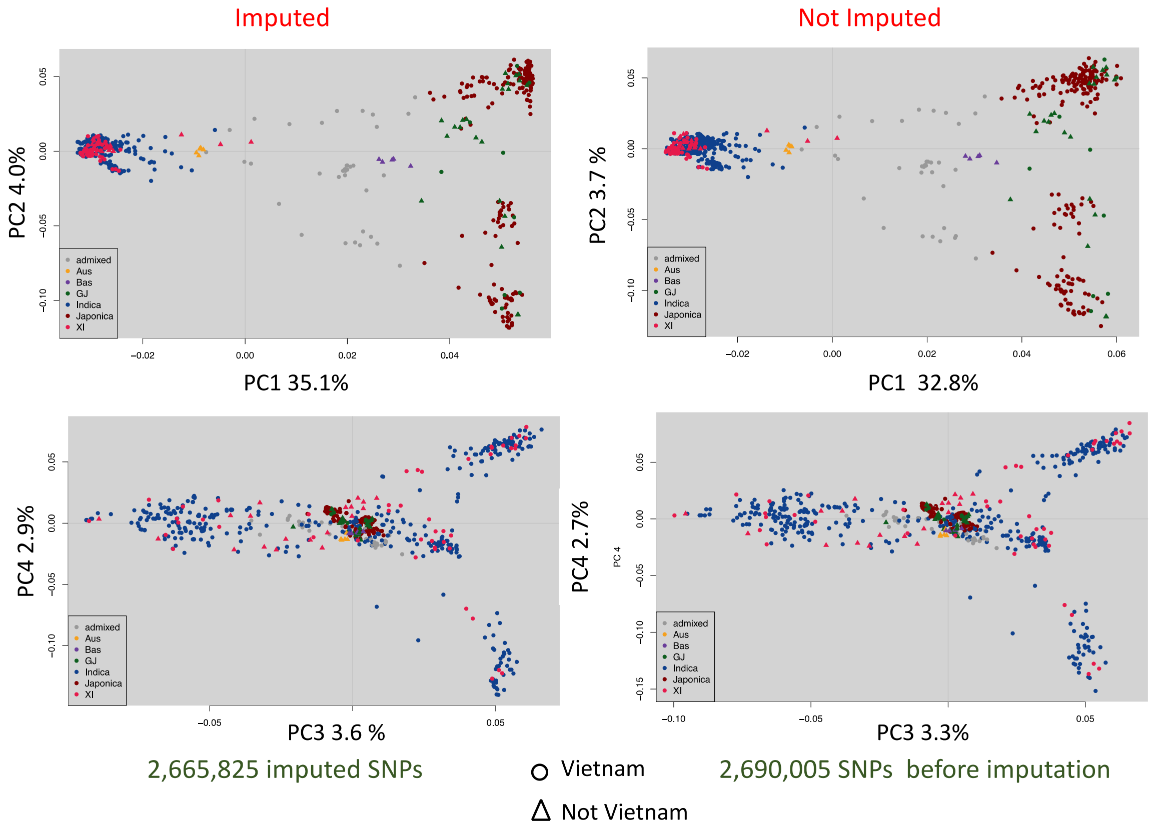


**Figure S16. PCA analysis of 723 samples before and after imputation.**

Comparing the 2,690,005 not imputed SNP set 3 to the 2,665,825 imputed SNP set 4

Both SNP set were filtered for 5% MAF.

Using PC1 and PC2 to separate the Japonica subpopulations.

Using PC3 and PC4 to separate the Indica subpopulations.
