## Supplementary material for "Evidence of selection, adaptation and untapped diversity in Vietnamese rice landraces": SUPPL FIGURE S21

**Figure S21. Allele Plots showing the "*High* impact" SNP position within candidate genes**

| | | | | | | | | | per gene | | | |
|---|---|---|---|---|---|---|---|---|---|---|---|---|
| Region | Gene (MSU) | ^ FST !5 vs I2,I3,I4 (b) | Symbol | impact | SNP | ref | alt | fst for SNP | mean_fst | min_fst | max_fst | number_snps |
| I5_5 | LOC_Os01g65904 | 0.788 | NA | stop_gained | 1_38274711 | A | T | 0.932 | 0.788 | 0.314 | 0.953 | 4 |
| I5_5 | LOC_Os01g65770 | 0.936 | NA | start_lost | 1_38192458 | C | A | 0.939 | 0.936 | 0.932 | 0.939 | 2 |
| I5_17 | LOC_Os03g12840 | 0.477 | DSM3\|OsITPK2 | stop gained | 3_6907102 | T | G | 0.043 | 0.477 | 0.000 | 0.963 | 23 |
| I5_29 | LOC_Os04g58740 | 0.818 | NA | start_lost | 4_34939233 | A | G | 0.818 | 0.818 | 0.818 | 0.818 | 1 |
| I5_29 | LOC_Os04g58870 | 0.813 | NA | splice_acceptor_variant&intron_variant | 4_35019178 | C | A | 0.841 | 0.813 | 0.721 | 0.873 | 8 |
| I5_30 | LOC_Os05g02260 | 0.617 | bip130 | stop_gained | 5_719563 | G | T | 0.626 | 0.617 | 0.151 | 0.737 | 36 |
| I5_37 | LOC_Os08g09110 | 0.904 | NA | stop_gained | 8_5278838 | T | A | NA | 0.904 | 0.239 | 0.978 | 12 |
| I5_47 | LOC_Os10g35604 | 0.661 | NA | stop_gained | 10_19045026 | G | T | 0.776 | 0.661 | 0.021 | 0.776 | 15 |
| I5_49 | LOC_Os11g09360 | 0.919 | OsFBX398 | splice_donor_variant&intron_variant | 11_5026584 | G | A | 0.924 | 0.919 | 0.723 | 0.953 | 15 |
| I5_49 | LOC_Os11g10070 | 0.721 | OsSEU2 | splice_donor_variant&intron_variant | 11_5424384 | G | A | 0.900 | 0.721 | 0.063 | 0.912 | 42 |
| | LOC_Os06g03860 | nan | OsSPX-MFS3 | splice_acceptor_variant&intron_variant | 6_1559993 | A | C | nan | | | | |

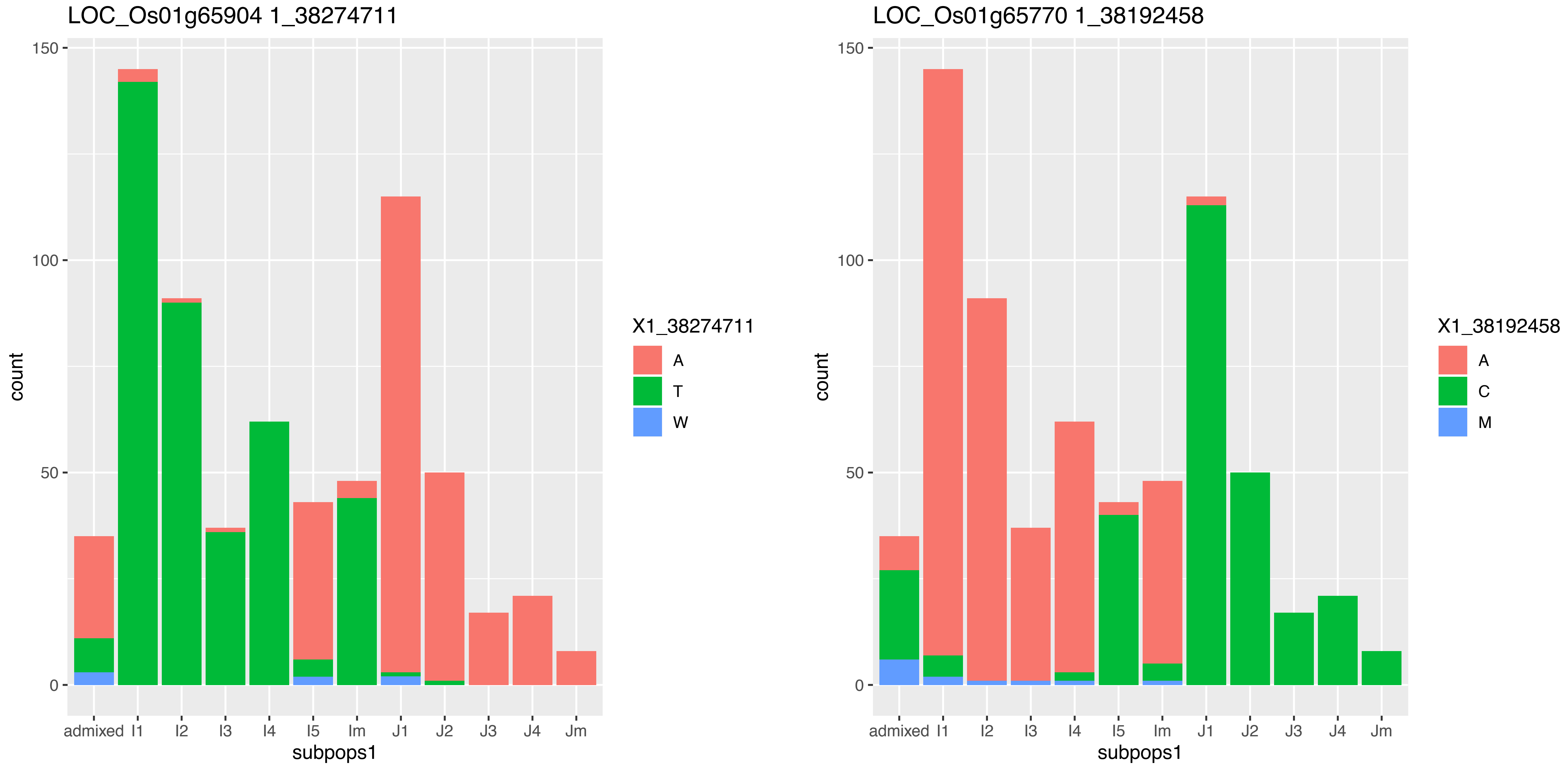
LOC_Os01g65904 1_38274711
count
subpops1
admixed
I1
I2
I3
I4
I5
Im
J1
J2
J3
J4
Jm
0
50
100
150
X1_38274711
A
T
W
LOC_Os01g65770 1_38192458
count
subpops1
X1_38192458
A
C
M

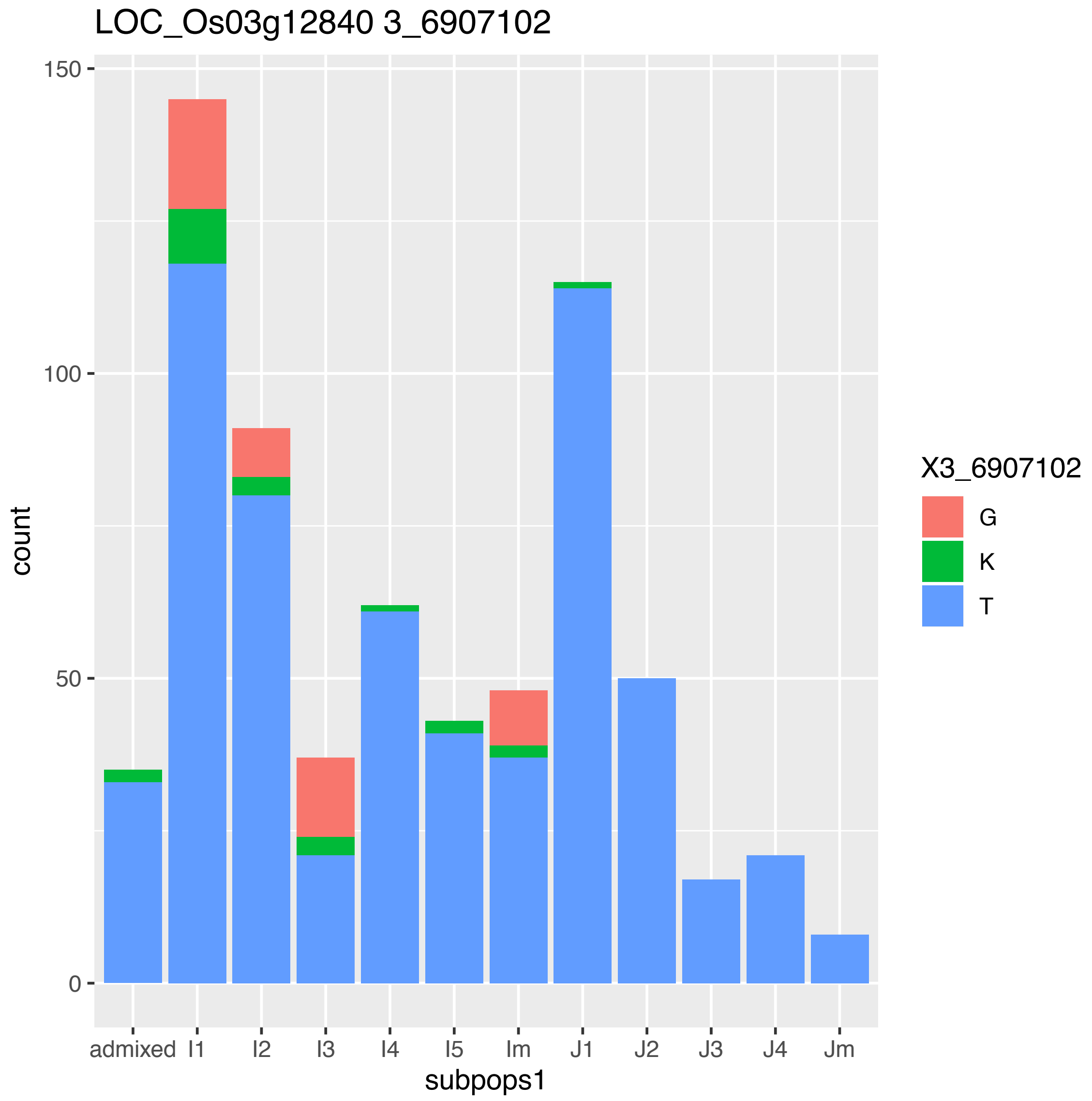
LOC_Os03g12840 3_6907102
count
subpops1
admixed
I1
I2
I3
I4
I5
Im
J1
J2
J3
J4
Jm
0
50
100
150
X3_6907102
G
K
T
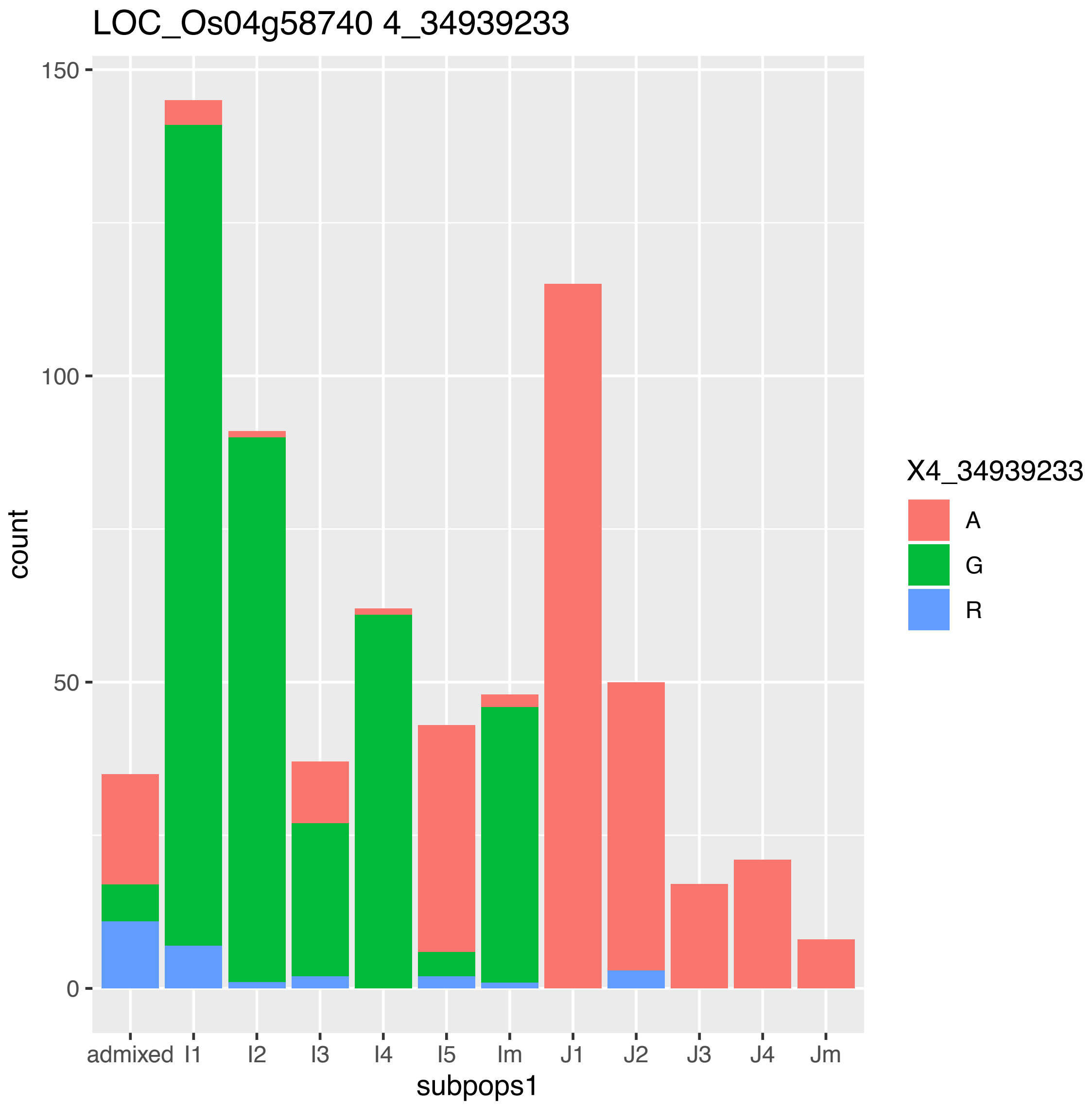
LOC_Os04g58740 4_34939233
count
subpops1
admixed
I1
I2
I3
I4
I5
Im
J1
J2
J3
J4
Jm
0
50
100
150
X4_34939233
A
G
R

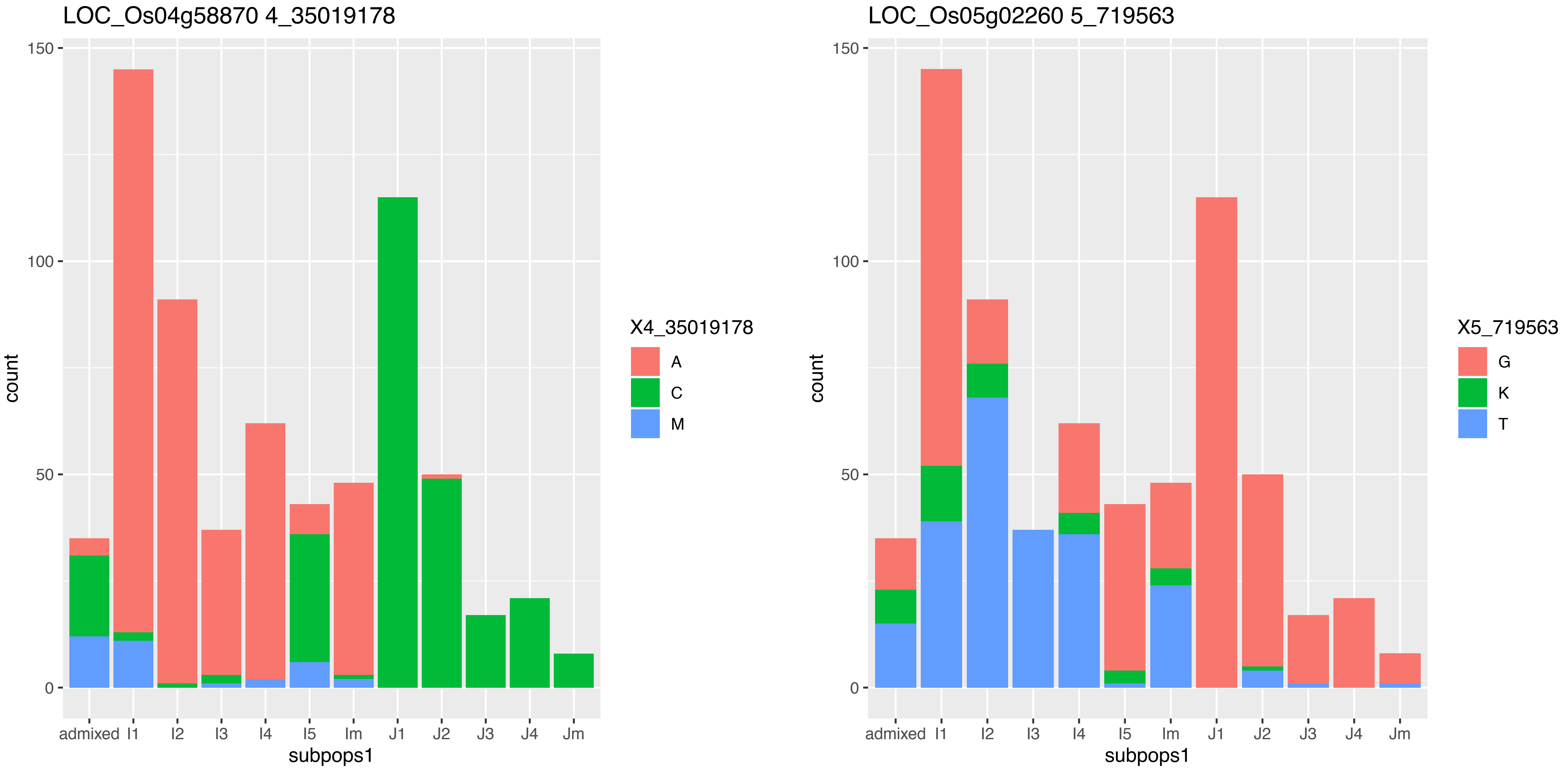
LOC_Os04g58870 4_35019178
count
150
100
50
0
admixed
I1
I2
I3
I4
I5
Im
J1
J2
J3
J4
Jm
subpops1
X4_35019178
A
C
M
LOC_Os05g02260 5_719563
X5_719563
G
K
T

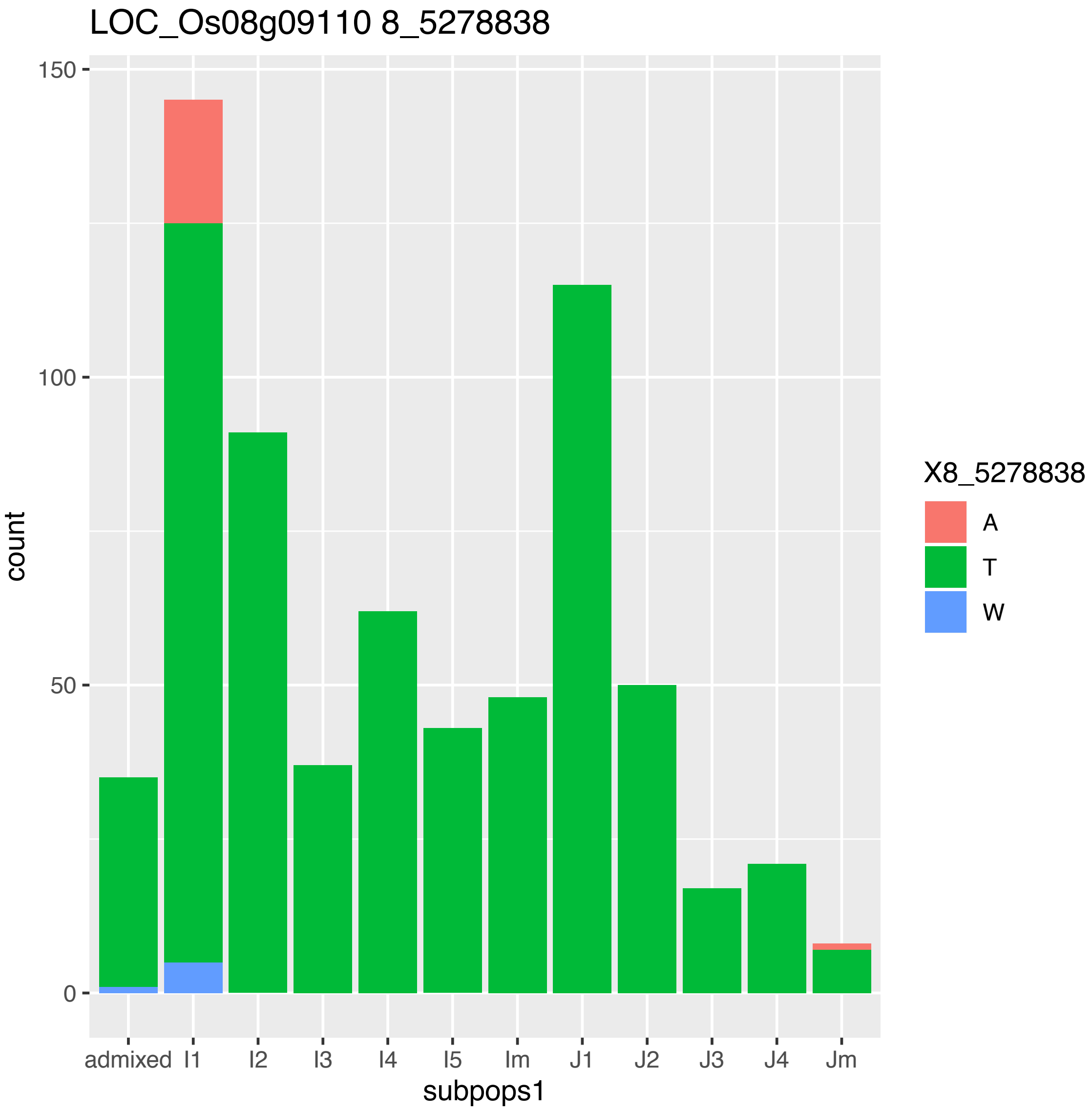
LOC_Os08g09110 8_5278838
count
150
100
50
0
admixed
I1
I2
I3
I4
I5
Im
J1
J2
J3
J4
Jm
subpops1
X8_5278838
A
T
W

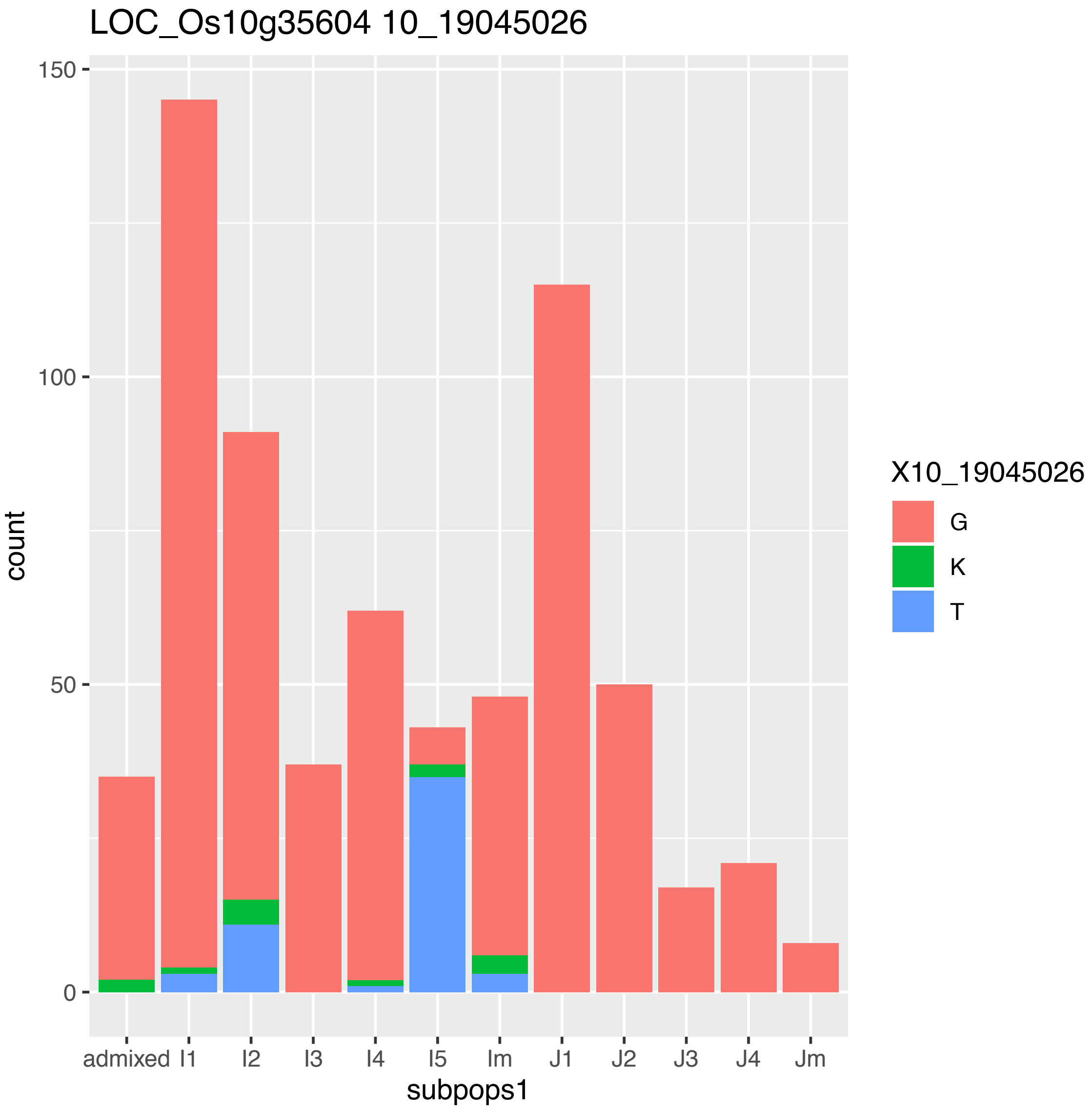
LOC_Os10g35604 10_19045026
count
150
100
50
0
admixed
I1
I2
I3
I4
I5
Im
J1
J2
J3
J4
Jm
subpops1
X10_19045026
G
K
T

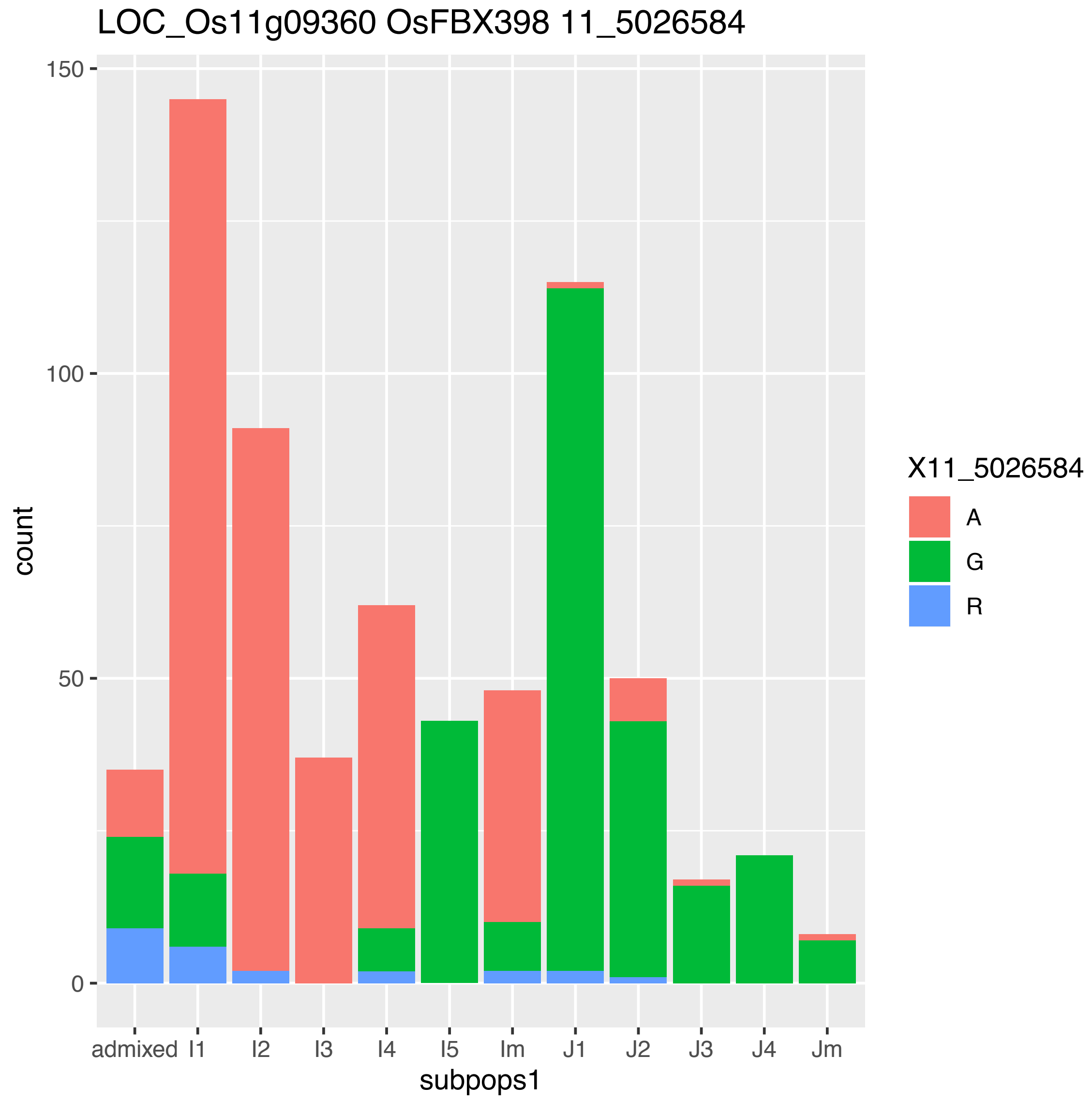
LOC_Os11g09360 OsFBX398 11_5026584
count
150
100
50
0
admixed
I1
I2
I3
I4
I5
Im
J1
J2
J3
J4
Jm
subpops1
X11_5026584
A
G
R

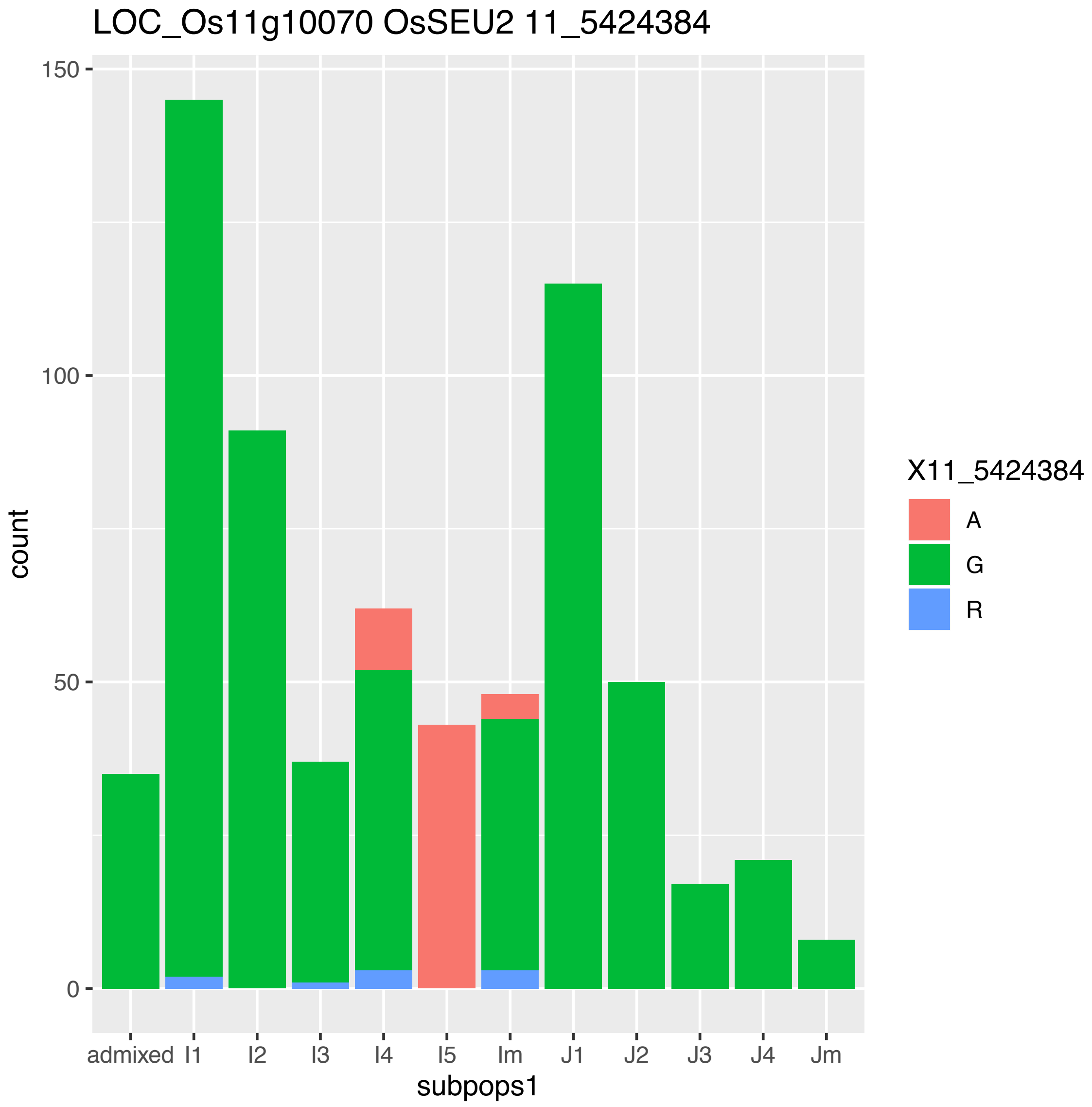
LOC_Os11g10070 OsSEU2 11_5424384
count
150
100
50
0
admixed
I1
I2
I3
I4
I5
Im
J1
J2
J3
J4
Jm
subpops1
X11_5424384
A
G
R

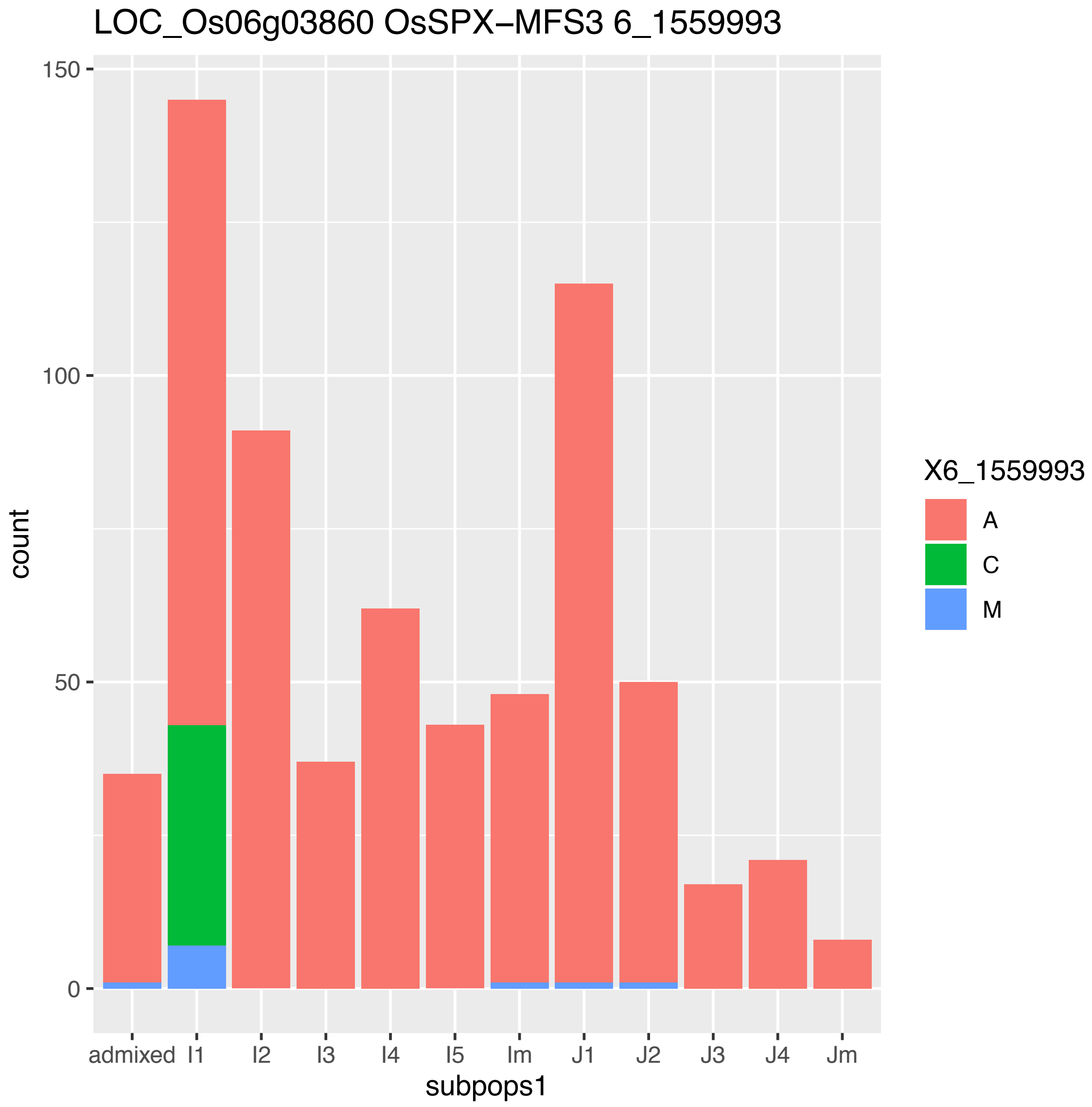
LOC_Os06g03860 OsSPX−MFS3 6_1559993
150
100
50
0
count
admixed
I1
I2
I3
I4
I5
Im
J1
J2
J3
J4
Jm
subpops1
X6_1559993
A
C
M
