## Supplementary material for "Evidence of selection, adaptation and untapped diversity in Vietnamese rice landraces": SUPPL FIGURES S19 AND S20

**Figure S19. Chromosome plots of regions selected in each Indica subpopulation showing the regions selected against each individual subpopulation and the shaded final selected regions which were selected against three subpopulations.**

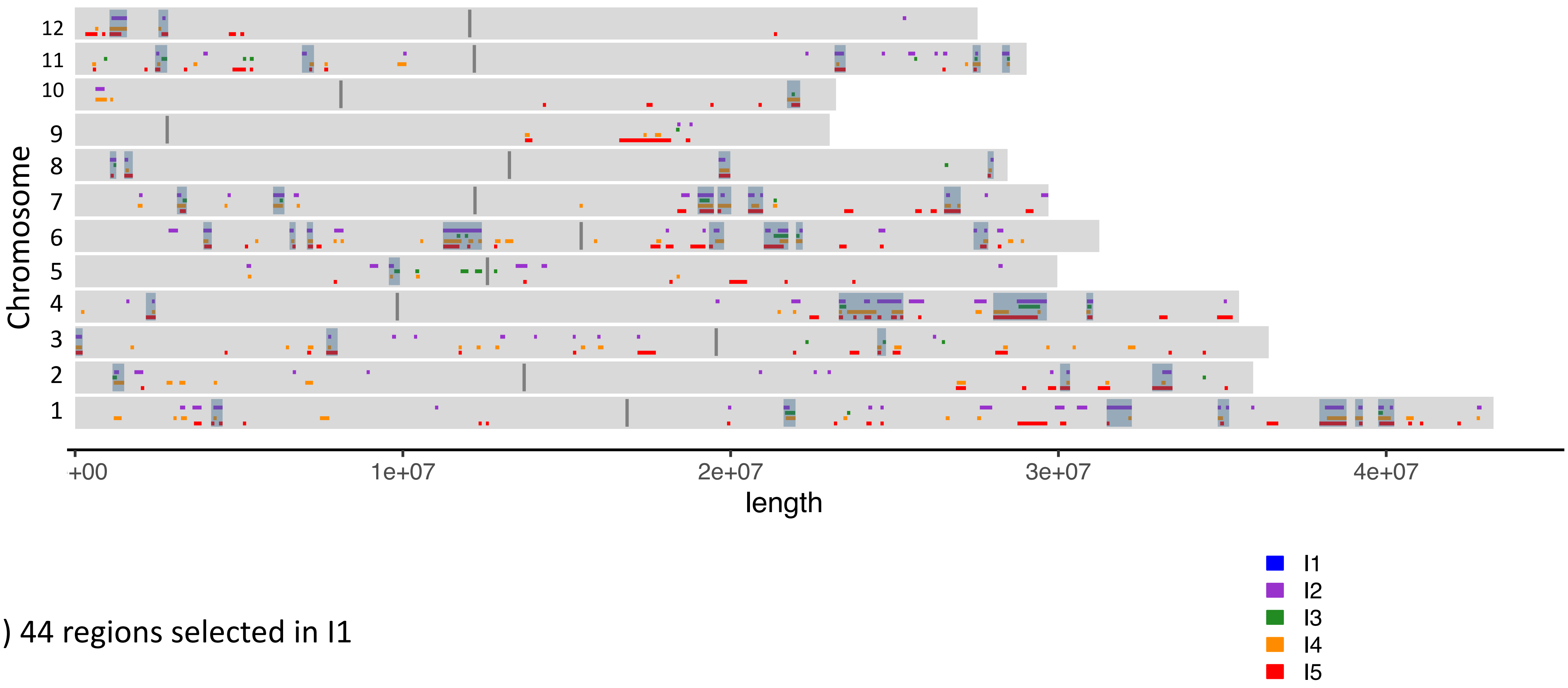


a) 44 regions selected in I1

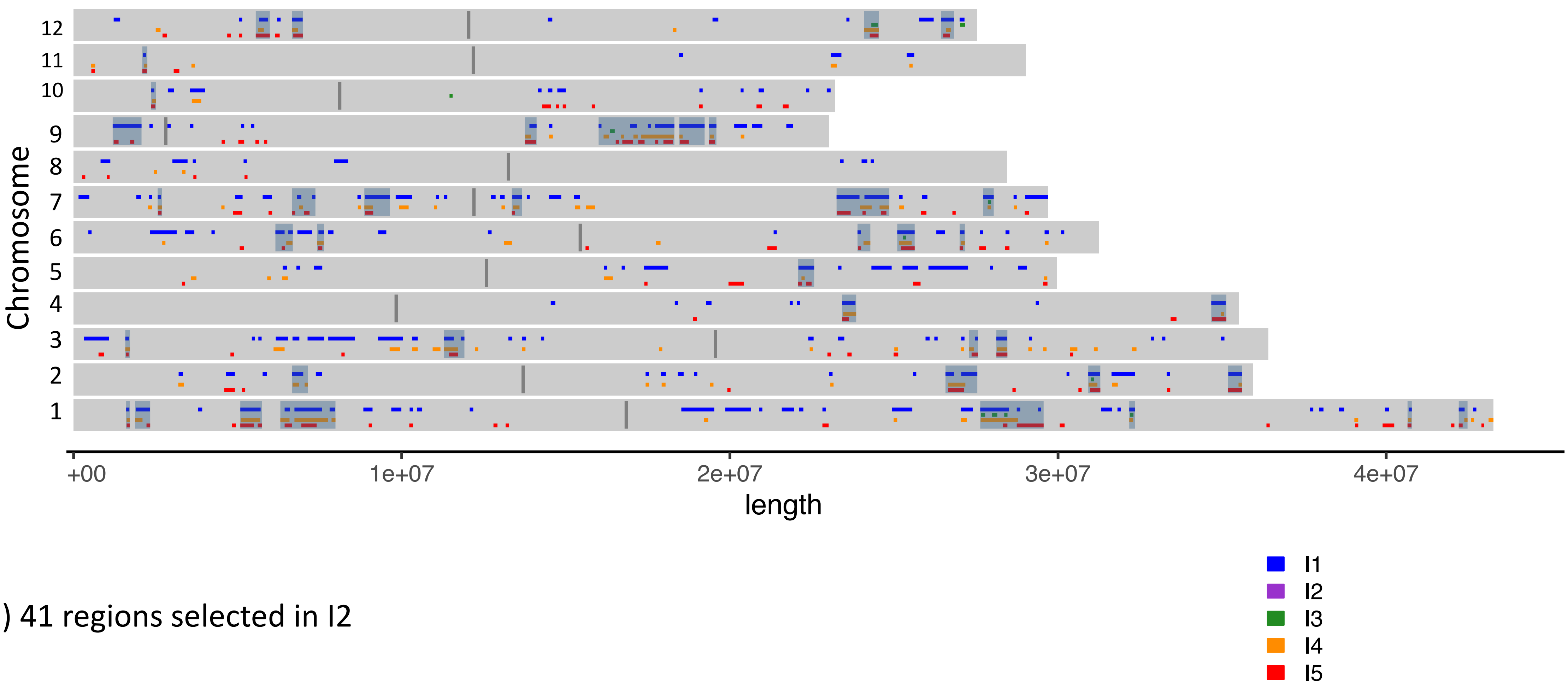


b) 41 regions selected in I2

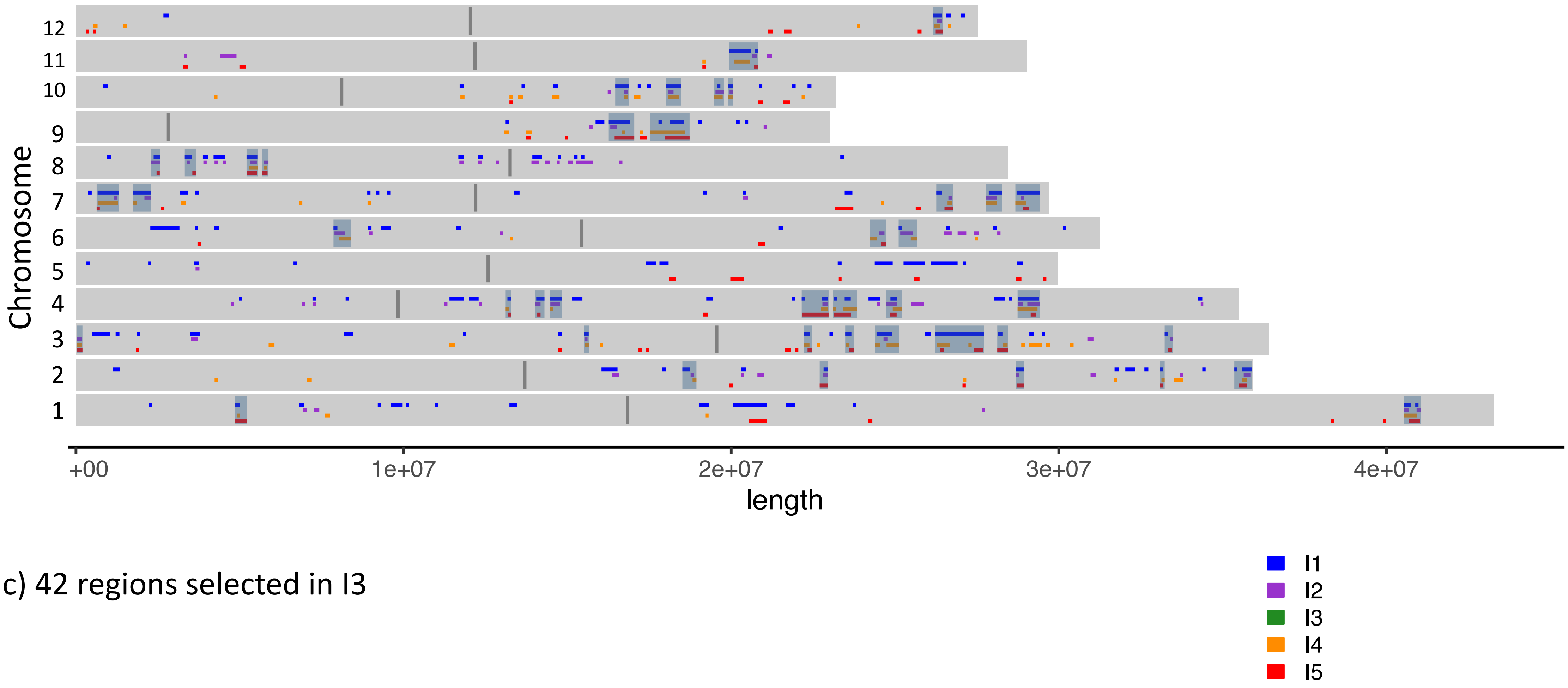


c) 42 regions selected in I3

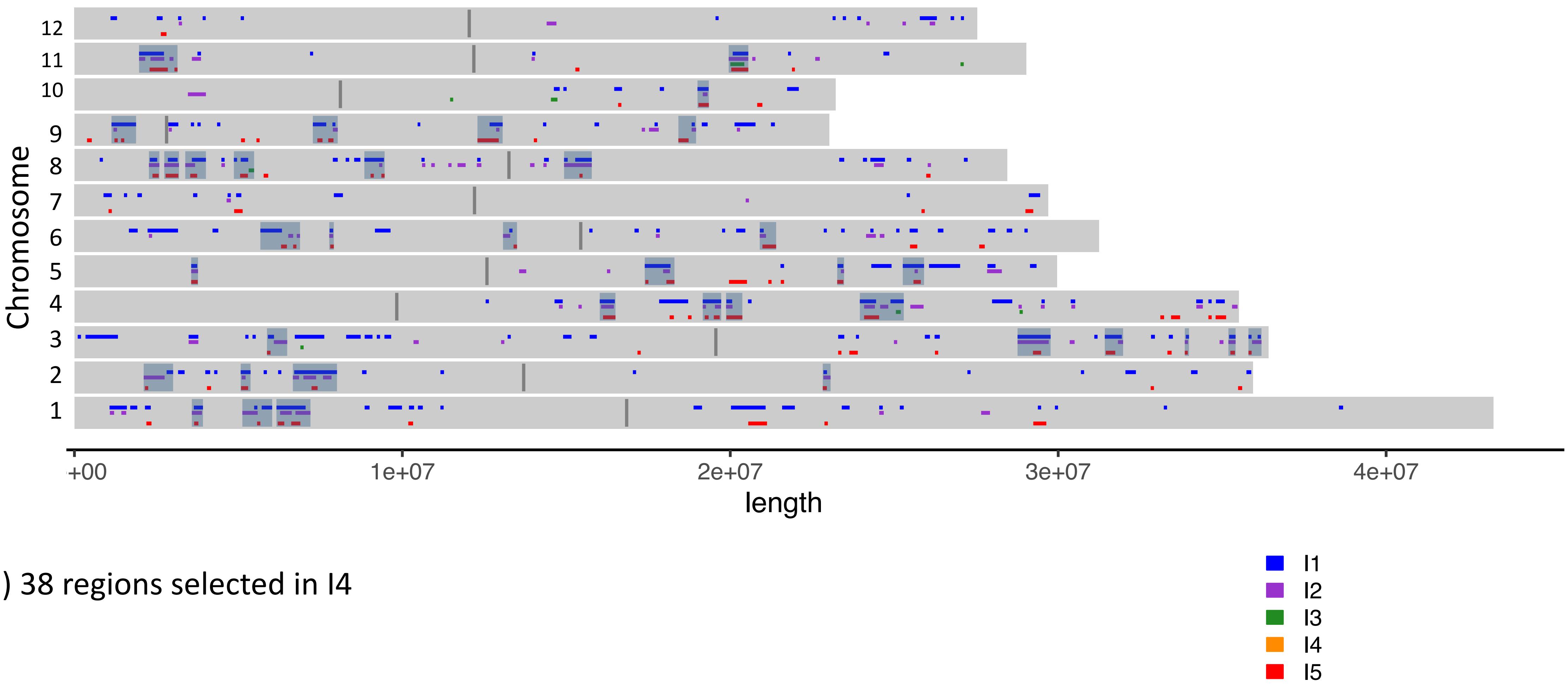


d) 38 regions selected in I4

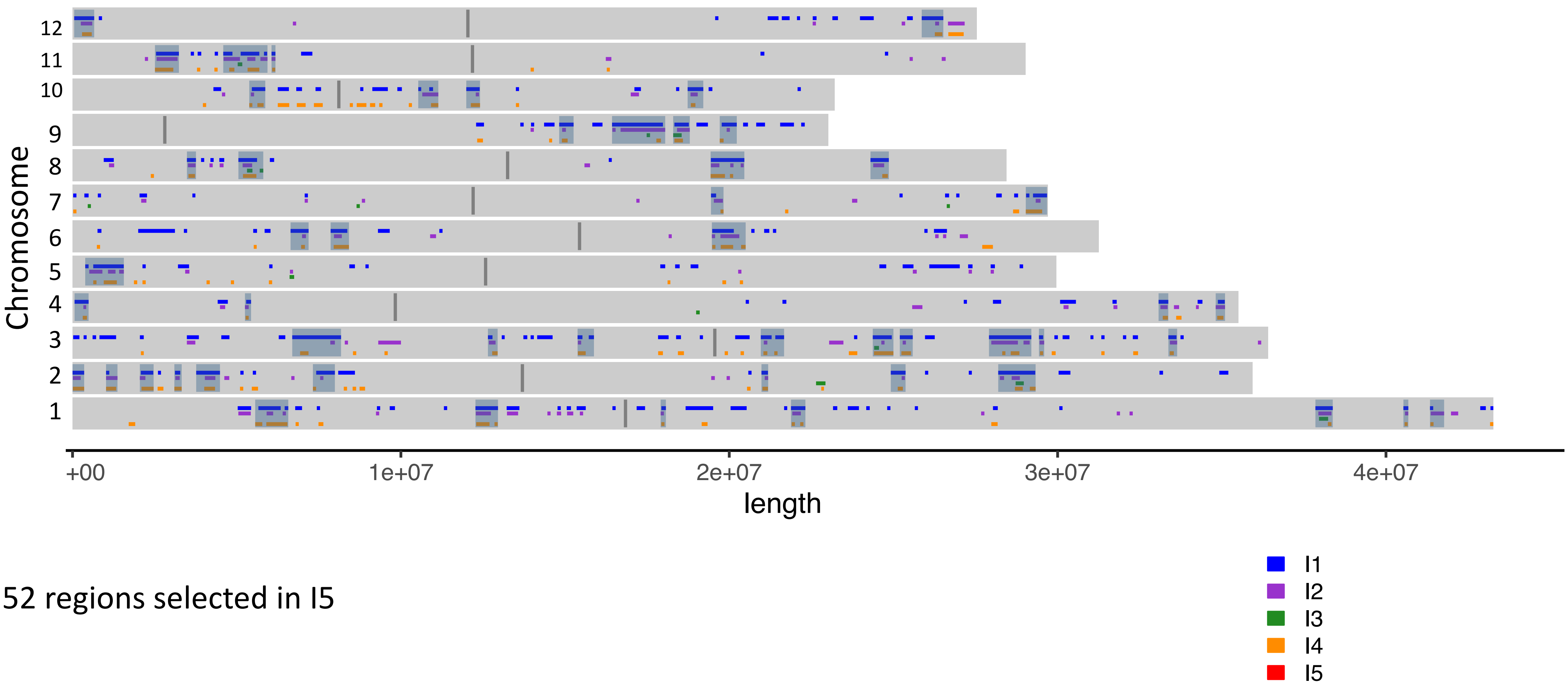


e) 52 regions selected in I5

**Figure S20. Chromosome plots of regions selected in each Japonica subpopulation showing the regions selected against each individual subpopulation and the shaded final selected regions which were selected against two subpopulations.**

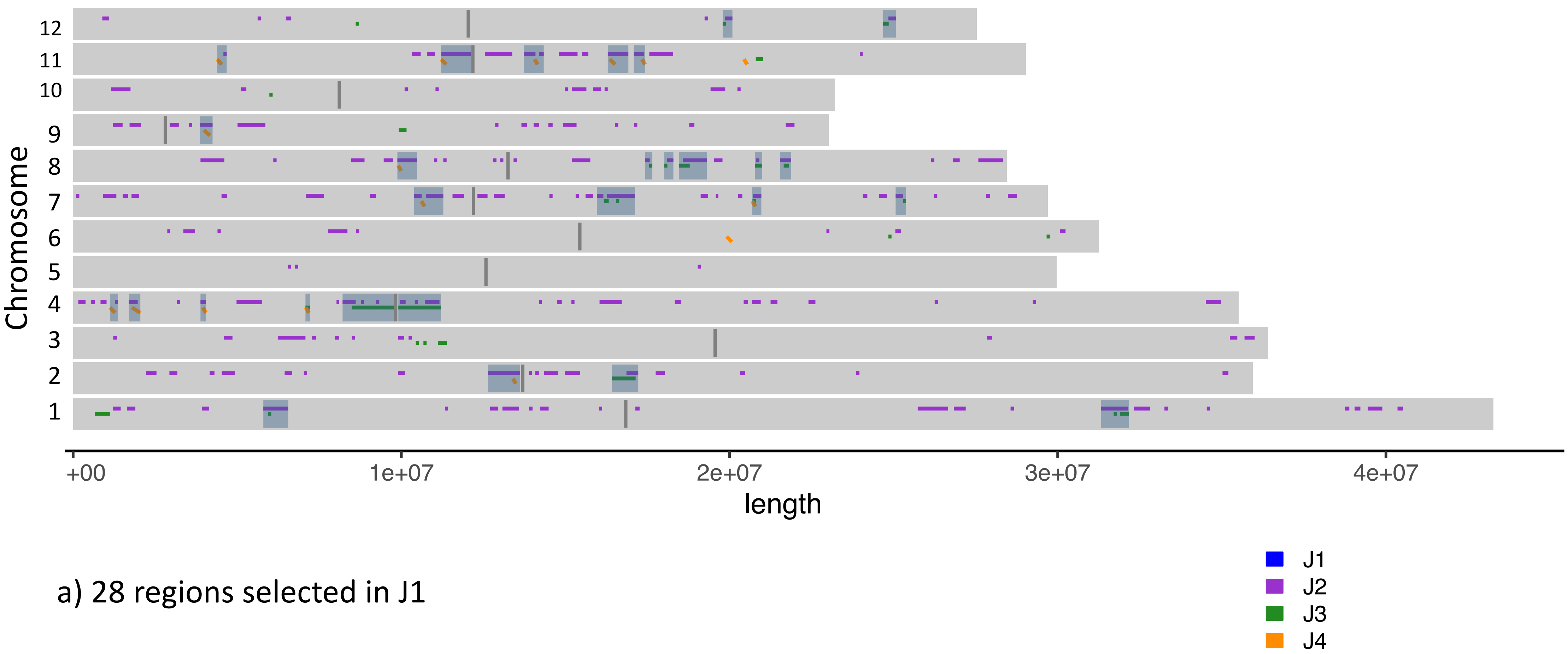


a) 28 regions selected in J1

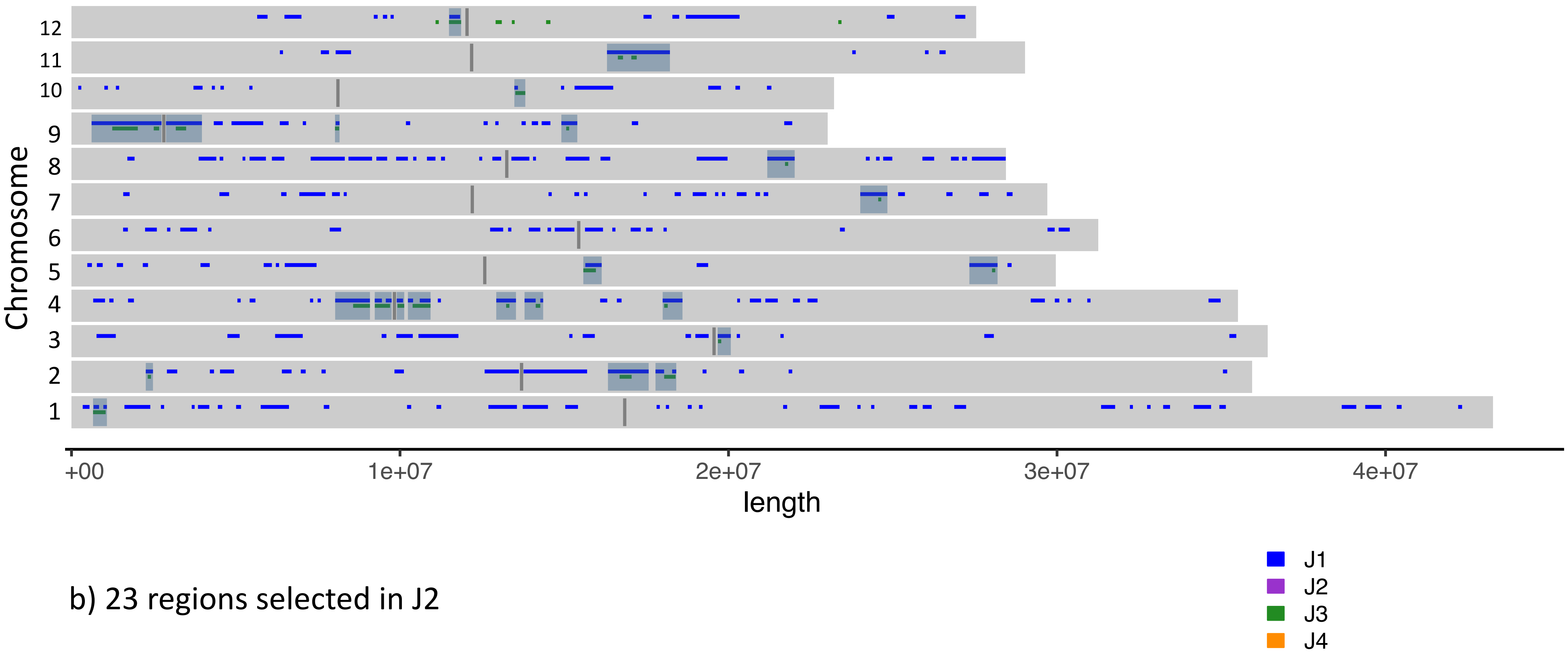


b) 23 regions selected in J2

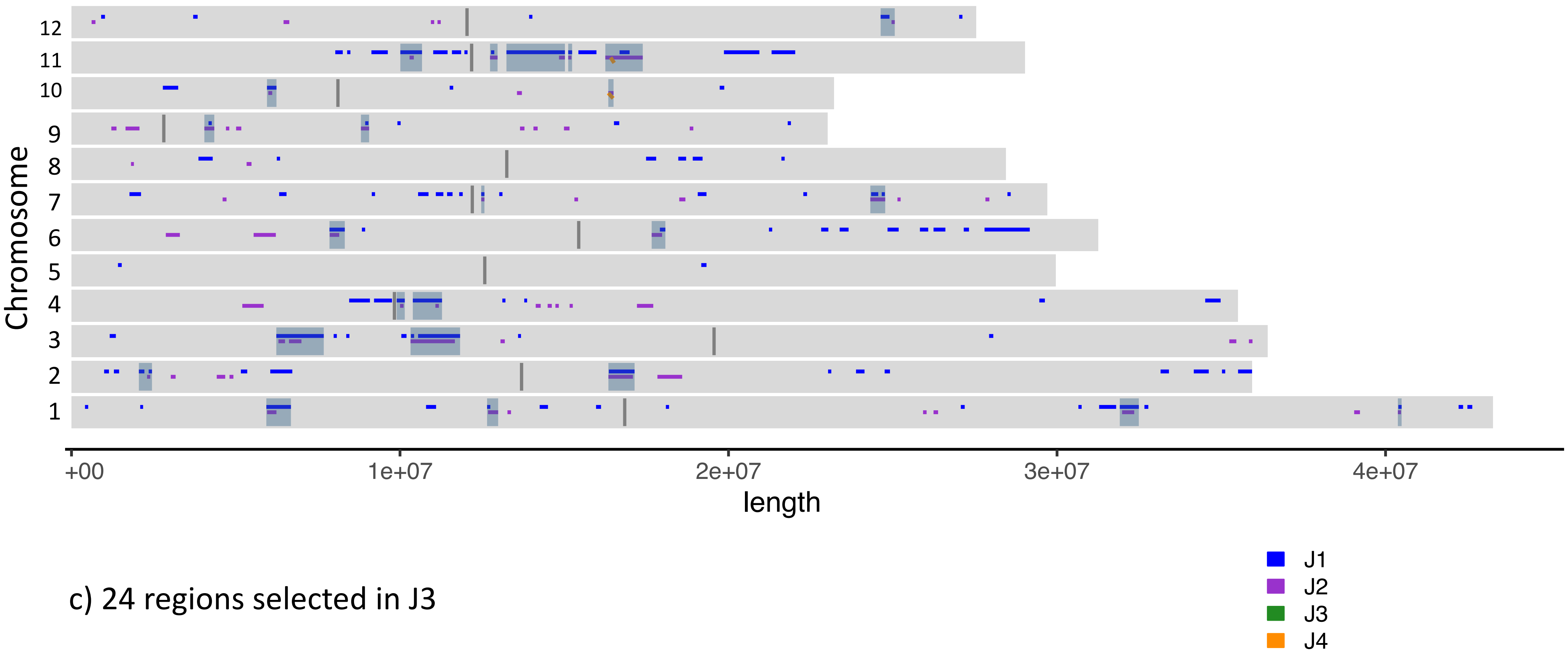


c) 24 regions selected in J3

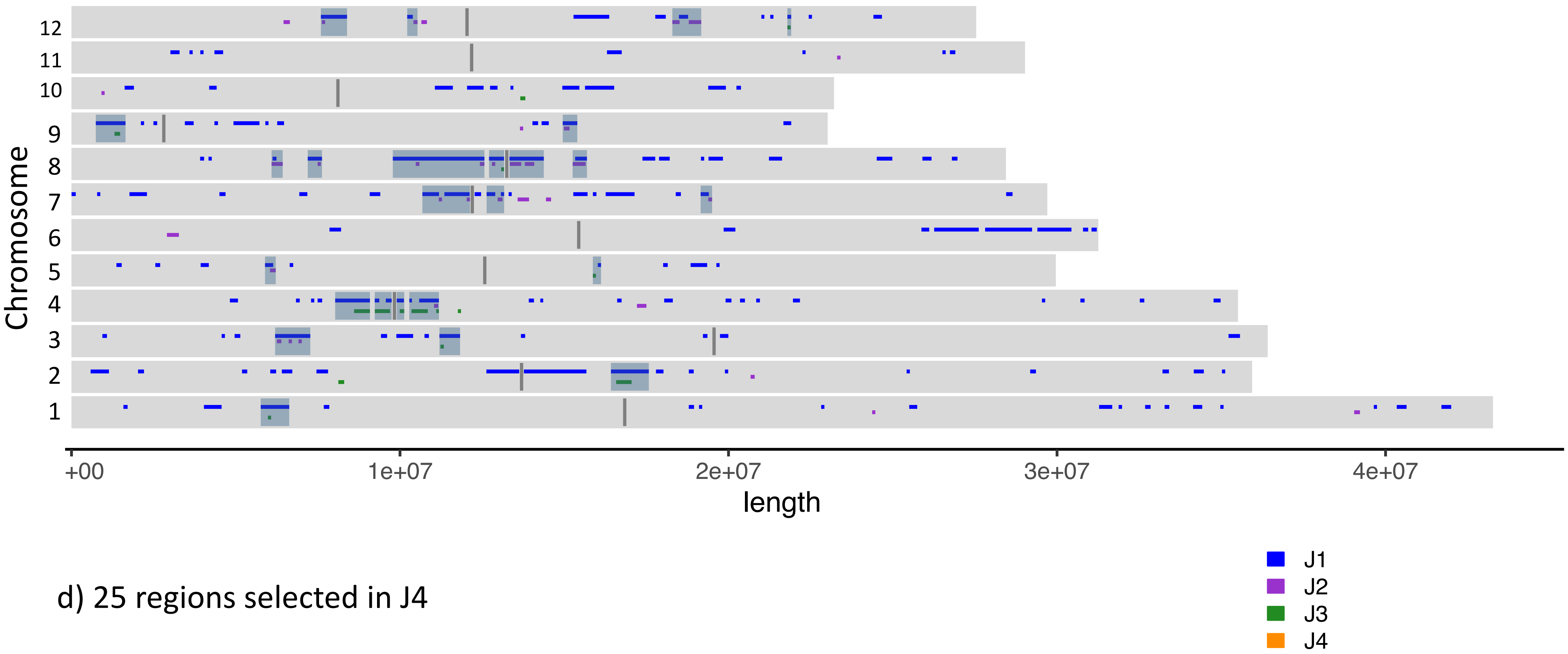


d) 25 regions selected in J4
