## Supplementary material for "Evidence of selection, adaptation and untapped diversity in Vietnamese rice landraces": SUPPL FIGURE S18

**Figure S18.** GWAS Manhattan and qq plots for the full panel and Indica and Japonica subpanels for Leaf Pubescence, Culm Number, Diameter Internode, Culm Length, Panicle Length and Floret Pubescence

Full Panel
672 samples
328 phenotypes

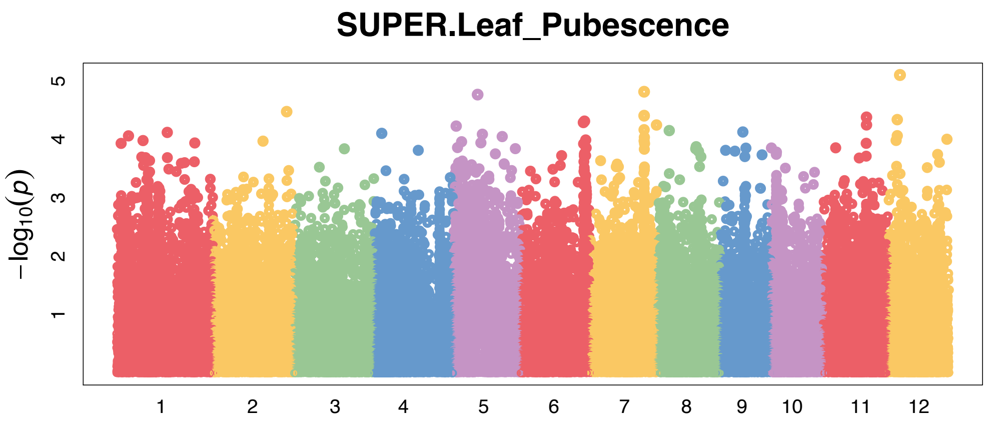


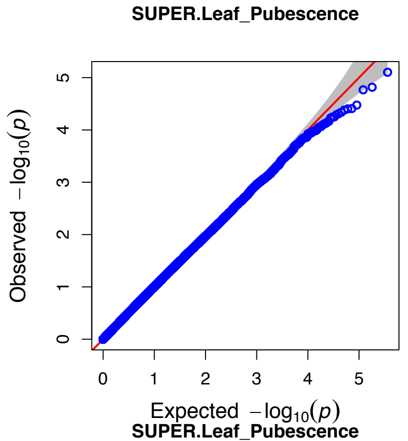


Indica Panel
426 samples
170 phenotypes

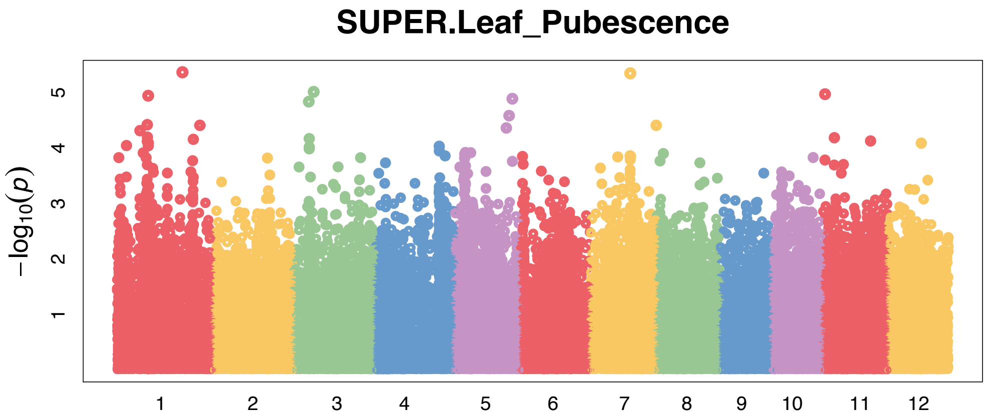


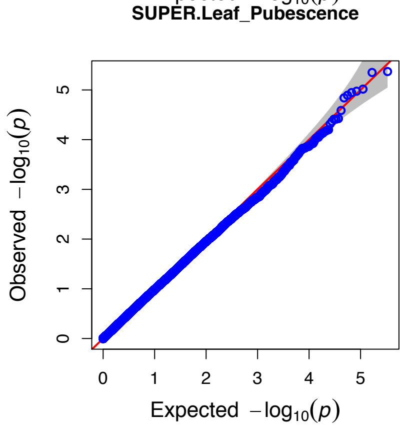


Japonica Panel
211 samples
134 phenotypes

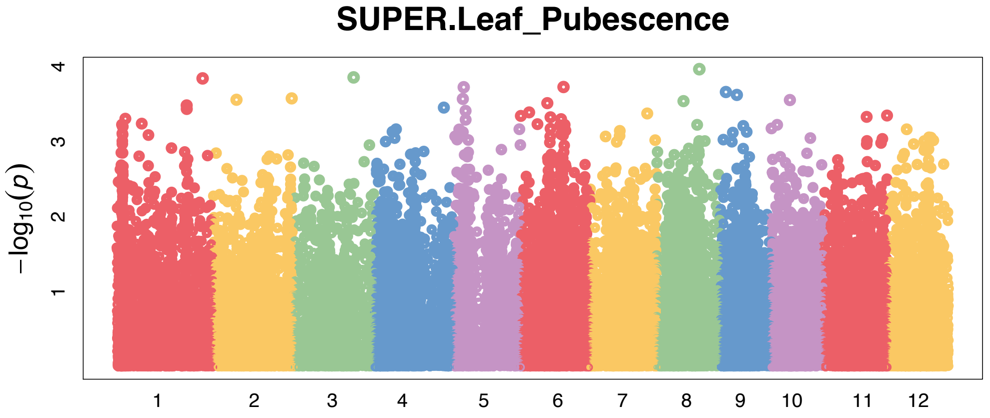


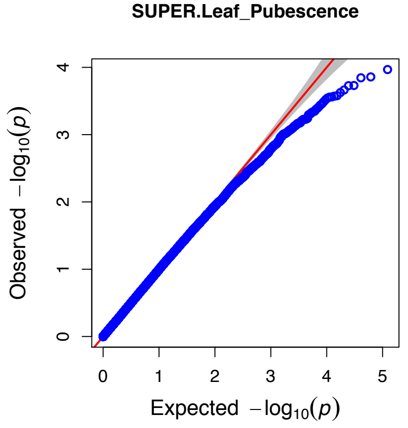

Full Panel
672 samples
454 phenotypes

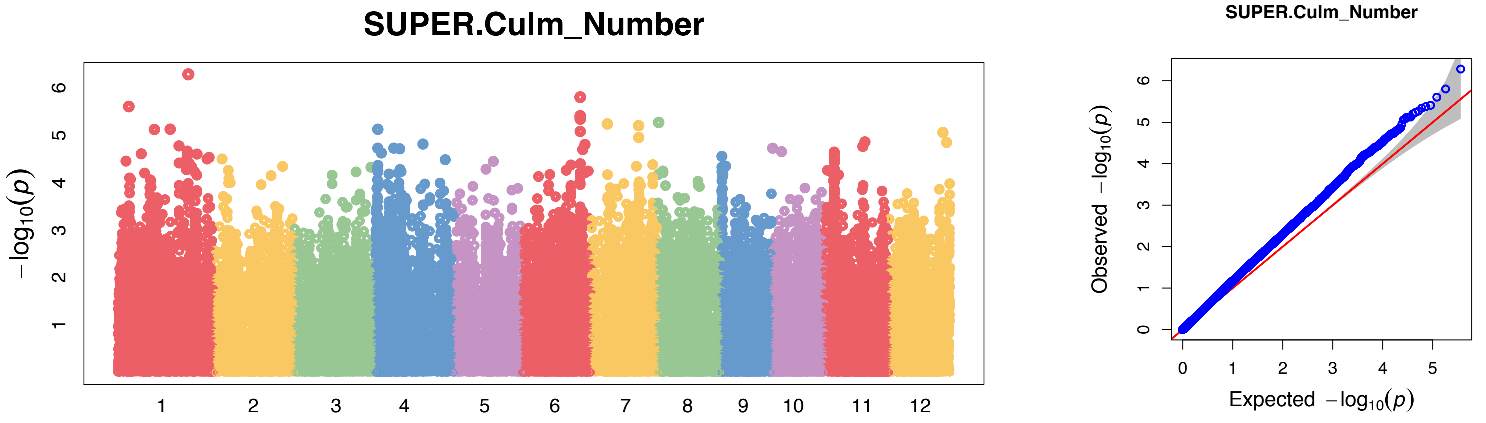


Indica Panel
426 samples
254 phenotypes

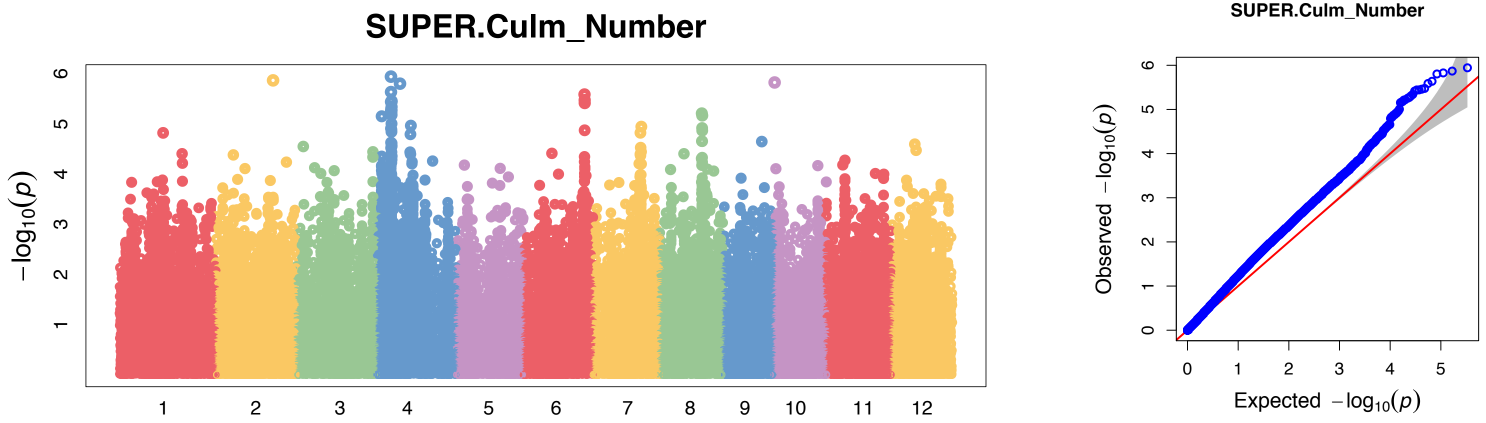


Japonica Panel
211 samples
172 phenotypes

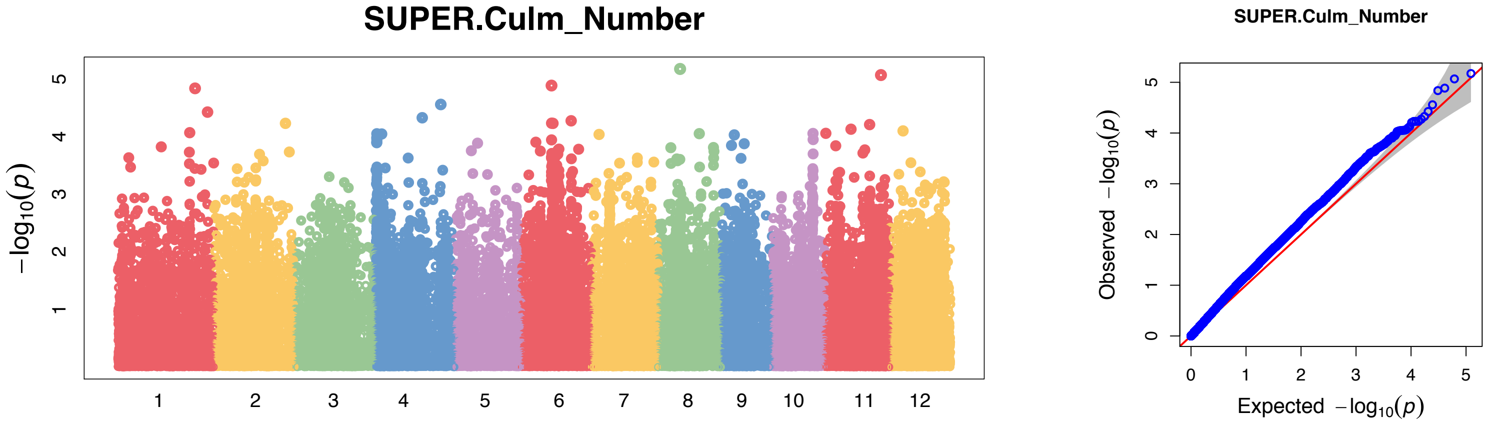

Full Panel
672 samples
485 phenotypes

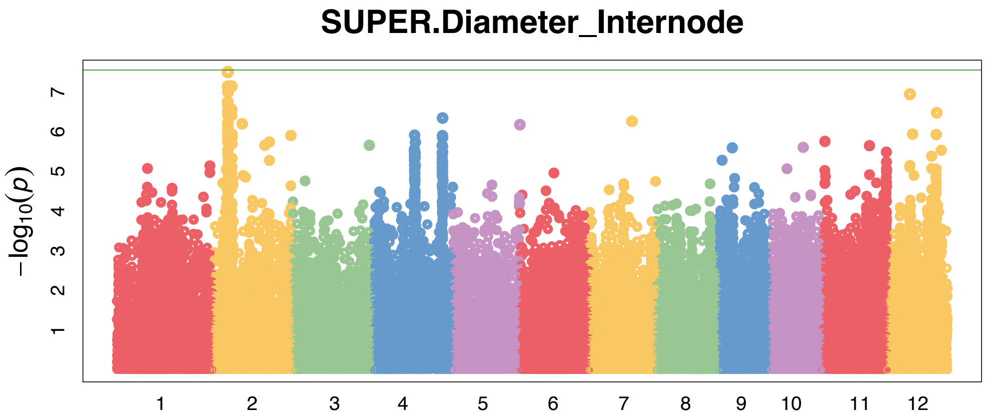


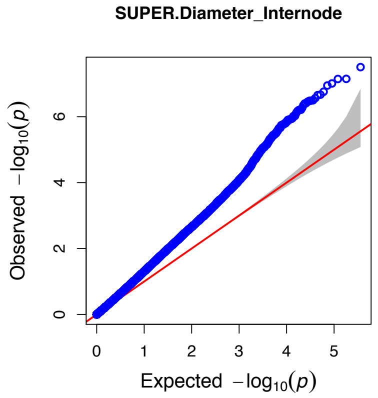


Indica Panel
426 samples
284 phenotypes

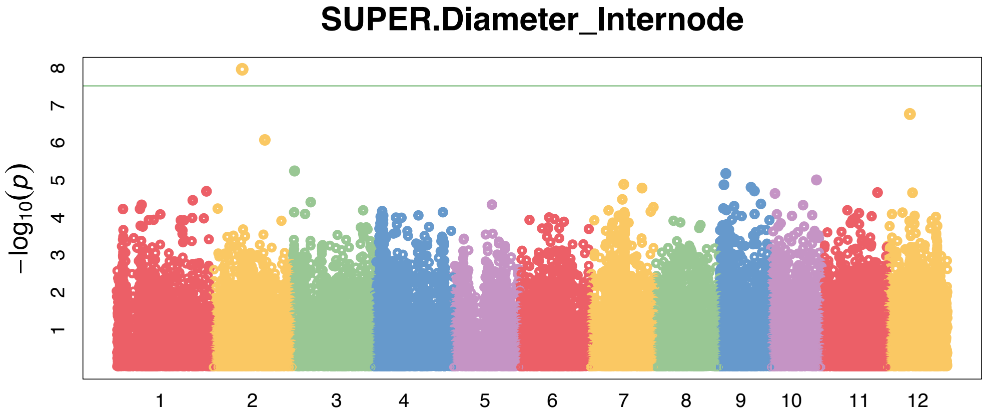


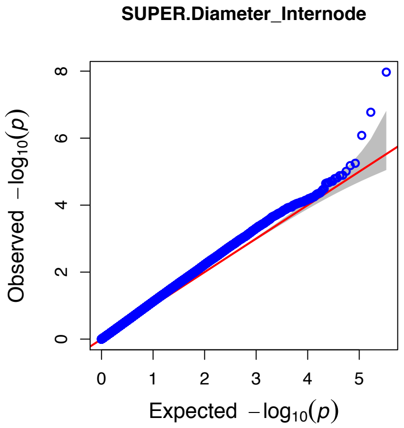


Japonica Panel
211 samples
174 phenotypes

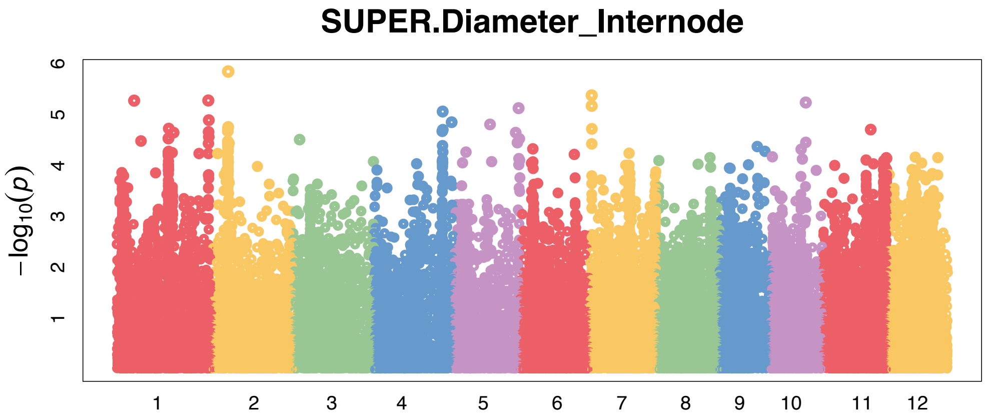


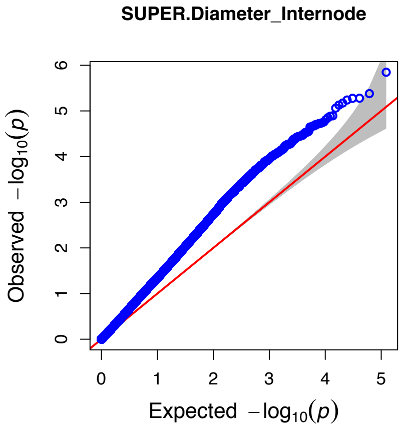

Full Panel
672 samples
485 phenotypes

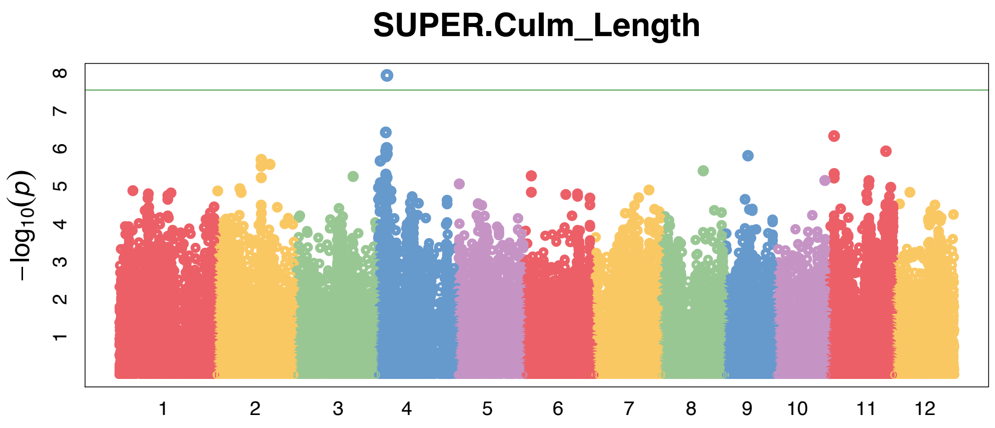


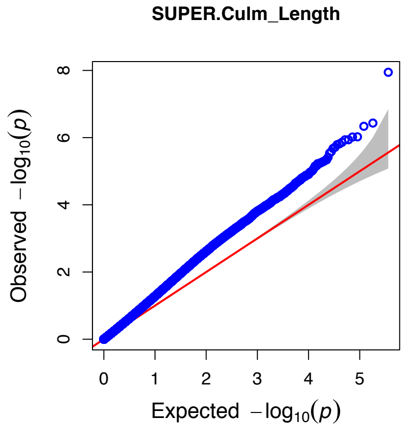


Indica Panel
426 samples
282 phenotypes

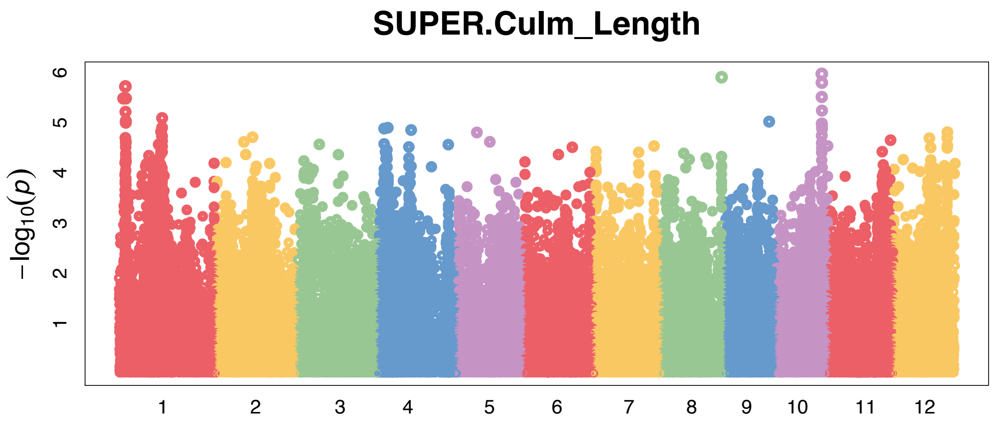


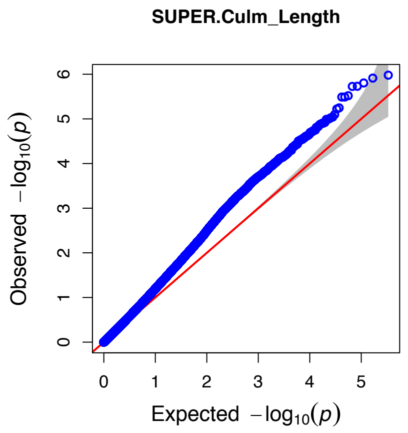


Japonica Panel
211 samples
175 phenotypes

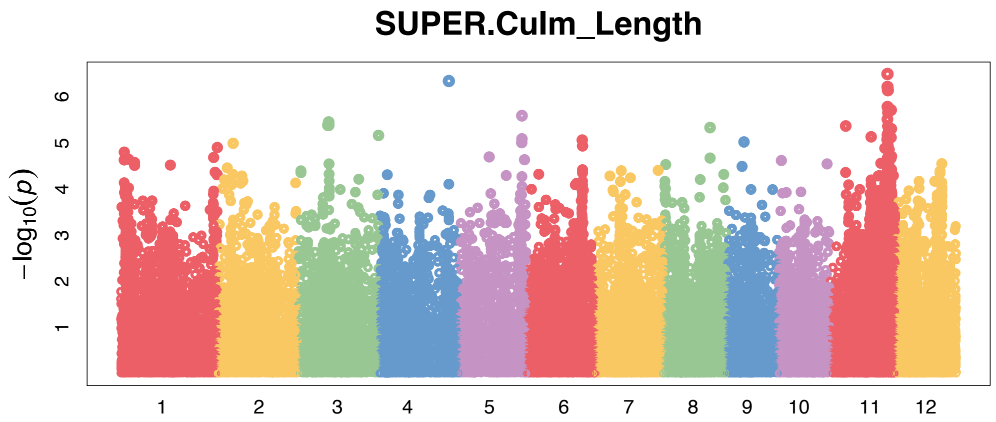


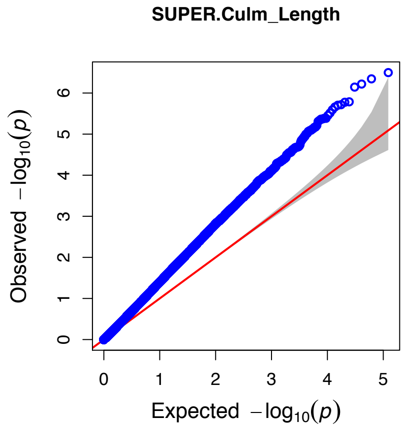

Full Panel
672 samples
486 phenotypes

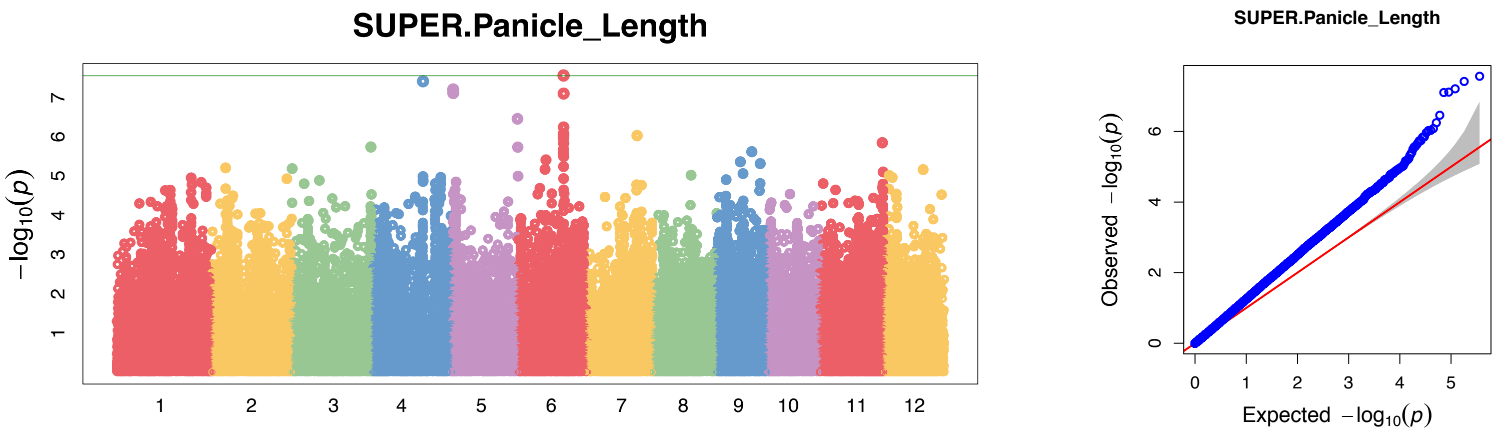


Indica Panel
426 samples
283 phenotypes

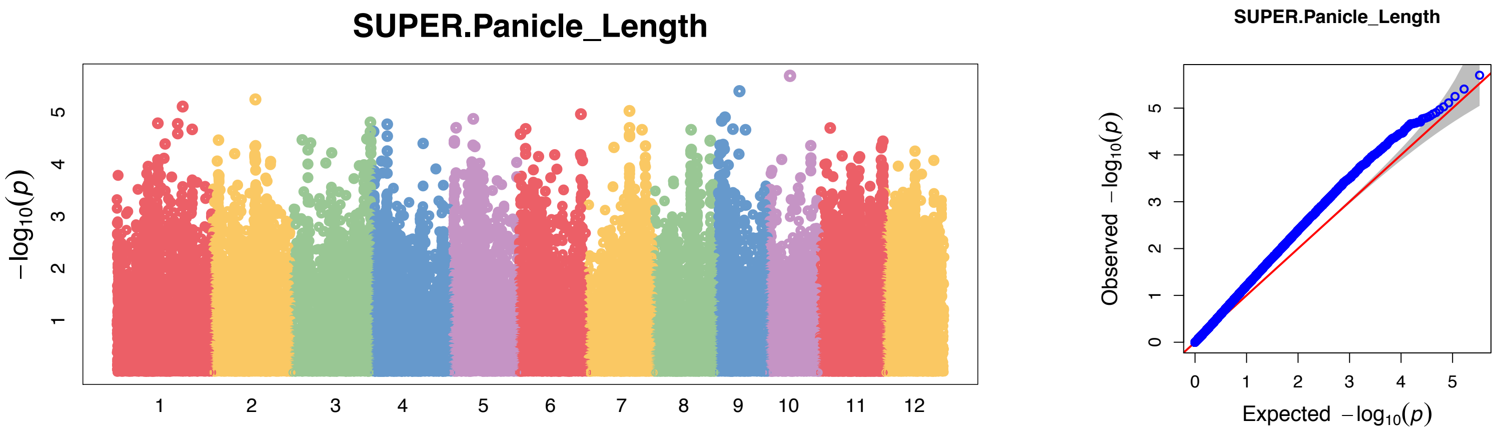


Japonica Panel
211 samples
175 phenotypes

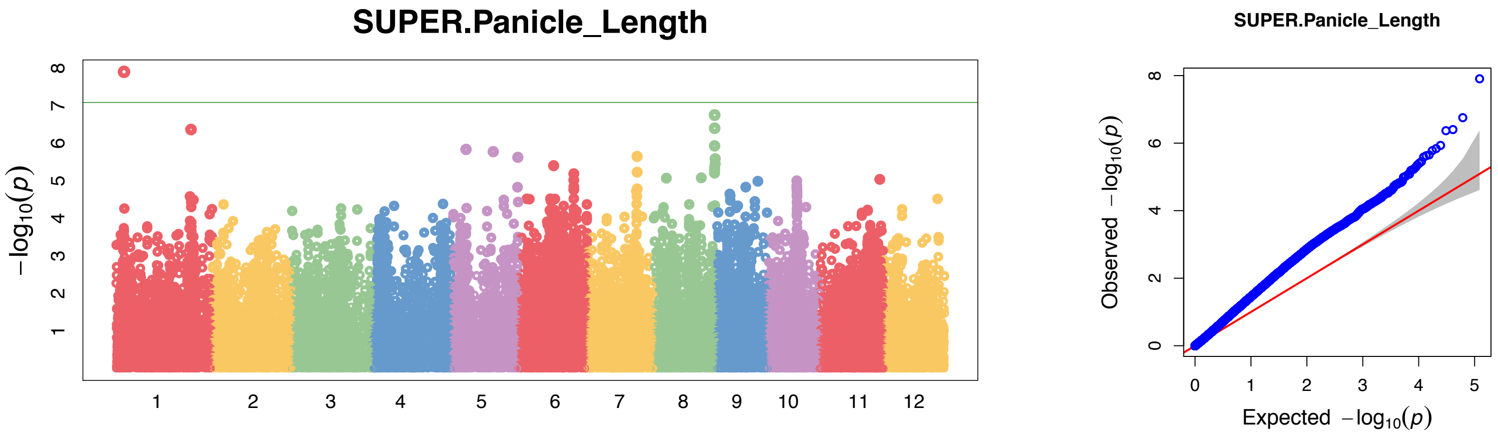

Full Panel
672 samples
488 phenotypes

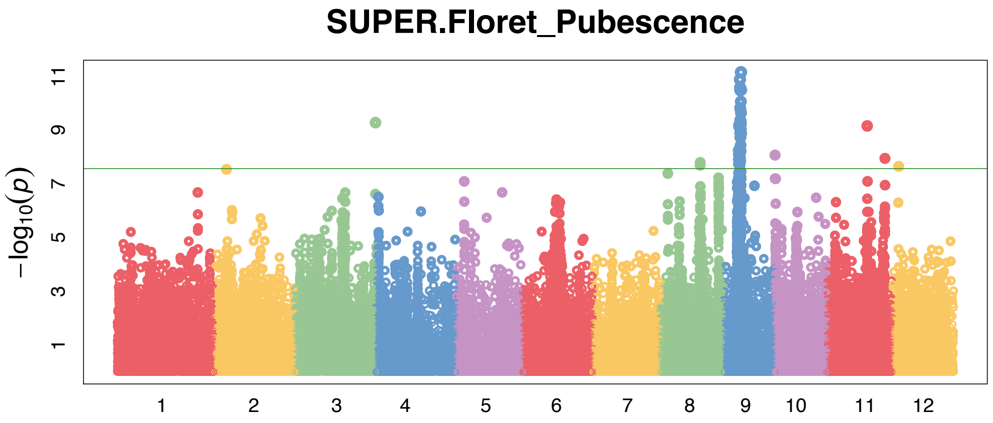


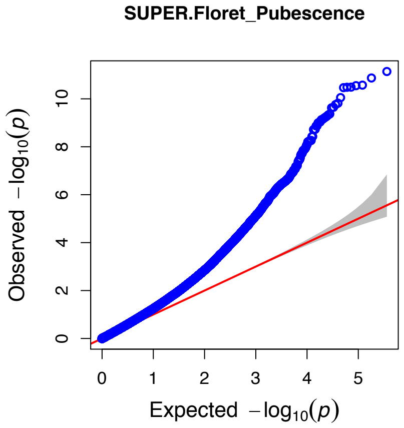


Indica Panel
426 samples
284 phenotypes

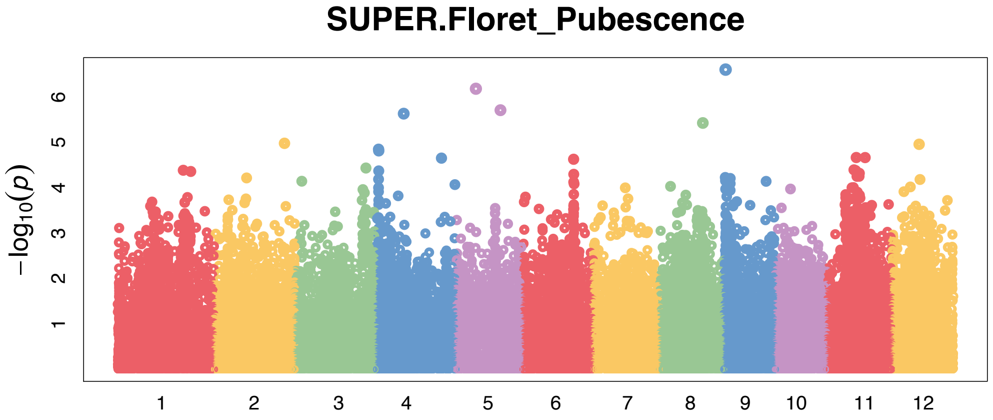


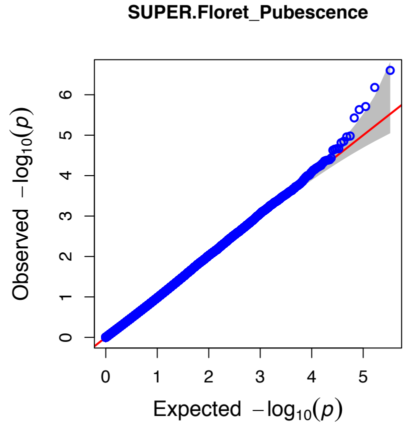


Japonica Panel
211 samples
176 phenotypes

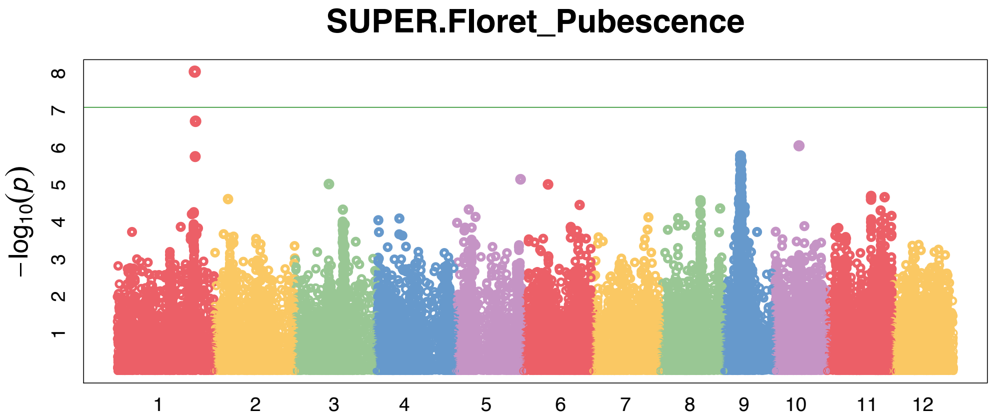


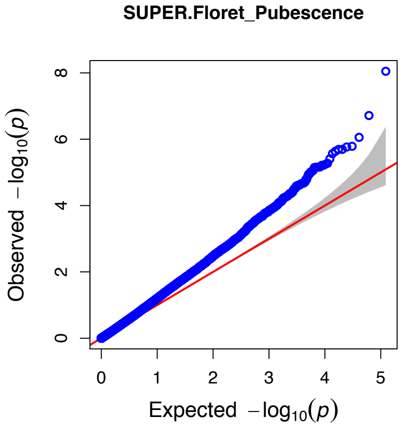
