## Supplementary material for "Evidence of selection, adaptation and untapped diversity in Vietnamese rice landraces": SUPPL FIGURE S17

**Figure S17.** GWAS Manhattan and qq plots for the full panel and Indica and Japonica subpanels for Grain Length, Grain Width, Grain length-to-width ratio, Heading Date, Culm Strength, Leaf Length and Leaf Width.

Full Panel
672 samples
501 phenotypes

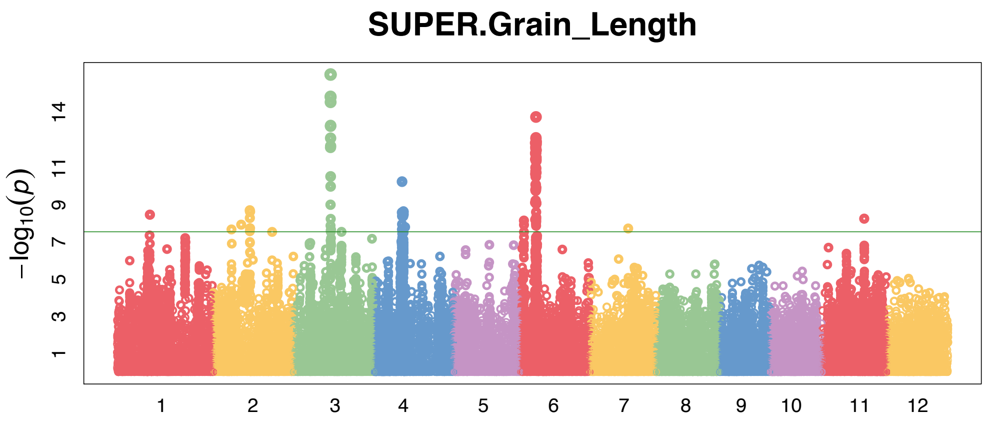


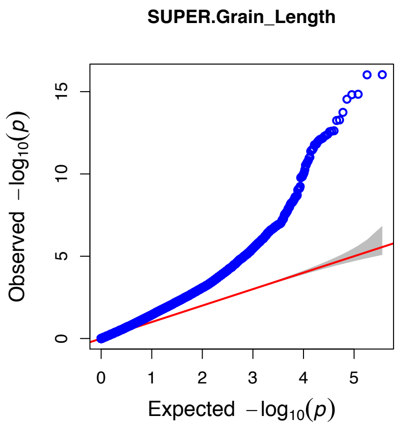


Indica Panel
426 samples
295 phenotypes

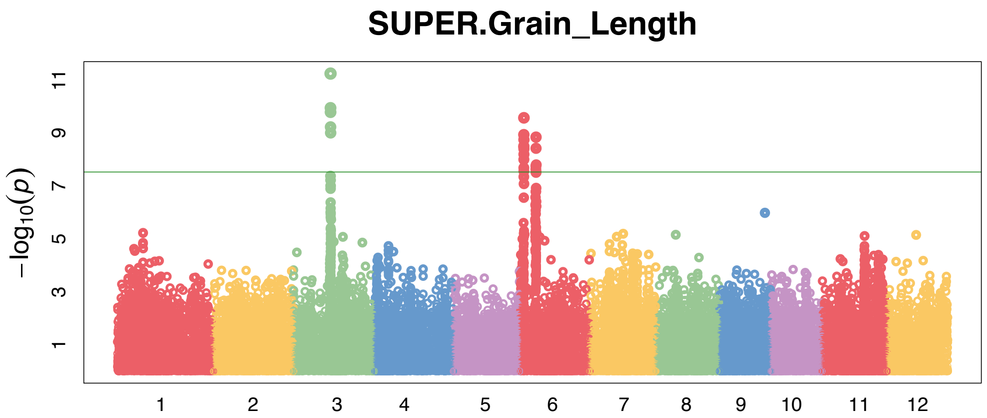


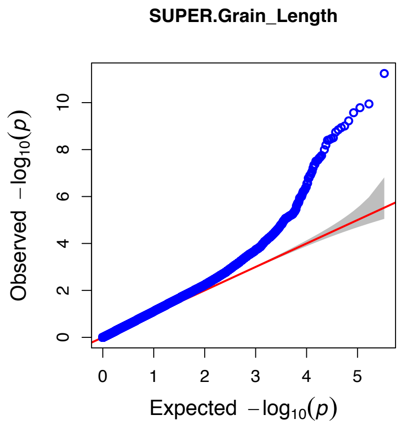


Japonica Panel
211 samples
178 phenotypes

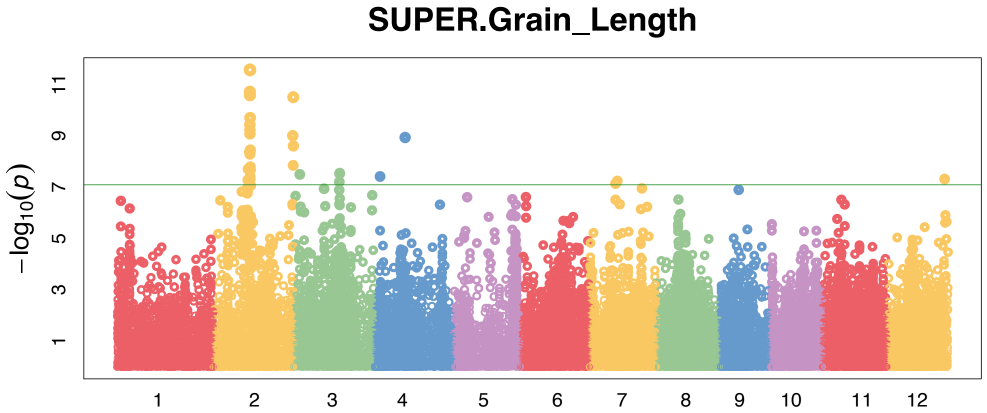


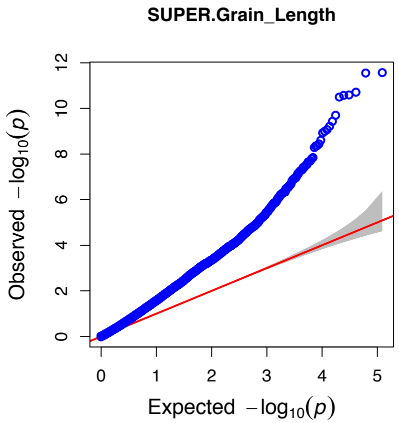

Full Panel
672 samples
503 phenotypes

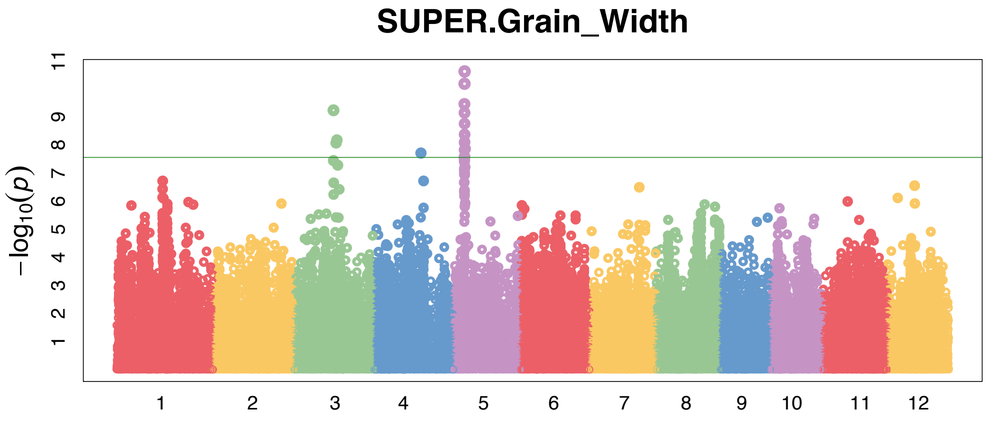


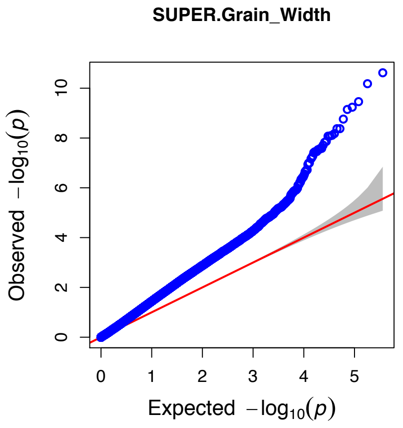


Indica Panel
426 samples
297 phenotypes

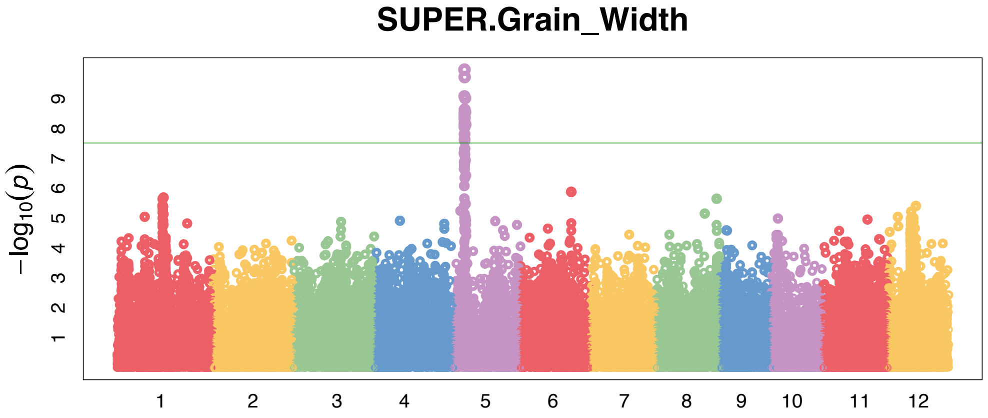


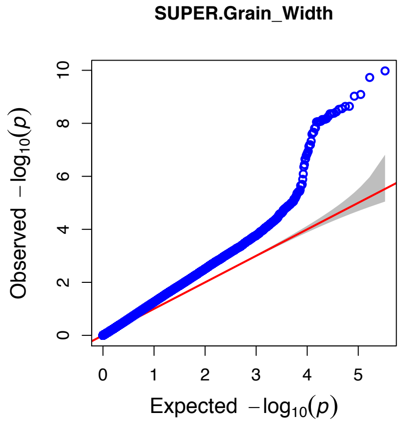


Japonica Panel
211 samples
178 phenotypes

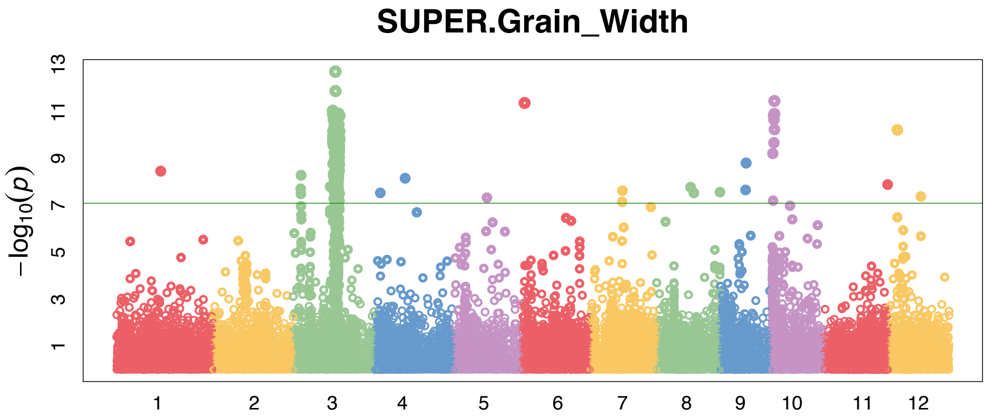


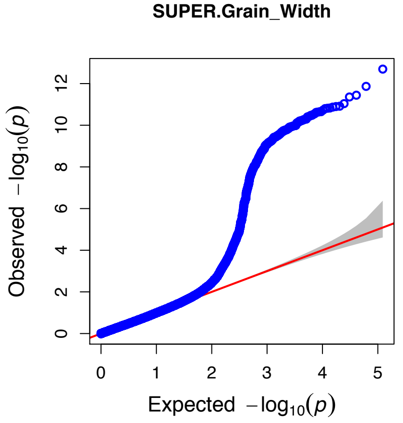

Full Panel
672 samples
500 phenotypes

Indica Panel
426 samples
295 phenotypes

Japonica Panel
211 samples
177 phenotypes

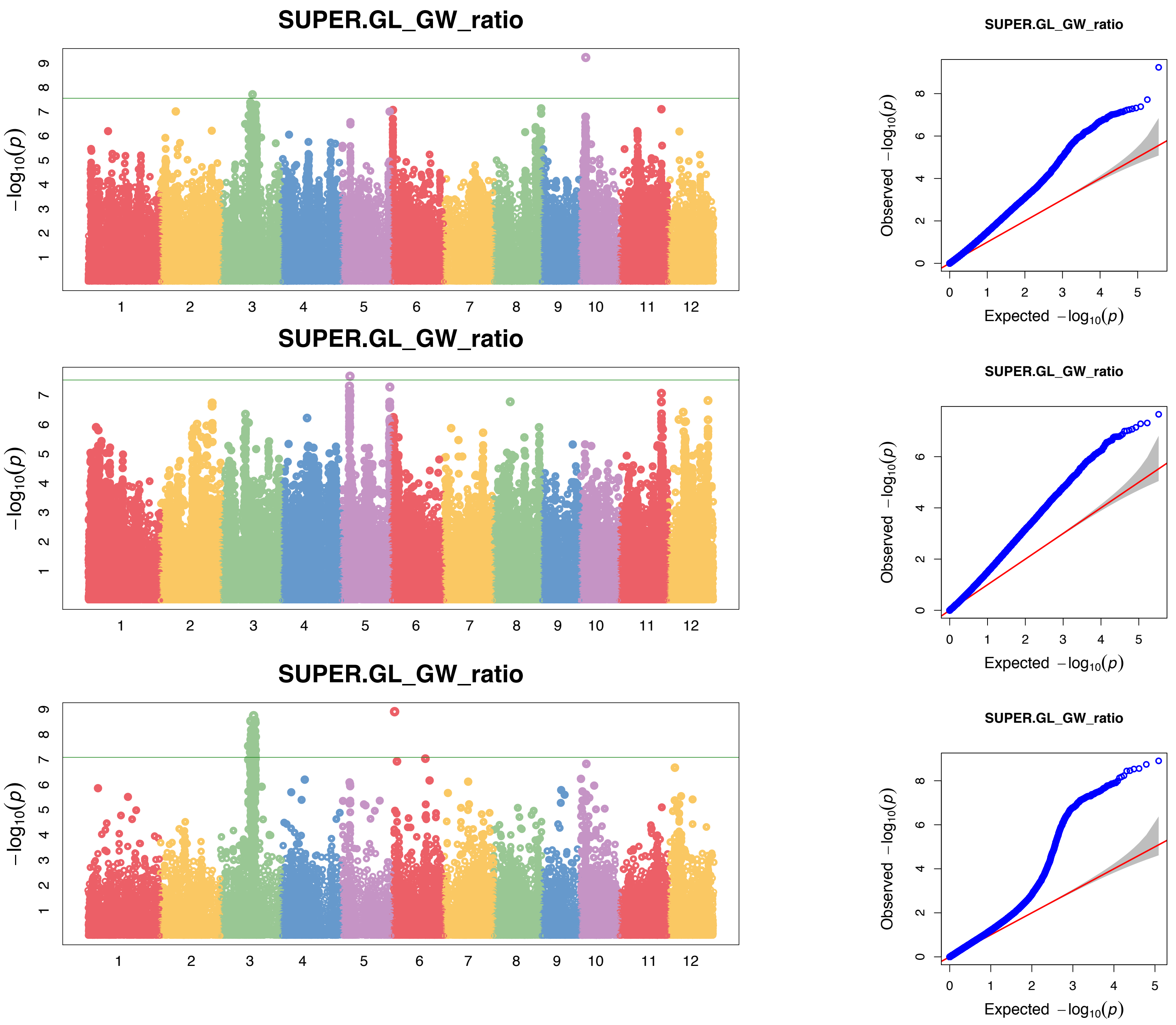

Full Panel
672 samples
486 phenotypes

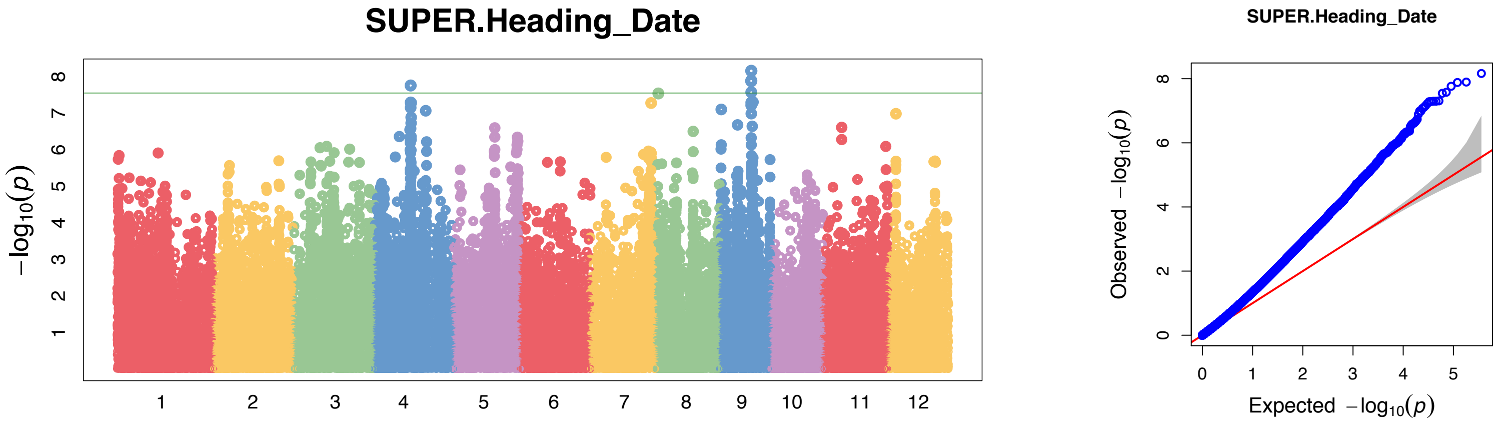


Indica Panel
426 samples
286 phenotypes

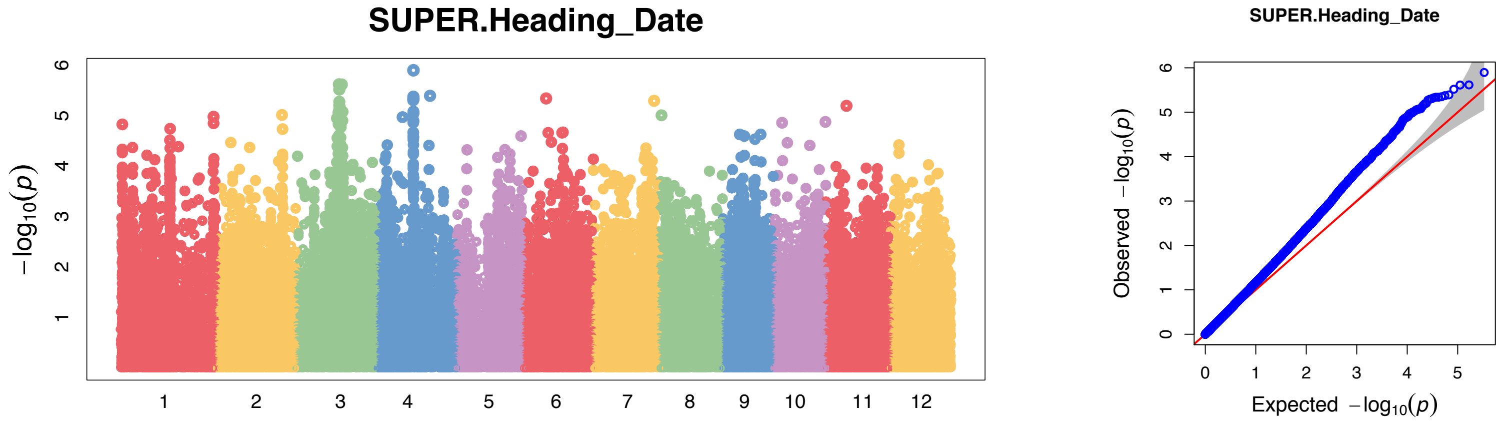


Japonica Panel
211 samples
172 phenotypes

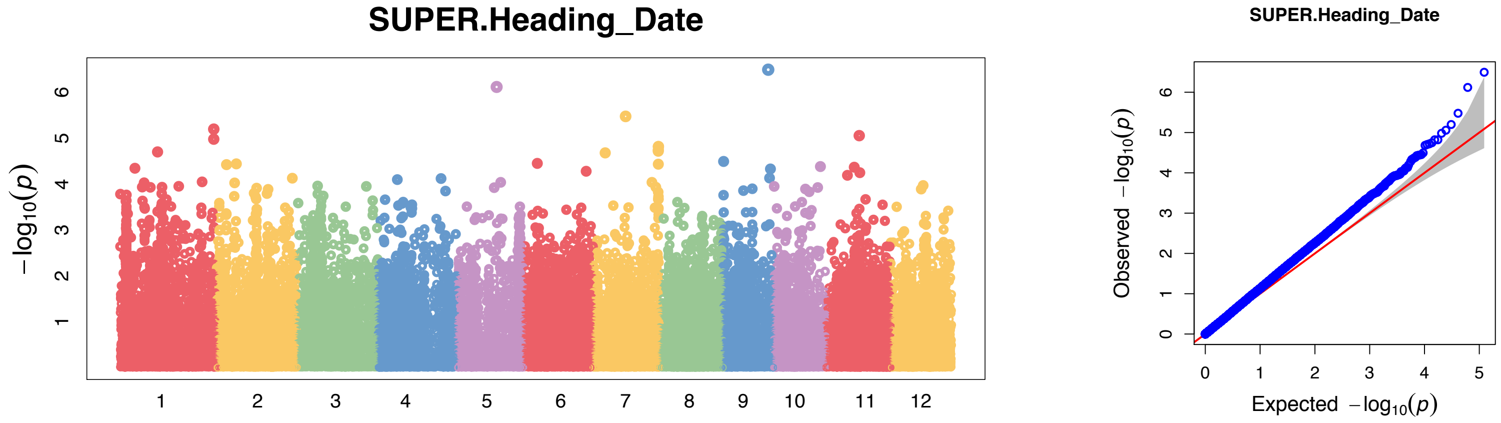

Full Panel
672 samples
452 phenotypes

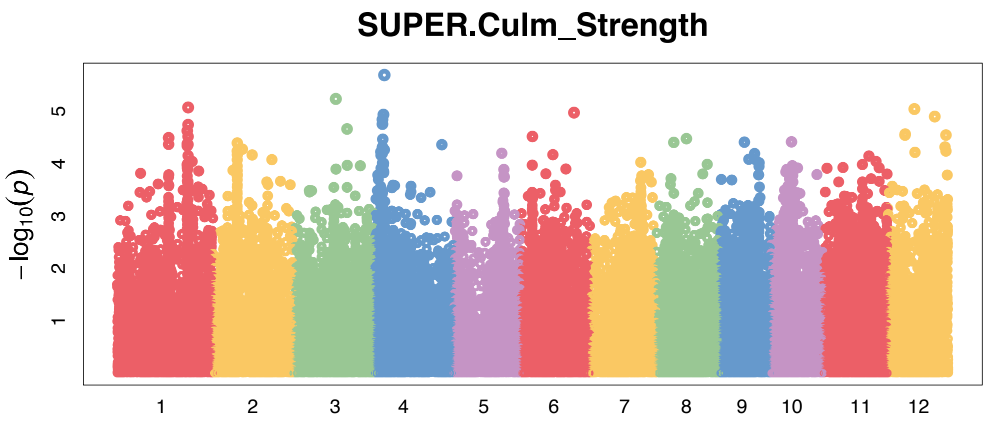


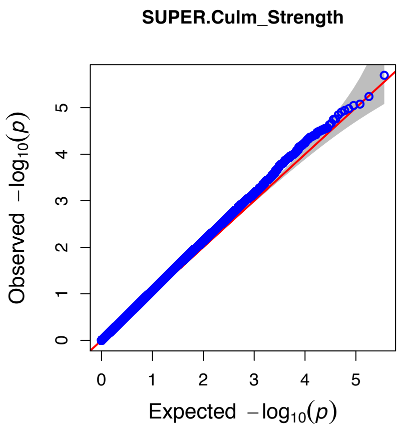


Indica Panel
426 samples
254 phenotypes

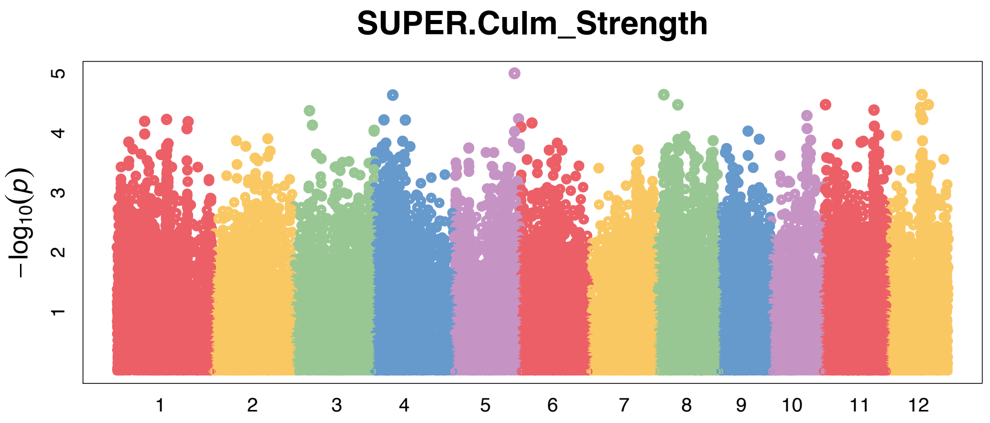


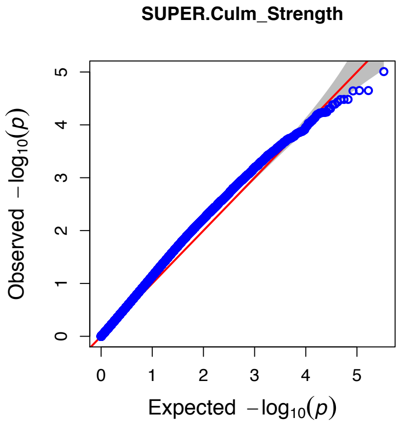


Japonica Panel
211 samples
170 phenotypes

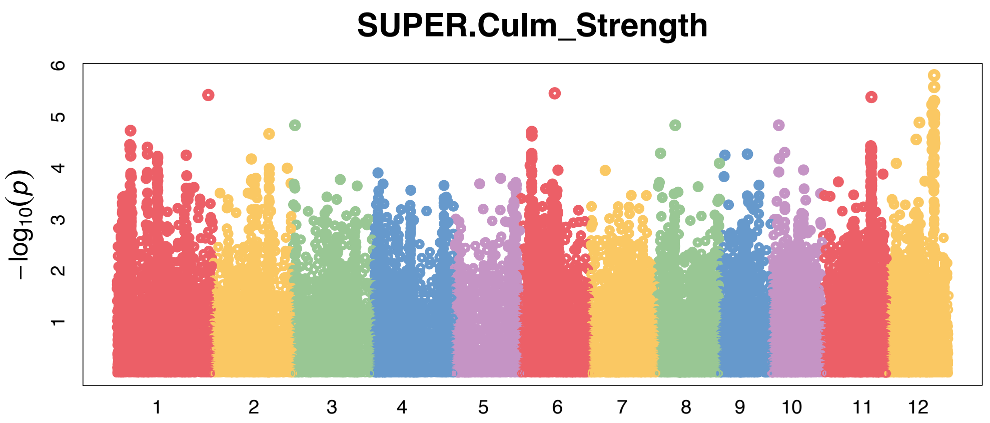


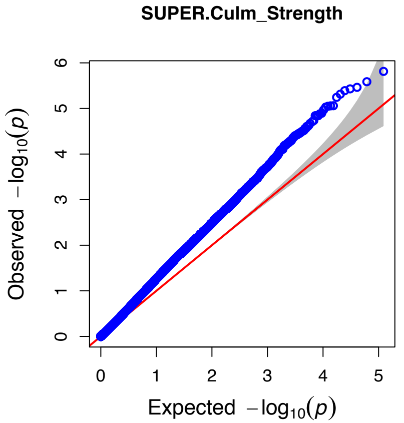

Full Panel
672 samples
356 phenotypes

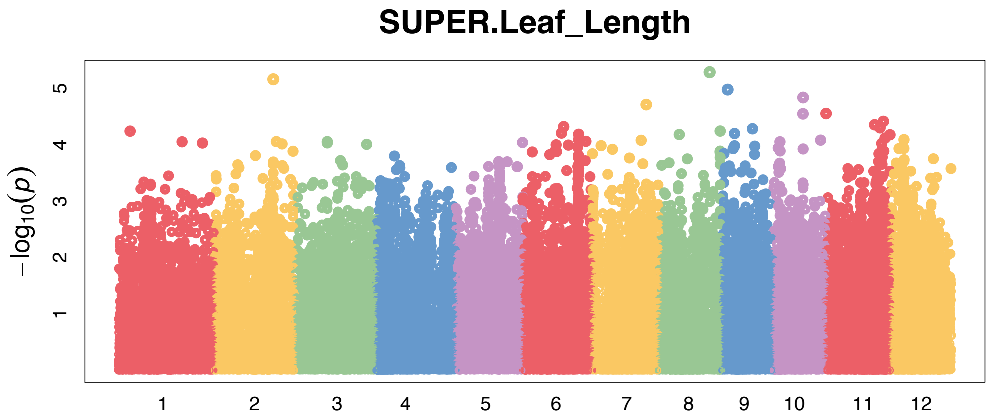


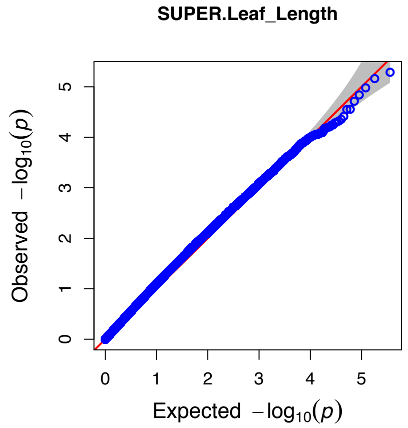


Indica Panel
426 samples
195 phenotypes

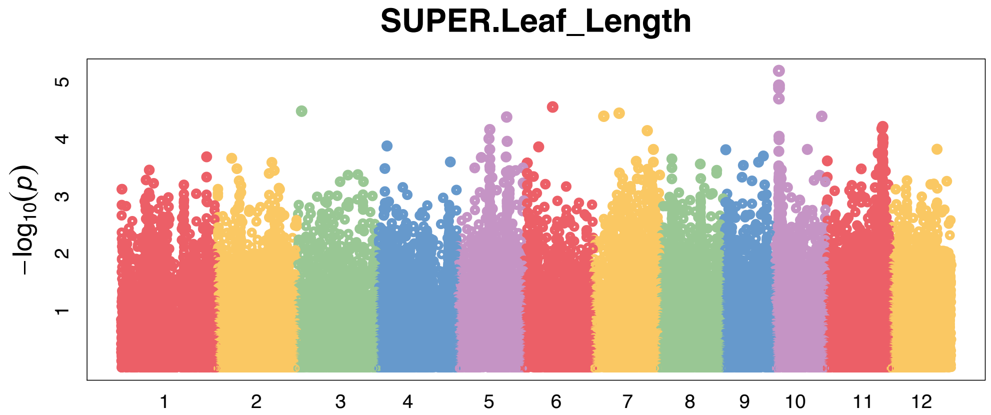


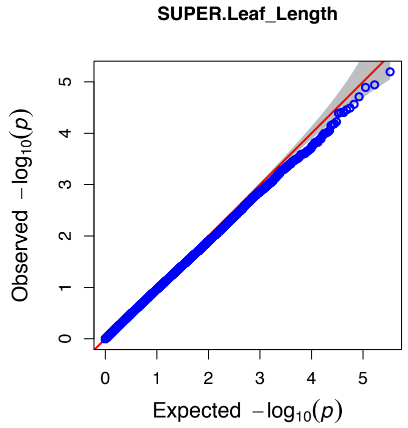


Japonica Panel
211 samples
137 phenotypes

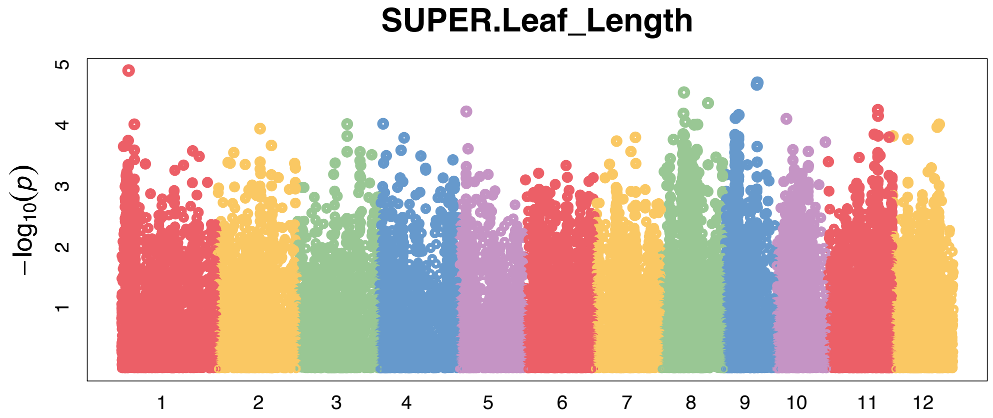


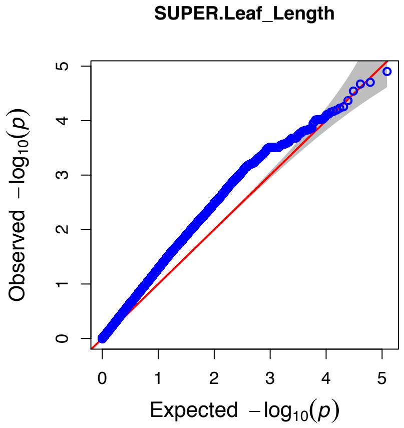

Full Panel
672 samples
355 phenotypes

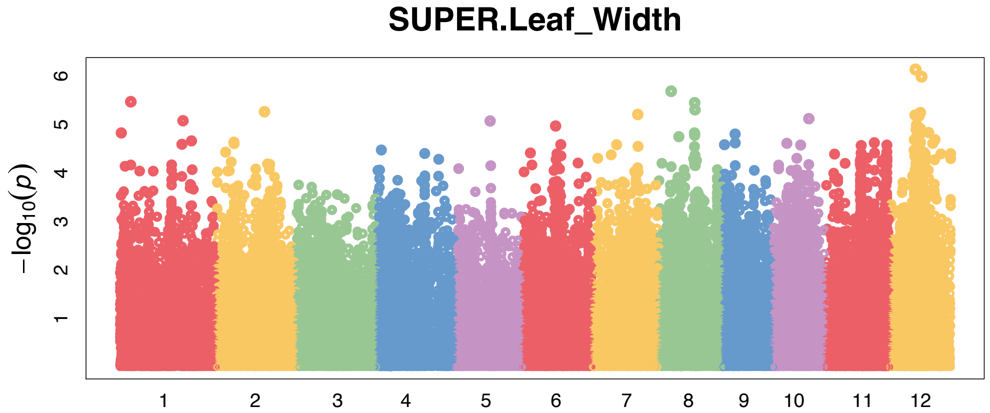


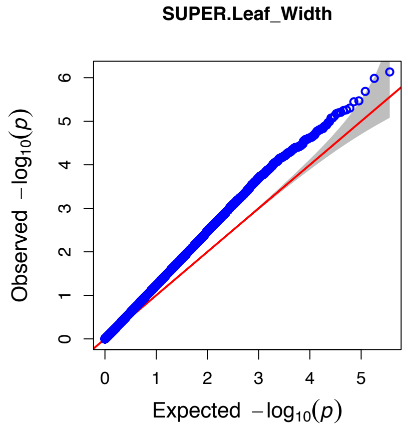


Indica Panel
426 samples
196 phenotypes

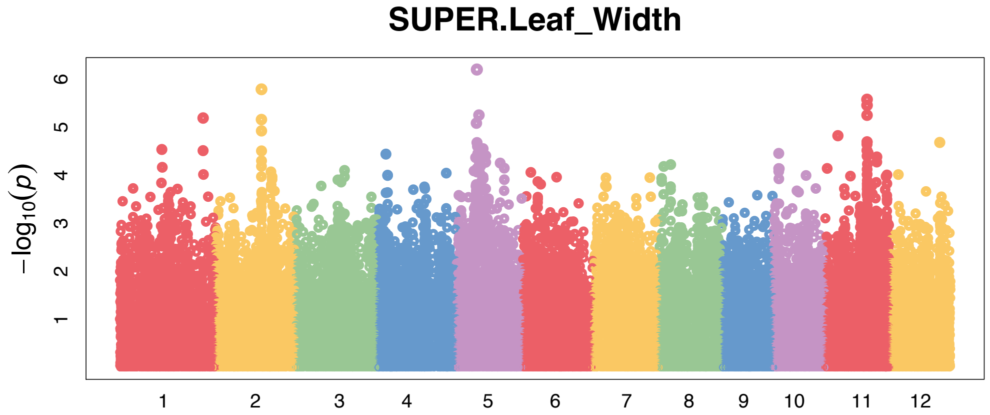


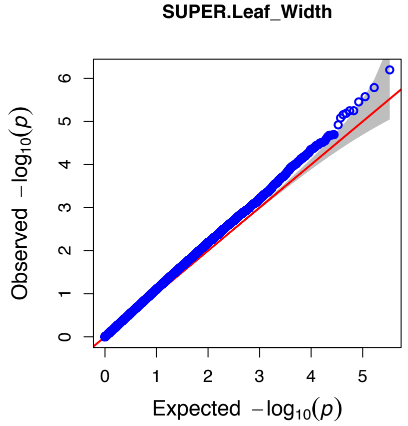


Japonica Panel
211 samples
136 phenotypes

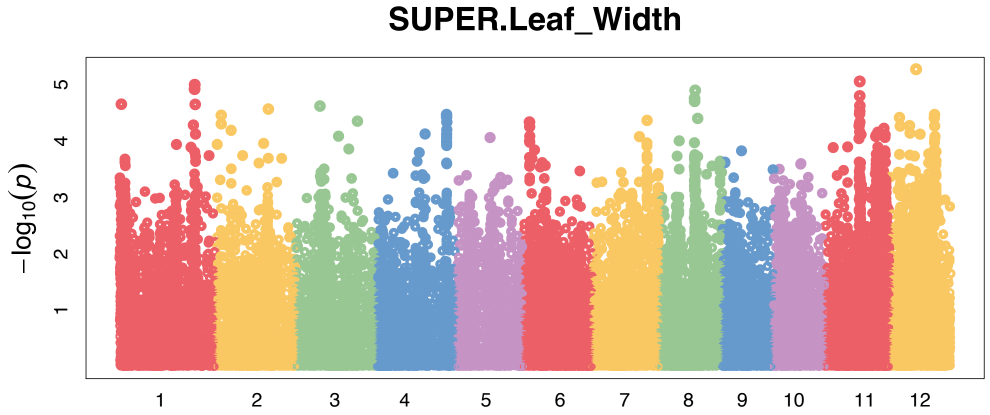


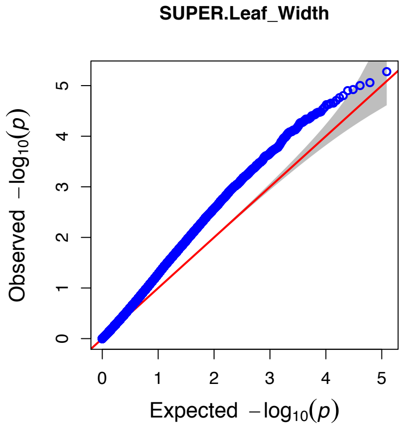
