## Supplementary material for "Evidence of selection, adaptation and untapped diversity in Vietnamese rice landraces": Table S20. Summary count of SNPs with effects on the genome (using MSU7 annotation)

| **Table S20. Summary count of SNPs with effects on the genome (using MSU7 annotation).** | | |  |  |
| --- | --- | --- | --- | --- |
| **Table S20.1 Number of effects by impact** |  |  |  |  |
| Impact type | Count | Percent |  |  |
| High-effect SNP | 21,639 | 0.197% |  |  |
| Low-effect SNP | 237,942 | 2.164% |  |  |
| Moderate-effect SNP | 304,612 | 2.770% |  |  |
| Modifier SNP | 10,430,885 | 94.869% |  |  |
| **Table S20.2 Number of effects by functional class** |  |  |  |  |
| Type | Count | Percent |  |  |
| MISSENSE | 306,492 | 57.89% |  |  |
| NONSENSE | 16,212 | 3.06% |  |  |
| SILENT | 206,778 | 39.05% |  |  |
| Missense / Silent ratio: 1.4822 |  |  |  |  |
| **Table S20.3 Number of effects by Type** |  |  |  |  |
| Type (alphabetical order) | Count | Percent | category |  |
| 3_prime_UTR_variant | 119,154 | 1.08% | MODIFIER |  |
| 5_prime_UTR_premature_start_codon_gain_variant | 8,111 | 0.07% | LOW |  |
| 5_prime_UTR_variant | 51,427 | 0.47% | MODIFIER |  |
| downstream_gene_variant | 3,333,691 | 30.23% | MODIFIER |  |
| initiator_codon_variant | 49 | 0% | LOW |  |
| intergenic_region | 2,515,136 | 22.81% | MODIFIER |  |
| intragenic_variant | 411 | 0.00% | MODIFIER |  |
| intron_variant | 836,876 | 7.59% | MODIFIER |  |
| missense_variant | 304,612 | 2.76% | MODERATE |  |
| non_coding_transcript_variant | 34 | 0% | MODIFIER |  |
| splice_acceptor_variant | 1,897 | 0.02% | HIGH |  |
| splice_donor_variant | 1,699 | 0.02% | HIGH |  |
| splice_region_variant | 31,629 | 0.29% | HIGH |  |
| start_lost | 741 | 0.01% | HIGH |  |
| stop_gained | 16,212 | 0.15% | HIGH |  |
| stop_lost | 1,090 | 0.01% | HIGH |  |
| stop_retained_variant | 397 | 0.00% | LOW |  |
| synonymous_variant | 206,381 | 1.87% | LOW |  |
| upstream_gene_variant | 3,599,233 | 32.64% | MODIFIER |  |
| **Table S20.4 Number of effects by Region** |  |  |  |  |
| Region | Count | Percent |  |  |
| DOWNSTREAM | 3,333,691 | 30.32% |  |  |
| EXON | 526,138 | 4.79% |  |  |
| INTERGENIC | 2,515,136 | 22.88% |  |  |
| INTRON | 811,799 | 7.38% |  |  |
| SPLICE_SITE_ACCEPTOR | 1,897 | 0.02% |  |  |
| SPLICE_SITE_DONOR | 1,699 | 0.02% |  |  |
| SPLICE_SITE_REGION | 26,348 | 0.24% |  |  |
| TRANSCRIPT | 445 | 0.00% |  |  |
| UPSTREAM | 3,599,233 | 32.74% |  |  |
| UTR_3_PRIME | 119,154 | 1.08% |  |  |
| UTR_5_PRIME | 59,538 | 0.54% |  |  |
| **Table S20.5 Base changes** |  |  |  |  |
|  | **A** | **C** | **G** | **T** |
| **A** | 0 | 107,604 | 453,160 | 150,533 |
| **C** | 174,709 | 0 | 101,463 | 885,652 |
| **G** | 887,039 | 101,426 | 0 | 175,869 |
| **T** | 151,574 | 452,967 | 108,625 | 0 |
| **Table S20.6 Ts/Tv** |  |  |  |  |
| Transitions | 1,084,607,622 |  |  |  |
| Transversions | 440,158,145 |  |  |  |
| Ts/Tv ratio | 2.4641 |  |  |  |
