## Supplementary material for "Evidence of selection, adaptation and untapped diversity in Vietnamese rice landraces": Table S10. Summary of XP-CLR comparisons for the Indica and Japonica subpopulations

| **Table S10. Summary of XP-CLR comparisons for the Indica and Japonica subpopulations.** | | | | | | |  |  |  |  |  |
| --- | --- | --- | --- | --- | --- | --- | --- | --- | --- | --- | --- |
| xp-clr A (selected) | xp-clr B | mean xp_clr score | cutoff (99 percentile) | cutoff score for selecting regions | number of selected regions | merged regions -d 100000 | filtered regions >80000 | mean length | total length | % of genome 373,245,519 bp | number of genes |
| **Japonica** | | | | | | | | | | | |
| J1 | J2 | 17.80 | 223 | 136 | 1151 | 181 | 179 | 264,874 | 47,412,524 |  |  |
| J1 | J3 | 7.64 | 108 | 136 | 187 | 28 | 28 | 259,695 | 7,271,469 |  |  |
| J1 | J4 | 6.13 | 76 | 136 | 33 | 16 | 16 | 129,925 | 2,078,792 |  |  |
| J1 merge |  | 10.52 | 136 |  |  |  | 28 | 576,707 | 16,147,785 | 4.33 | 2427 |
| J2 | J1 | 49.54 | 487 | 256 | 2074 | 230 | 227 | 323,361 | 73,402,859 |  |  |
| J2 | J3 | 21.62 | 200 | 256 | 137 | 30 | 30 | 237,042 | 7,111,263 |  |  |
| J2 | J4 | 6.65 | 80 | 256 | 0 | 0 | 0 |  |  |  |  |
| J2 merge |  | 25.94 | 256 |  |  |  | 23 | 726,689 | 16,713,841 | 4.48 | 2439 |
| J3 | J1 | 24.42 | 365 | 228 | 887 | 109 | 108 | 292,759 | 31,617,951 |  |  |
| J3 | J2 | 17.92 | 241 | 228 | 407 | 66 | 64 | 246,651 | 15,785,683 |  |  |
| J3 | J3 | 5.91 | 79 | 228 | 7 | 3 | 2 | 128,661 | 257,321 |  |  |
| J3 merge |  | 16.08 | 228 |  |  |  | 24 | 577,089 | 13,850,139 | 3.71 | 2007 |
| J4 | J1 | 46.12 | 477 | 297 | 1502 | 160 | 159 | 331,402 | 52,692,947 |  |  |
| J4 | J2 | 17.45 | 221 | 297 | 125 | 36 | 34 | 179,487 | 6,102,559 |  |  |
| J4 | J3 | 17.85 | 192 | 297 | 62 | 15 | 15 | 213,497 | 3,202,452 |  |  |
| J4 merge |  | 27.14 | 297 |  |  |  | 25 | 731,341 | 18,283,522 | 4.90 | 2643 |
| **Indica** | | | | | | | | | | | |
| I1 | I2 | 8.51 | 179 | 161 | 453 | 110 | 109 | 199,615 | 21,757,988 |  |  |
|  | I3 | 4.00 | 84 | 161 | 94 | 35 | 35 | 156,335 | 5,471,709 |  |  |
|  | I4 | 8.17 | 169 | 161 | 397 | 111 | 110 | 182,287 | 20,051,616 |  |  |
|  | I5 | 9.79 | 211 | 161 | 569 | 120 | 119 | 233,178 | 27,748,233 |  |  |
| merge |  | 7.62 | 161 |  |  |  | 44 | 453,570 | 19,957,065 | 5.35 | 3077 |
| I2 | I1 | 28.81 | 410 | 275 | 1104 | 197 | 195 | 250,882 | 48,922,046 |  |  |
|  | I3 | 7.03 | 114 | 275 | 23 | 11 | 11 | 124,789 | 1,372,675 |  |  |
|  | I4 | 15.65 | 286 | 275 | 399 | 104 | 104 | 196,457 | 20,431,532 |  |  |
|  | I5 | 17.30 | 291 | 275 | 425 | 114 | 113 | 180,500 | 20,396,496 |  |  |
| merge |  | 17.20 | 275 |  |  |  | 41 | 550,836 | 22,584,270 | 6.05 | 3346 |
| I3 | I1 | 40.21 | 492 | 401 | 739 | 147 | 146 | 227,018 | 33,144,691 |  |  |
|  | I2 | 24.53 | 361 | 401 | 269 | 79 | 79 | 165,492 | 13,073,845 |  |  |
|  | I4 | 23.15 | 374 | 401 | 309 | 75 | 75 | 196,812 | 14,760,892 |  |  |
|  | I5 | 23.66 | 375 | 401 | 297 | 70 | 69 | 206,996 | 14,282,693 |  |  |
| merge |  | 27.89 | 401 |  |  |  | 42 | 474,009 | 19,908,387 | 5.33 | 2993 |
| I4 | I1 | 34.07 | 462 | 306 | 1099 | 183 | 183 | 247,829 | 45,352,723 |  |  |
|  | I2 | 21.46 | 343 | 306 | 492 | 96 | 96 | 217,528 | 20,882,658 |  |  |
|  | I3 | 7.44 | 119 | 306 | 24 | 8 | 8 | 166,618 | 1,332,945 |  |  |
|  | I5 | 18.62 | 300 | 306 | 349 | 85 | 85 | 197,064 | 16,750,419 |  |  |
| merge |  | 20.40 | 306 |  |  |  | 38 | 619,404 | 23,537,343 | 6.31 | 3465 |
| I5 | I1 | 63.64 | 611 | 440 | 1255 | 208 | 207 | 267,024 | 55,274,009 | 14.81 |  |
|  | I2 | 44.58 | 462 | 440 | 471 | 120 | 120 | 204,496 | 24,539,544 | 6.57 |  |
|  | I3 | 18.01 | 233 | 440 | 58 | 15 | 14 | 162,347 | 2,272,862 | 0.61 |  |
|  | I4 | 39.21 | 453 | 440 | 423 | 123 | 122 | 184,271 | 22,481,109 | 6.02 |  |
| merge |  | 41.36 | 440 |  |  |  | 52 | 583,706 | 30,352,734 | 8.13 | 4576 |
