## Supplementary material for "Evidence of selection, adaptation and untapped diversity in Vietnamese rice landraces": Table S6. Diversity of each subpopulation

| **Table S6. Diversity** (π) **of each subpopulation** | |
| --- | --- |
| Indica subpopulation | mean Diversity (∏) |
| I1 | 0.001440 |
| I2 | 0.001269 |
| I3 | 0.001210 |
| I4 | 0.001196 |
| I5 | 0.001034 |
| Japonica subpopulation | mean Diversity (∏) |
| J1 | 0.000577 |
| J2 | 0.000535 |
| J3 | 0.000697 |
| J4 | 0.000531 |
