## Supplementary material for "Evidence of selection, adaptation and untapped diversity in Vietnamese rice landraces": Table S7. GWAS results; List of the 21 QTLs and the positions of the individual QTLs for each panel.

| **Table S7. GWAS results.** List of the 21 QTLs and the positions of the individual QTLs for each panel. | | | | | | |  |  |  |  |  |  |  |
| --- | --- | --- | --- | --- | --- | --- | --- | --- | --- | --- | --- | --- | --- |
|  |  |  |  |  |  |  |  | Details of the position with the highest significance | | | Final set of QTLs | | |
| QTL | Trait | Panel | Chromosome | start | end | length | Number of significant SNPs | max position | P.value | FDR | start (incl LD) | end (incl LD) | length |
| 1_DI | Diameter_Internode | FP | 2 | 6,805,273 | 6,923,410 | 118,137 | 3 | 6,805,273 | 3.12E-08 | 8.16E-03 | 6,805,273 | 6,923,410 | 118,137 |
| 2_GL | Grain_Length | FP | 2 | 16,368,834 | 16,496,986 | 128,152 | 6 | 16,368,834 | 1.93E-09 | 1.46E-05 | 15,480,976 | 16,798,043 | 1,317,067 |
|  | Grain_Length | Japonica | 2 | 15,480,976 | 16,798,043 | 1,317,067 | 21 | 16,406,945 | 2.69E-12 | 1.72E-07 |  |  |  |
| 3_GL_jap | Grain_Length | Japonica | 2 | 35,638,527 | 35,927,940 | 289,413 | 4 | 35,871,898 | 3.16E-11 | 6.48E-07 | 35,638,527 | 35,927,940 | 289,413 |
| 4_GW_jap | Grain_Width | Japonica | 3 | 3,334,516 | 3,532,506 | 197,990 | 3 | 3,532,506 | 5.26E-09 | 3.47E-06 | 3,334,516 | 3,532,506 | 197,990 |
| 5_GS | Grain_Length | FP | 3 | 16,676,504 | 16,908,475 | 231,971 | 20 | 16,706,516 | 9.26E-17 | 1.72E-11 | 16,520,656 | 16,908,475 | 387,819 |
|  | Grain_Length | Indica | 3 | 16,667,059 | 16,732,936 | 65,877 | 8 | 16,684,014 | 5.73E-12 | 1.92E-06 |  |  |  |
|  | Grain_Width | Japonica | 3 | 16,520,656 | 16,524,731 | 4,075 | 2 | 16,524,731 | 2.72E-09 | 2.02E-06 |  |  |  |
| 6_GS | Grain_Width | FP | 3 | 18,051,736 | 19,975,669 | 1,923,933 | 5 | 18,070,507 | 5.79E-10 | 5.15E-05 | 17,686,248 | 20,833,777 | 3,147,529 |
|  | GL_GW_ratio | FP | 3 | 17,748,797 | 20,713,844 | 2,965,047 | 12 | 18,750,652 | 9.51E-08 | 1.96E-03 |  |  |  |
|  | GL_GW_ratio | Japonica | 3 | 17,686,248 | 20,817,812 | 3,131,564 | 89 | 18,005,179 | 2.94E-09 | 7.37E-05 |  |  |  |
|  | Grain_Width | Japonica | 3 | 17,686,248 | 20,833,777 | 3,147,529 | 246 | 18,869,675 | 2.02E-13 | 2.48E-08 |  |  |  |
|  | Grain_Length | Japonica | 3 | 20,618,252 | 20,793,865 | 175,613 | 3 | 20,793,865 | 2.88E-08 | 1.53E-04 |  |  |  |
| 7_GL | Grain_Length | FP | 4 | 12,043,539 | 13,108,767 | 1,065,228 | 14 | 12,378,123 | 5.51E-11 | 5.86E-07 | 12,043,539 | 13,108,767 | 1,065,228 |
| 8_HD | Heading_Date | FP | 4 | 16,165,354 | 16,384,087 | 218,733 | 4 | 16,167,035 | 1.72E-08 | 1.56E-03 | 16,165,354 | 16,384,087 | 218,733 |
| 9_PL | Panicle_Length | FP | 5 | 717,557 | 743,760 | 26,203 | 2 | 717,557 | 6.17E-08 | 5.69E-03 | 667,557 | 767,557 | 100,000 |
| 10_GS | Grain_Width | FP | 5 | 4,802,345 | 5,237,830 | 435,485 | 21 | 4,932,498 | 2.40E-11 | 8.65E-06 | 4,802,345 | 5,383,914 | 581,569 |
|  | Grain_Width | Indica | 5 | 4,833,479 | 5,383,914 | 550,435 | 31 | 4,833,479 | 1.05E-10 | 3.11E-05 |  |  |  |
|  | GL_GW_ratio | Indica | 5 | 4,833,479 | 5,237,830 | 404,351 | 5 | 5,143,046 | 2.22E-08 | 3.77E-03 |  |  |  |
| 11_GL | Grain_Length | FP | 6 | 1,638,768 | 1,664,716 | 25,948 | 4 | 1,638,768 | 6.91E-09 | 3.84E-05 | 1,561,006 | 1,664,716 | 103,710 |
|  | Grain_Length | Indica | 6 | 1,561,006 | 1,646,729 | 85,723 | 12 | 1,620,330 | 2.68E-10 | 2.25E-05 |  |  |  |
| 12_GL | Grain_Length | FP | 6 | 6,680,831 | 7,190,137 | 509,306 | 44 | 7,051,455 | 1.81E-14 | 1.09E-09 | 6,680,831 | 7,190,137 | 509,306 |
|  | Grain_Length | Indica | 6 | 6,971,077 | 7,142,806 | 171,729 | 7 | 7,050,752 | 1.44E-09 | 6.02E-05 |  |  |  |
| 13_GL | Grain_Length | FP | 6 | 7,503,914 | 7,560,865 | 56,951 | 2 | 7,503,914 | 5.90E-08 | 2.29E-04 | 7,453,914 | 7,553,914 | 100,000 |
| 14_PL | Panicle_Length | FP | 6 | 20,450,110 | 20,485,120 | 35,010 | 2 | 20,450,110 | 2.72E-08 | 5.69E-03 | 20,400,110 | 20,500,110 | 100,000 |
| 15_GL_jap | Grain_Length | Japonica | 7 | 11,519,294 | 12,296,525 | 777,231 | 3 | 12,296,525 | 5.76E-08 | 2.28E-04 | 11,519,294 | 12,296,525 | 777,231 |
| 16_FP | Floret_Pubescence | FP | 8 | 18,026,668 | 18,054,654 | 27,986 | 2 | 18,054,654 | 1.64E-08 | 1.35E-04 | 18,004,654 | 18,104,654 | 100,000 |
| 17_FP | Floret_Pubescence | FP | 8 | 26,225,268 | 26,265,955 | 40,687 | 2 | 26,225,268 | 6.06E-08 | 3.84E-04 | 26,175,268 | 26,275,268 | 100,000 |
| 18_FP | Floret_Pubescence | FP | 9 | 6,656,837 | 7,940,621 | 1,283,784 | 51 | 7,764,859 | 7.23E-12 | 1.76E-06 | 6,656,837 | 7,940,621 | 1,283,784 |
| 19_HD | Heading_Date | FP | 9 | 14,067,272 | 14,807,406 | 740,134 | 7 | 14,807,406 | 6.86E-09 | 1.56E-03 | 14,067,272 | 14,807,406 | 740,134 |
| 20_GW_jap | Grain_Width | Japonica | 10 | 1,098,998 | 1,404,807 | 305,809 | 6 | 1,404,807 | 3.61E-12 | 1.35E-07 | 1,098,998 | 1,404,807 | 305,809 |
| 21_LW | Leaf_width | FP | 12 | 17,445,137 | 17,511,823 | 66,686 | 2 | 17,511,823 | 2.14E-09 | 7.74E-04 | 17,445,137 | 17,561,823 | 116,686 |
