## Supplementary material for "Evidence of selection, adaptation and untapped diversity in Vietnamese rice landraces": Table S5. Phenotype statistics (mean and coefficient of variation) and population comparisons

| **Table S5. Phenotype statistics (mean and coefficient of variation) and population comparisons.** | | | | | | |  |  |  |  |  |  |  |
| --- | --- | --- | --- | --- | --- | --- | --- | --- | --- | --- | --- | --- | --- |
| **Phenotype abbrevation** | http://csdl.prc.org.vn/ | IRRI | viet672 | | | ind426 | | | jap211 | | | T-test Ind Jap ^ | *T-test irri_viet |
|  |  |  | Number of samples | mean | CV | Number of samples | mean | CV | Number of samples | mean | CV |  |  |
| Grain_Length | Yes | grlt | 501 | 8.8 | 10.5 | 295 | 8.8 | 10.1 | 178 | 9.0 | 11.0 | 1.85E-02 | 0.7626 |
| Grain_Width | Yes | grwd | 503 | 3.0 | 15.3 | 297 | 2.8 | 13.5 | 178 | 3.3 | 11.4 | **2.20E-16** | 0.6272 |
| GL_GW_ratio |  |  | 500 | 3.0 | 20.0 | 295 | 3.2 | 19.0 | 177 | 2.7 | 18.0 | **1.61E-15** | 0.4826 |
| Heading_Date | Yes | hdg_80head | 486 | 129.6 | 14.5 | 286 | 128.5 | 16.8 | 172 | 129.1 | 8.5 | 7.22E-01 | 0.9366 |
| Culm_Strength |  |  | 452 | 4.8 | 55.2 | 254 | 5.2 | 56.2 | 170 | 4.3 | 50.1 | 2.69E-04 |  |
| Panicle_Exsertion |  |  | 455 | 1.9 | 66.2 | 256 | 2.1 | 60.4 | 171 | 1.7 | 67.9 | 5.13E-04 |  |
| Leaf_Length |  | llt | 356 | 48.7 | 21.0 | 195 | 50.0 | 23.4 | 137 | 46.7 | 16.6 | 4.05E-03 | 0.09754 |
| Leaf_Width |  | lwd | 355 | 1.3 | 19.9 | 196 | 1.2 | 21.3 | 136 | 1.4 | 15.1 | **1.64E-11** | 0.592 |
| Leaf_Pubescence |  |  | 328 | 2.1 | 34.4 | 170 | 2.3 | 25.3 | 134 | 1.8 | 42.2 | **1.74E-11** |  |
| Leaf_Angle |  | la | 462 | 3.2 | 70.5 | 260 | 3.0 | 78.3 | 174 | 3.6 | 56.4 | 1.06E-02 | 0.7367 |
| Flag_Leaf_Angle |  | fla_repro | 497 | 3.9 | 58.9 | 291 | 3.1 | 67.0 | 178 | 5.2 | 39.4 | **2.20E-16** | 0.6011 |
| Culm_Number | Yes |  | 454 | 5.7 | 31.6 | 254 | 6.3 | 28.3 | 172 | 4.9 | 29.8 | **9.52E-16** |  |
| Culm_Angle |  |  | 475 | 2.0 | 72.8 | 269 | 2.0 | 79.1 | 178 | 2.0 | 63.6 | 7.91E-01 |  |
| Diameter_Internode |  | cudi_repro | 485 | 4.3 | 24.3 | 284 | 4.2 | 26.6 | 174 | 4.4 | 21.2 | 8.99E-03 | 0.07265 |
| Culm_Length | Yes | cult_repro | 485 | 103.9 | 23.5 | 282 | 107.6 | 26.9 | 175 | 98.2 | 14.6 | **7.19E-05** | 0.6965 |
| Panicle_Length |  | plt_post | 486 | 25.6 | 10.9 | 283 | 24.9 | 10.3 | 175 | 26.4 | 11.0 | **3.12E-09** | 0.2416 |
| Panicle_Type |  |  | 455 | 4.6 | 58.9 | 254 | 4.7 | 58.6 | 173 | 4.4 | 60.9 | 3.15E-01 |  |
| Awning |  |  | 463 | 0.6 | 290.8 | 262 | 0.3 | 351.7 | 173 | 1.0 | 242.3 | 1.52E-04 |  |
| Floret_Colour |  |  | 467 | 1.7 | 118.5 | 264 | 1.1 | 121.2 | 175 | 2.4 | 103.4 | **3.50E-11** |  |
| Floret_Pubescence |  | lppub | 488 | 3.3 | 42.5 | 284 | 3.8 | 28.6 | 176 | 2.6 | 60.4 | **2.20E-16** | 0.7461 |
| CV Coefficient of variation | |  |  |  |  |  |  |  |  |  |  |  |  |
| *compare subpopulation I2 (IRRI 14 samples, our dataset 68 samples) | | | | |  |  |  |  |  |  |  |  |  |
| ^ 10 traits were significantly different between the Indica and Japonica subtypes (p-value < 0.0001) | | | | | | |  |  |  |  |  |  |  |
