## Supplementary material for "Evidence of selection, adaptation and untapped diversity in Vietnamese rice landraces": Table S4. Phenotyping abbreviations and details

| **Table S4. Phenotype abbreviations and details.** | |  |  |  |  |  |  |  |  |  |  |  |  |  |  |  |  |  |  |
| --- | --- | --- | --- | --- | --- | --- | --- | --- | --- | --- | --- | --- | --- | --- | --- | --- | --- | --- | --- |
| **Phenotype abbrevation** | **Heading_Date** | **Culm_Strength** | **Panicle_Exsertion** | **Leaf_Length** | **Leaf_Width** | **Leaf_Pubescence** | **Leaf_Angle** | **Flag_Leaf_Angle** | **Culm_Number** | **Culm_Angle** | **Diameter_Internode** | **Culm_Length** | **Panicle_Length** | **Panicle_Type** | **Awning** | **Floret_Colour** | **Floret_Pubescence** | **Grain_Length** | **Grain_Width** |
| Phenotype Details | Maturity (Date) | Culm Strength (Cs) | Panicle Exsertion (Exs) | Leaf Length (LL) - cm | Leaf Width (LW) - cm | Leaf Blade Pubescence (LBP) | Leaf Angle (LA) | Flag Leaf Angle (FLA) | Culm Number (CmN) | Culm Angle (CmA) | Diameter of Basal Internode (DBI) - mm | Culm Length (Cml) - cm | Panicle Length (PnL) - cm | Panicle Type (PnT) | Awning (An) | Lemma and Palea Color (LmPC) | Lemma and Palea Pubescence (LmPb) | Grain Length (GrL) - mm | Grain Width (GrW) - mm |
| Phenotype Codes |  | 1 - Strong (no bending) | 1 - Well exserted |  |  | 1 - Glabrous | 1 - Erect | 1 - Erect |  | 1 - Erect, < 30^0^ |  |  |  | 1 - Compact | 0 - Absent | 0 - Straw | 1 - Glabrous |  |  |
|  |  | 3 - Moderately strong (most plants bending) | 3 - Moderately difficult (1-5%) |  |  | 2 - Intermediate | 5 - Horizontal | 3 - Intermediate |  | 3 - Intermediate, = 45^0^ |  |  |  | 7 - Intermediate | 1 - Short and partly awned | 1 - Gold and gold furrows on straw background | 2 - Hairs on lemma keel |  |  |
|  |  | 5- Intermediate (most plants moderately bending) | 7 - Partly exserted |  |  | 3 - Pubescent |  | 5 - Horizontal |  | 5 - Open, = 60^0^ |  |  |  | 9 - Open | 9 - Long and fully awned | 2 - Brown spots on straw | 3 - Hairs on upper portion |  |  |
|  |  | 7- Weak (most plants nearly flat) |  |  |  |  |  | 7 - Descending |  |  |  |  |  |  |  | 3 - Brown furrows on straw | 4 - Short hairs |  |  |
|  |  | 9 - Very weak (all plants flat) |  |  |  |  |  |  |  |  |  |  |  |  |  | 4 - Brown (tawny) | 5 - Long hairs (velvety) |  |  |
|  |  |  |  |  |  |  |  |  |  |  |  |  |  |  |  | 5 - Reddish to light purple |  |  |  |
|  |  |  |  |  |  |  |  |  |  |  |  |  |  |  |  | 6 - Purple spots on straw |  |  |  |
| Equivalent phenotype in 3K RGP study | hdg_80head |  |  | llt | lwd |  | la | fla_repro |  |  | cudi_repro | cult_repro | plt_post |  |  |  | lppub | grlt | grwd |
| Details of phenotype in 3K RGP study | Number of days at 80% fully headed |  |  | Leaf length (cm) - cultivated | Leaf width (cm) - cultivated |  | Leaf blade attitude angle of the penultimate leaf prior to heading | Flagleaf (attitude of blade ) - Late observation |  |  | Culm diameter (mm) of basal internode at repro. | Culm length | Panicle length (cm) at post-harvest |  |  |  | Lemma and palea pubescence | Grain length (mm) | Grain Width(mm) |
