## Supplementary material for "Evidence of selection, adaptation and untapped diversity in Vietnamese rice landraces": Table S15. Overlap of the selected regions in Japonica subpopulations with QTL found in Vietnamese rice datasets

| **Table S15. Overlap of the selected regions in Japonica subpopulations with QTL found in Vietnamese rice datasets.** | | | | | | | | |  |  |  |  |  |  |  |  |  |  |
| --- | --- | --- | --- | --- | --- | --- | --- | --- | --- | --- | --- | --- | --- | --- | --- | --- | --- | --- |
| Japonica J1 | | | |  | Japonica J2 | | | |  | Japonica J3 | | | |  | Japonica J4 | | | |
| chrom | start | end | QTL |  | chrom | start | end | QTL |  | chrom | start | end | QTL |  | chrom | start | end | QTL |
| 1 | 5796804 | 6558683 | Rq73_F_I_DWB60_DRW,Rq84_F_I_DRP |  | 1 | 655932 | 1078100 | pQTL_1_I_PBN |  | 1 | 5930743 | 6679572 | Rq84_F_I_DRP |  | 1 | 5760002 | 6627165 | Rq73_F_I_DWB60_DRW,Rq84_F_I_DRP,pQTL_2_F_SpN |
| 1 | 31320515 | 32167518 | . |  | 2 | 2261095 | 2479861 | . |  | 1 | 12650333 | 12987303 | . |  | 2 | 16420678 | 17573812 | 2_GL,pQTL_9_F_SBN_SpN |
| 2 | 12640224 | 13619048 | . |  | 2 | 16330202 | 17569375 | 2_GL,pQTL_9_F_SBN_SpN |  | 1 | 31910287 | 32488909 | . |  | 3 | 6197552 | 7269261 | . |
| 2 | 16420958 | 17219685 | 2_GL,pQTL_9_F_SBN_SpN |  | 2 | 17776877 | 18409421 | . |  | 1 | 40374534 | 40488895 | . |  | 3 | 11201170 | 11828657 | . |
| 4 | 1121684 | 1365036 | . |  | 3 | 19670020 | 20069317 | 6_GS |  | 2 | 2051414 | 2449915 | . |  | 4 | 8026541 | 9085022 | . |
| 4 | 1698255 | 2049982 | . |  | 4 | 8026541 | 9085022 | . |  | 2 | 16343055 | 17139837 | 2_GL,pQTL_9_F_SBN_SpN |  | 4 | 9230994 | 9745983 | . |
| 4 | 3881392 | 4049745 | . |  | 4 | 9230994 | 9745983 | . |  | 3 | 6234792 | 7678879 | . |  | 4 | 9900227 | 10128449 | . |
| 4 | 7080014 | 7219727 | . |  | 4 | 9900227 | 10128449 | . |  | 3 | 10320001 | 11828657 | . |  | 4 | 10283341 | 11184521 | . |
| 4 | 8210733 | 9771609 | . |  | 4 | 10241140 | 10929960 | . |  | 4 | 9900227 | 10144277 | . |  | 5 | 5890466 | 6217179 | . |
| 4 | 9910401 | 11208944 | . |  | 4 | 12930032 | 13529165 | 7_GL,Rq80_F_SRP |  | 4 | 10390007 | 11278312 | . |  | 5 | 15870459 | 16119560 | Tq4_F_I_Score4 |
| 7 | 10390449 | 11279603 | . |  | 4 | 13790399 | 14358393 | . |  | 6 | 7853230 | 8318785 | Rq2_J_LLGHT |  | 7 | 10680586 | 12129983 | 15_GL_jap |
| 7 | 15961245 | 17119673 | . |  | 4 | 17995330 | 18598255 | . |  | 6 | 17660650 | 18077384 | L10_I_FW_TW,Tq5_F_RCGR |  | 7 | 12637200 | 13169862 | . |
| 7 | 20690044 | 20969695 | Rq12_F_I_TIL,Tq8_F_I_RWC_T2_Score1 |  | 5 | 15580524 | 16139206 | Tq4_F_I_Score4 |  | 7 | 12471401 | 12569984 | . |  | 7 | 19150674 | 19499804 | . |
| 7 | 25060419 | 25378844 | . |  | 5 | 27331513 | 28189811 | . |  | 7 | 24316306 | 24769551 | . |  | 8 | 6090058 | 6429397 | . |
| 8 | 9880958 | 10479941 | . |  | 7 | 24010103 | 24835905 | . |  | 9 | 4044622 | 4347038 | . |  | 8 | 7190730 | 7625064 | . |
| 8 | 17430543 | 17647934 | . |  | 8 | 21179865 | 22013833 | . |  | 9 | 8810169 | 9059939 | . |  | 8 | 9781089 | 12568668 | . |
| 8 | 18010382 | 18289928 | 16_FP |  | 9 | 611511 | 2739949 | pQTL_22_F_RL |  | 10 | 5950430 | 6239713 | . |  | 8 | 12710190 | 13167344 | . |
| 8 | 18468764 | 19309846 | pQTL_21_F_PBintL |  | 9 | 2876799 | 3969917 | . |  | 10 | 16340269 | 16499989 | . |  | 8 | 13330627 | 14378167 | . |
| 8 | 20776098 | 20996272 | . |  | 9 | 8020913 | 8159764 | . |  | 11 | 10010125 | 10669797 | . |  | 8 | 15255553 | 15689928 | Rq26_F_DEPTH |
| 8 | 21540079 | 21878198 | . |  | 9 | 14910920 | 15397189 | . |  | 11 | 12740856 | 12969453 | . |  | 9 | 740091 | 1644785 | pQTL_22_F_RL |
| 9 | 3862228 | 4249235 | . |  | 10 | 13480427 | 13814148 | . |  | 11 | 13240076 | 15019698 | . |  | 9 | 14953138 | 15397189 | . |
| 11 | 4392040 | 4679985 | . |  | 11 | 16300552 | 18216478 | Rq13_J_TIL,Rq29_J_DEPTH,Rq30_J_DEPTH,Rq46_F_NCR,Rq63_J_THK |  | 11 | 15120111 | 15239075 | . |  | 12 | 7590964 | 8387853 | Rq32_F_DEPTH |
| 11 | 11210032 | 12119088 | . |  | 12 | 11491382 | 11859638 | . |  | 11 | 16250245 | 17389489 | Rq13_J_TIL,Rq46_F_NCR,Rq63_J_THK |  | 12 | 10220030 | 10532643 | . |
| 11 | 13730360 | 14339954 | . |  |  |  |  |  |  | 12 | 24630060 | 25059631 | . |  | 12 | 18290084 | 19168766 | . |
| 11 | 16291062 | 16918915 | Rq13_J_TIL,Rq46_F_NCR |  |  |  |  |  |  |  |  |  |  |  | 12 | 21791160 | 21908064 | . |
| 11 | 17080219 | 17429388 | Rq63_J_THK |  |  |  |  |  |  |  |  |  |  |  |  |  |  |  |
| 12 | 19790532 | 20088682 | . |  |  |  |  |  |  |  |  |  |  |  |  |  |  |  |
| 12 | 24680110 | 25069171 | . |  |  |  |  |  |  |  |  |  |  |  |  |  |  |  |
