## Supplementary material for "Evidence of selection, adaptation and untapped diversity in Vietnamese rice landraces": Table S14. Overlap of the selected regions in Indica subpopulations with QTL found in Vietnamese rice datasets

| **Table S14. Overlap of the selected regions in Indica subpopulations with QTL found in Vietnamese rice datasets.** | | | | | | | | | |  |  |  |  |  |  |  |  |  |  |  |  |  |  |
| --- | --- | --- | --- | --- | --- | --- | --- | --- | --- | --- | --- | --- | --- | --- | --- | --- | --- | --- | --- | --- | --- | --- | --- |
| Indica I1 | | | |  | Indica I2 | | | |  | Indica I3 |  |  |  |  | Indica I4 |  |  |  |  | Indica I5 |  |  |  |
| chrom | start | end | QTL |  | chrom | start | end | QTL |  | chrom | start | end | QTL |  | chrom | start | end | QTL |  | chrom | start | end | QTL |
| 1 | 4150316 | 4499460 | Rq5_F_TIL |  | 1 | 1602005 | 1708849 | qTTW1_I |  | 1 | 4847914 | 5209671 | . |  | 1 | 3580136 | 3919849 | . |  | 1 | 5563164 | 6569946 | Rq73_F_I_DWB60_DRW,Rq84_F_I_DRP,pQTL_2_F_SpN |
| 1 | 21613959 | 21979848 | . |  | 1 | 1870949 | 2334664 | Rq17_F_J_DEPTH,Rq83_F_J_DRP |  | 1 | 40530842 | 41049609 | L3_I_FW_TW,qSHL2_F |  | 1 | 5120479 | 6029761 | Rq73_F_I_DWB60_DRW,pQTL_2_F_SpN |  | 1 | 12270588 | 12957024 | . |
| 1 | 31472671 | 32239937 | . |  | 1 | 5080269 | 5739672 | pQTL_2_F_SpN |  | 2 | 18510227 | 18939401 | . |  | 1 | 6160663 | 7198137 | Rq84_F_I_DRP,pQTL_3_F_PBintL |  | 1 | 17910736 | 18069653 | . |
| 1 | 34860353 | 35207478 | . |  | 1 | 6300803 | 7979979 | Rq84_F_I_DRP,pQTL_3_F_PBintL |  | 2 | 22700005 | 22957766 | . |  | 2 | 2111412 | 3009215 | . |  | 1 | 21880287 | 22319111 | . |
| 1 | 37960982 | 38789946 | L2_I_FW_TW,qSHW1_qTTW3_I |  | 1 | 27630084 | 29559603 | Rq36_J_NCR |  | 2 | 28690981 | 28939777 | qRTL1_F_I |  | 2 | 5070311 | 5371041 | . |  | 1 | 37850965 | 38378420 | L2_I_FW_TW |
| 1 | 39050584 | 39289403 | Rq19_F_DEPTH |  | 1 | 32170424 | 32349385 | . |  | 2 | 33090089 | 33229942 | . |  | 2 | 6661353 | 8006638 | 1_DI |  | 1 | 40530842 | 40679473 | qSHL2_F |
| 1 | 39752942 | 40246023 | . |  | 1 | 40650110 | 40779365 | qSHL2_F |  | 2 | 35356767 | 35888657 | 3_GL_jap,Rq57_I_THK,Rq58_F_J_THK |  | 2 | 22820187 | 23059109 | . |  | 1 | 41340158 | 41769828 | . |
| 2 | 1140934 | 1497939 | . |  | 1 | 42210570 | 42479273 | . |  | 3 | 18647 | 198375 | . |  | 3 | 5870918 | 6488416 | . |  | 2 | 677 | 354653 | . |
| 2 | 30050002 | 30355131 | . |  | 2 | 6661686 | 7139620 | 1_DI |  | 3 | 15501782 | 15657428 | . |  | 3 | 28760342 | 29767679 | . |  | 2 | 1020920 | 1369616 | . |
| 2 | 32860707 | 33479992 | . |  | 2 | 26570785 | 27539831 | . |  | 3 | 22220103 | 22469108 | . |  | 3 | 31420261 | 31979360 | . |  | 2 | 2050537 | 2469973 | . |
| 3 | 18647 | 229810 | . |  | 2 | 30920522 | 31289143 | . |  | 3 | 23481393 | 23739981 | . |  | 3 | 33860286 | 33988653 | . |  | 2 | 3101251 | 3324479 | . |
| 3 | 7660454 | 8009419 | . |  | 2 | 35180287 | 35609667 | Rq57_I_THK,Rq58_F_J_THK |  | 3 | 24385013 | 25119339 | . |  | 3 | 35190028 | 35409927 | qTTW4_I |  | 2 | 3765987 | 4489973 | . |
| 3 | 24470025 | 24738979 | . |  | 3 | 1576126 | 1719972 | . |  | 3 | 26223613 | 27717921 | Rq8_F_TIL_NCR |  | 3 | 35801311 | 36199606 | Rq59_I_THK |  | 2 | 7320174 | 7989585 | . |
| 4 | 2160005 | 2459528 | . |  | 3 | 11280376 | 11909531 | . |  | 3 | 28122515 | 28448296 | . |  | 4 | 16020012 | 16499177 | 8_HD |  | 2 | 20981182 | 21185348 | . |
| 4 | 23300503 | 25268402 | pQTL_15_I_RL |  | 3 | 27282759 | 27569861 | . |  | 3 | 33230031 | 33487345 | . |  | 4 | 19160012 | 19722273 | . |  | 2 | 24920001 | 25369666 | . |
| 4 | 28010009 | 29648043 | . |  | 3 | 28120306 | 28455896 | . |  | 4 | 13115228 | 13278339 | Rq80_F_SRP |  | 4 | 19871085 | 20368434 | . |  | 2 | 28191142 | 29329745 | qRTL1_F_I |
| 4 | 30851269 | 31059987 | . |  | 4 | 23420010 | 23848589 | . |  | 4 | 14020352 | 14299893 | . |  | 4 | 23950209 | 25289937 | pQTL_15_I_RL |  | 3 | 6691656 | 8177456 | . |
| 5 | 9571530 | 9909669 | . |  | 4 | 34670218 | 35129844 | . |  | 4 | 14470522 | 14829897 | . |  | 5 | 3560148 | 3769018 | . |  | 3 | 12650311 | 12946658 | . |
| 6 | 3911289 | 4169389 | Rq67_F_I_DW2040,Rq69_F_I_DW4060_DRW_RDW |  | 5 | 22080192 | 22569232 | . |  | 4 | 22150905 | 22972874 | . |  | 5 | 17390917 | 18297730 | Rq34_F_MRL |  | 3 | 15380808 | 15879517 | . |
| 6 | 6540668 | 6728210 | 12_GL |  | 6 | 6150070 | 6679742 | . |  | 4 | 23113340 | 23839932 | . |  | 5 | 23260571 | 23469490 | . |  | 3 | 20960124 | 21669162 | . |
| 6 | 7070142 | 7259895 | 12_GL |  | 6 | 7420340 | 7629449 | 13_GL |  | 4 | 24730034 | 25219983 | . |  | 5 | 25260383 | 25908908 | . |  | 3 | 24370632 | 24999498 | . |
| 6 | 11223260 | 12409952 | . |  | 6 | 23890037 | 24279585 | . |  | 4 | 28740828 | 29429404 | . |  | 6 | 5670161 | 6879691 | 12_GL |  | 3 | 25193549 | 25587517 | . |
| 6 | 19340269 | 19799359 | . |  | 6 | 25100965 | 25629859 | . |  | 6 | 7860166 | 8399897 | Rq2_J_LLGHT |  | 6 | 7771233 | 7909949 | Rq2_J_LLGHT |  | 3 | 27910148 | 29199870 | . |
| 6 | 21010770 | 21759494 | Rq82_I_SRP_DRP |  | 6 | 27000229 | 27159758 | . |  | 6 | 24230987 | 24729326 | Rq88_F_J_R-S |  | 6 | 13070070 | 13496098 | . |  | 3 | 29431523 | 29589724 | . |
| 6 | 21991432 | 22199212 | . |  | 7 | 2560765 | 2689832 | . |  | 6 | 25110334 | 25669789 | . |  | 6 | 20897950 | 21399021 | . |  | 3 | 33372038 | 33639859 | . |
| 6 | 27410550 | 27859734 | . |  | 7 | 6660098 | 7369144 | . |  | 7 | 630899 | 1318272 | . |  | 8 | 2271137 | 2589970 | . |  | 4 | 62390 | 489186 | . |
| 7 | 3107321 | 3409772 | . |  | 7 | 8860389 | 9643034 | . |  | 7 | 1750216 | 2289476 | . |  | 8 | 2740035 | 3188493 | . |  | 4 | 5251107 | 5436839 | . |
| 7 | 6040539 | 6387372 | . |  | 7 | 13353887 | 13668125 | . |  | 7 | 26260178 | 26769981 | . |  | 8 | 3380100 | 4009807 | Tq10_F_RWC_T3,qSHL4_F |  | 4 | 33073892 | 33369648 | . |
| 7 | 18991429 | 19489875 | pQTL_18_F_PBintL |  | 7 | 23250466 | 24858300 | . |  | 7 | 27780086 | 28269536 | . |  | 8 | 4862084 | 5479199 | pQTL_19_F_J_PBN_PBL |  | 4 | 34813879 | 35098724 | . |
| 7 | 19601007 | 20018912 | Tq7_F_RWC_S3 |  | 7 | 27710685 | 28039679 | . |  | 7 | 28674534 | 29429594 | . |  | 8 | 8842224 | 9459456 | . |  | 5 | 386347 | 1563159 | 9_PL |
| 7 | 20530677 | 20989643 | Rq12_F_I_TIL,Tq8_F_I_RWC_T2_Score1 |  | 9 | 1190635 | 2069922 | qTTW7_J |  | 8 | 2298155 | 2568547 | . |  | 8 | 14930903 | 15776544 | Rq26_F_DEPTH |  | 6 | 6640258 | 7189250 | 12_GL |
| 7 | 26510388 | 27018603 | . |  | 9 | 13750284 | 14108832 | 19_HD |  | 8 | 3320606 | 3667332 | qSHL4_F |  | 9 | 1130128 | 1879894 | qTTW7_J |  | 6 | 7860166 | 8418475 | Rq2_J_LLGHT |
| 8 | 1061532 | 1256342 | . |  | 9 | 15998759 | 18308374 | . |  | 8 | 5206699 | 5549978 | pQTL_19_F_J_PBN_PBL |  | 9 | 7270061 | 8029789 | 18_FP |  | 6 | 19470641 | 20499968 | 14_PL,Rq35_F_J_MRL_NCR |
| 8 | 1500033 | 1759838 | . |  | 9 | 18460288 | 19228937 | . |  | 8 | 5681438 | 5875735 | . |  | 9 | 12290300 | 13058911 | . |  | 7 | 19443608 | 19825988 | Tq7_F_RWC_S3 |
| 8 | 19631318 | 19989977 | . |  | 9 | 19361714 | 19589466 | . |  | 9 | 16250715 | 17042053 | . |  | 9 | 18417227 | 18956920 | . |  | 7 | 29030233 | 29677525 | Rq25_F_DEPTH |
| 8 | 27840071 | 28024852 | . |  | 10 | 2360045 | 2509134 | . |  | 9 | 17520205 | 18729927 | . |  | 10 | 19000712 | 19345847 | . |  | 8 | 3484045 | 3758632 | qSHL4_F |
| 10 | 21720070 | 22119997 | . |  | 11 | 2094852 | 2247719 | . |  | 10 | 16460849 | 16869847 | . |  | 11 | 1970277 | 3142910 | Tq12_F_I_RWC_T2_RWC_T3_RWC_S2_Score3 |  | 8 | 5052017 | 5809093 | pQTL_19_F_J_PBN_PBL |
| 11 | 2440310 | 2808594 | Tq12_F_I_RWC_T2_RWC_T3_RWC_S2_Score3 |  | 12 | 5550062 | 5979273 | pQTL_28_F_SBintL |  | 10 | 18000537 | 18469985 | L11_I_FW_TW_DW,pQTL_24_F_PBN |  | 11 | 19951867 | 20549979 | Tq17_F_I_Score4,pQTL_25_F_J_RL,qSHL5_J |  | 8 | 19431460 | 20459346 | . |
| 11 | 6921014 | 7288214 | Tq14_F_I_RWC_T1_RWC_S1 |  | 12 | 6657064 | 6989784 | L12_F_FW_TW |  | 10 | 19480156 | 19763968 | . |  |  |  |  |  |  | 8 | 24300313 | 24859863 | . |
| 11 | 23170362 | 23507750 | . |  | 12 | 24090147 | 24537223 | qSHW7_qTTW8_I |  | 10 | 19901715 | 20063176 | . |  |  |  |  |  |  | 9 | 14820651 | 15259615 | . |
| 11 | 27381691 | 27627281 | . |  | 12 | 26430898 | 26838283 | L13_F_FW_TW |  | 11 | 19930315 | 20819799 | Tq17_F_I_Score4,pQTL_25_F_J_RL,qSHL5_J |  |  |  |  |  |  | 9 | 16430191 | 18049085 | . |
| 11 | 28280726 | 28518974 | . |  |  |  |  |  |  | 12 | 26170080 | 26459232 | . |  |  |  |  |  |  | 9 | 18292494 | 18798654 | . |
| 12 | 1050059 | 1582315 | Rq47_F_I_NCR,Rq71_F_I_DW4060 |  |  |  |  |  |  |  |  |  |  |  |  |  |  |  |  | 9 | 19710325 | 20229472 | . |
| 12 | 2542327 | 2839661 | qSHW6_ qSHL6_F |  |  |  |  |  |  |  |  |  |  |  |  |  |  |  |  | 10 | 5381471 | 5869967 | . |
|  |  |  |  |  |  |  |  |  |  |  |  |  |  |  |  |  |  |  |  | 10 | 10528884 | 11139609 | . |
|  |  |  |  |  |  |  |  |  |  |  |  |  |  |  |  |  |  |  |  | 10 | 11991467 | 12409929 | . |
|  |  |  |  |  |  |  |  |  |  |  |  |  |  |  |  |  |  |  |  | 10 | 18732199 | 19209687 | . |
|  |  |  |  |  |  |  |  |  |  |  |  |  |  |  |  |  |  |  |  | 11 | 2510079 | 3239747 | Tq12_F_I_RWC_T2_RWC_T3_RWC_S2_Score3 |
|  |  |  |  |  |  |  |  |  |  |  |  |  |  |  |  |  |  |  |  | 11 | 4590276 | 5937318 | Rq43_F_I_NCR |
|  |  |  |  |  |  |  |  |  |  |  |  |  |  |  |  |  |  |  |  | 11 | 6060058 | 6179872 | . |
|  |  |  |  |  |  |  |  |  |  |  |  |  |  |  |  |  |  |  |  | 12 | 50720 | 659181 | . |
|  |  |  |  |  |  |  |  |  |  |  |  |  |  |  |  |  |  |  |  | 12 | 25861119 | 26518838 | . |
